# Mechanisms of mRNA processing defects in inherited *THOC6* intellectual disability syndrome

**DOI:** 10.1101/2022.09.06.506502

**Authors:** Elizabeth Werren, Geneva R. LaForce, Anshika Srivastava, Delia R. Perillo, Katherine Johnson, Safa Baris, Brandon Berger, Samantha L. Regan, Christian D. Pfennig, Sonja de Munnik, Rolph Pfundt, Malavika Hebbar, Raúl Jimenez-Heredia, Elif Karakoc-Aydiner, Ahmet Ozen, Jasmin Dmytrus, Ana Krolo, Ken Corning, EJ Prijoles, Raymond J. Louie, Robert R. Lebel, Thuy-Linh Le, Chris Gordon, Kaan Boztug, Katta M. Girish, Anju Shukla, Stephanie L. Bielas, Ashleigh E. Schaffer

## Abstract

*THOC6* is the genetic basis of autosomal recessive *THOC6* Intellectual Disability Syndrome (TIDS). THOC6 facilitates the formation of the Transcription Export complex (TREX) tetramer, composed of four THO monomers. The TREX tetramer supports mammalian mRNA processing that is distinct from yeast TREX dimer functions. Human and mouse TIDS model systems allow novel THOC6-dependent TREX tetramer functions to be investigated. Biallelic loss-of-function (LOF) *THOC6* variants do not influence the expression and localization of TREX members in human cells, but our data suggests reduced binding affinity of ALYREF. Impairment of TREX nuclear export functions were not detected in cells with biallelic *THOC6* LOF. Instead, mRNA mis-splicing was observed in human and mouse neural tissue, revealing novel insights into THOC6-mediated TREX coordination of mRNA processing. We demonstrate that THOC6 is required for regulation of key signaling pathways in human corticogenesis that dictate the transition from proliferative to neurogenic divisions that may inform TIDS neuropathology.

## INTRODUCTION

Intellectual disability (ID) is a clinical feature of neurodevelopmental disorders characterized by limitations in cognitive ability and adaptive behavior (Schalock et al., 2021). In recent years, genetic etiologies of syndromic ID have become increasingly heterogeneous due to broad use of exome-based genetic testing (Anazi et al., 2017; Gieldon et al., 2018; Retterer et al., 2016; Vasudevan and Suri, 2017; Yang et al., 2014). Monogenetic causes account for a substantial portion of syndromic ID, with many following an autosomal-recessive mode of inheritance (Kochinke et al., 2016). ID-associated genes are enriched in diverse biological networks including metabolism, nervous system development, RNA metabolism, transcription, sonic hedgehog signaling, glutamate signaling, peroxisomes, glycosylation, and cilia (Kochinke *et al.*, 2016). THOC6 Intellectual Disability Syndrome (TIDS; OMIM# 613680) is one such recessive disorder, attributed to biallelic pathogenic variants in *THOC6* (Accogli et al., 2018; Amos et al., 2017; Anazi et al., 2016; Boycott et al., 2010; Casey et al., 2016; Gupta et al., 2020; Hassanvand Amouzadeh et al., 2020; Kiraz et al., 2022; Mattioli et al., 2018; Ruaud et al., 2022; Zhang et al., 2020). Individuals with TIDS exhibit moderate to severe syndromic ID with microcephaly and multi-organ involvement. Genetic testing is necessary for diagnosis of TIDS, as clinical features overlap other inherited neurodevelopmental disorders (Lemire et al., 2020).

THOC6 is a subunit of the six member THO (suppressors of the transcriptional defects of hpr1 delta by overexpression) complex (Jimeno and Aguilera, 2010). The THO complex is a core component of the transcription/export (TREX) complex. Prior to translation, mRNA transcripts progress through a series of coordinated steps that include mRNA 5’ capping, splicing, and 3’ end processing to create a messenger ribonucleoprotein complex (mRNP) capable of translocating through the nuclear pore complex into the cytoplasm (Heath et al., 2016; Köhler and Hurt, 2007; Xie and Ren, 2019). TREX is necessary for proper mRNA processing and nuclear export required for gene expression, with a more appreciated role in export based on current literature (Chi et al., 2013; Heath *et al.*, 2016; Luna et al., 2012; Masuda et al., 2005; Peña et al., 2012; Rondón et al., 2010; Wickramasinghe and Laskey, 2015). TREX is recruited to the maturing mRNP during transcription (Heath *et al.*, 2016; Viphakone et al., 2019), in coordination with pre-mRNA binding proteins at the cap-binding complex (CBC), exon-junction complex (EJC), and the polyadenylated 3’ end (Cheng et al., 2006; Gromadzka et al., 2016; Merz et al., 2007; Shi et al., 2017). Despite a well-studied role in mRNA export, the precise timing of initial TREX recruitment to maturing mRNPs in mammalian cells is unclear, but mounting evidence suggests mammalian TREX associates to the mRNPs during splicing (Chi *et al.*, 2013; Luo et al., 2001; Masuda *et al.*, 2005). Ultimately, TREX promotes the loading of licensing export factors required for nuclear pore docking and export (Hautbergue et al., 2008; Köhler and Hurt, 2007; Strässer and Hurt, 2001; Taniguchi and Ohno, 2008).

While the general role of TREX in mRNA processing and export is thought to be conserved, there are notable species differences in TREX composition and function that mirror the evolutionary complexity of mRNA processing requirements. In yeast, the TREX dimer is composed of two five-subunit THO monomers, with THOC6 being the notable exception. Yeast TREX monomers dimerize via the coiled coil domains of Thp2 and Mft1, the yeast orthologs of ­­­­­­­­­­­THOC5 and THOC7 (Pühringer et al., 2020). In humans, the THO monomers are composed of six subunits, including THOC6, that form a tetramer with THOC6 serving as the central tethering component (Pühringer *et al.*, 2020). The increased size and molecular complexity of the mammalian TREX tetramer correlates with increased mRNA processing demands that have evolved in organisms with higher transcriptome complexity and mRNP composition, namely expression of long genes with high levels of complex splicing patterns (Singh et al., 2015). For example, introns comprise ∼24% of mammalian genomes (Venter et al., 2001). On the level of gene organization, introns are longer and are present in >95% of all human genes (Chen et al., 2006; Lander et al., 2001; Nagasaki et al., 2005). By contrast, introns constitute only 5% of yeast genes, are short relative to the gene length, and mostly limited to one per gene (Chen *et al.*, 2006; Chervitz et al., 1999; Juneau et al., 2006; Nagasaki *et al.*, 2005). Thus, alternative splicing is rare in yeast, but plays a major role in gene expression in mammals, especially humans. Functional differences between a TREX dimer and tetramer may correlate to differences in complex size and number of molecules that can be simultaneously accrued to coordinate mRNP processing through export (Pühringer *et al.*, 2020).

Yeast TREX dimer are configured to brings two bound Sub2 helicases (yeast ortholog to the metazoan DDX39B/UAP56 RNA helicase which couples ATP hydrolysis to mRNA release) in proximity. This formation enables Yra1 (yeast ortholog of ALYREF), a mRNP processing/export factor to bridge the THO monomers by binding the N- and C-terminal of aligned Sub2 components. Components of the mammalian TREX tetramer exhibit the ability to interact with a diversity of mRNP processing and export factors. The TREX tetramer also binds UAP56 molecules, however the tetramer organization means the putative loading sites for ALYREF are doubled, the potential of affording increased affinity of processing adaptors to their target mRNPs (Pühringer *et al.*, 2020). The tetramer structure is predicted to enhance physical support for processing longer mammalian transcripts, allowing TREX to serve as a mRNA chaperone to prevent formation of DNA-RNA hybrid or R-loop structures that can promote genome instability (Luna et al., 2019; Pérez-Calero et al., 2020). In line with the mammalian mRNP processing complexity, the tetramer recruits a portfolio of functionally diverse auxiliary factors, predicted to enhance coordination of mRNA processing steps. Several TREX adaptor proteins in metazoans that have no yeast ortholog include, DDX39A, CHTOP, UIF, LUZP4, POLDIP3, ZC3H11A, ERH, ZC3H18, SRRT, and NCBP3, representing proteins that participate in each step of mRNP processing and export and hinting at a level of coordination required as the complexity of mRNP processing evolved with the emergence of longer transcripts with elevated splicing (Dufu et al., 2010; Heath *et al.*, 2016).

While functions of THO within TREX have primarily focused on mRNP export (Chi *et al.*, 2013; Guria et al., 2011; Maeder et al., 2018), disruption of the tetramer is predicted to disrupt coordination of mRNP processing steps that precede export. TREX-associated functions of CHTOP and ALYREF exemplify this molecular biology. CHTOP and ALYREF bind to THO-bound UAP56 in a mutually exclusive manner (Chang et al., 2013). ALYREF exhibits preferential binding to the 5’ end and 5’ splice sites of mRNA to regulate splicing fidelity, whereas CHTOP preferentially binds the 3’ UTR to regulate polyadenylation site choice along with THOC5 (Viphakone *et al.*, 2019). Lastly, ALYREF, CHTOP, and THOC5 interact with and load the global NXF1-NXT1 licensing heterodimer onto mRNA for export (Hautbergue *et al.*, 2008; Köhler and Hurt, 2007; Strässer and Hurt, 2001; Taniguchi and Ohno, 2008). This organization provides structural logic for how the molecular component required for the many aspects of mRNP processing can be coordinated, but the impact of disrupting a single step on progression of the entire process has not been evaluated.

The conservation of THO dimer functions in mammalian cells is an open question. THOC1, THOC3, THOC5, and THOC7 exhibit high probability of loss-of-function intolerance (pLI) in gnomAD and have not been identified as the genetic basis of developmental disorders, suggesting conserved THO components are likely embryonic lethal. *THOC2* is the genetic basis of an X-linked neurodevelopmental disorder (Kumar et al., 2015). Depletion of THO components THOC1-THOC5 and THOC7 lead to strong nuclear export defects (Chi *et al.*, 2013). This leads to the speculation that dimers retain mRNP functions in mammals, and that THOC6-dependent tetramer functions enhance the efficiency and coordination of these activities. This would also explain the tissue sensitivity in TIDS, where development is disrupted in tissues that disproportionately express long genes. In addition, neural-expressed genes display many isoforms which enhance the mRNP processing burden.

THOC6 evolved as a scaffolding protein to create a larger TREX complex, that has implication for mRNA processing coordination in metazoans relative to yeast. Despite extensive research on TREX, there is a lack of research in neural cells, cells that undergo extensive changes in mRNA processing during differentiation, particularly alternative splicing with increased intron retention (Mauger et al., 2016). Given this, there may be functions of THO within a tetramer conformation mediated by THOC6 that have not been the main focus of study. Here, we investigated this by utilizing a series of models with pathogenic alleles that disrupt THOC6 to assess essential functions of tetramers in features of mRNA processing in neural development. First, we contribute to the *THOC6* pathogenic allele series and TIDS clinical phenotypic spectrum, extending the total number of reported *THOC6* variants in TIDS to 20 and the total number of reported affected individuals to 34. Second, we generated a *Thoc6* mouse model and human induced pluripotent stem cell (iPSC)-derived cell culture models to investigate shared pathogenic mechanisms of mammalian *THOC6.* Given the high penetrance of microcephaly in TIDS, we focus on mRNA processing/export changes and accompanying phenotypes that occur during cortical development by analyzing primary mouse and dorsal forebrain fated human organoids. We propose a model of unproductive selective mRNA processing/export from partial TREX disruption due to loss-of-function *THOC6* alleles leading to dysregulation of proliferation and differentiation. Our findings reveal a broader supportive role across mRNA processing within the context of THOC6 variants than has previously been attributed to THO.

## RESULTS

### Biallelic missense and nonsense *THOC6* variants are the genetic basis of TIDS

While initially detected in Hutterite populations (Boycott *et al.*, 2010), a growing number of pathogenic biallelic *THOC6* variants are being discovered across the globe in individuals of diverse ancestry (Accogli *et al.*, 2018; Amos *et al.*, 2017; Anazi *et al.*, 2016; Beaulieu et al., 2013; Casey *et al.*, 2016; Gupta *et al.*, 2020; Hassanvand Amouzadeh *et al.*, 2020; Kiraz *et al.*, 2022; Mattioli *et al.*, 2018; Ruaud *et al.*, 2022; Zhang *et al.*, 2020). As part of an ongoing effort to determine the genetic etiologies of syndromic ID, we discovered nine *THOC6* variants by exome-based genetic testing (Figure 1A). We confirmed six recurrent variants at W100, G190, V234, and G275 amino acids and identified three novel alleles (p.Q47*, p.E188K, and p.R247Q). The clinical phenotype associated with *THOC6* variants affirm penetrance of the core clinical features of TIDS, namely global developmental delay, moderate to severe ID and facial dysmorphisms (Figure 1B). Variable expressivity of cardiac and renal malformations, structural brain abnormalities with and without seizures, urogenital defects, recurrent infections, and feeding complications were also noted, clinical features that highlight the multiorgan involvement of this developmental syndrome (Figure 1C). Detailed clinical summaries for all individuals are provided in Table S1 and S2.

**Figure 1.**
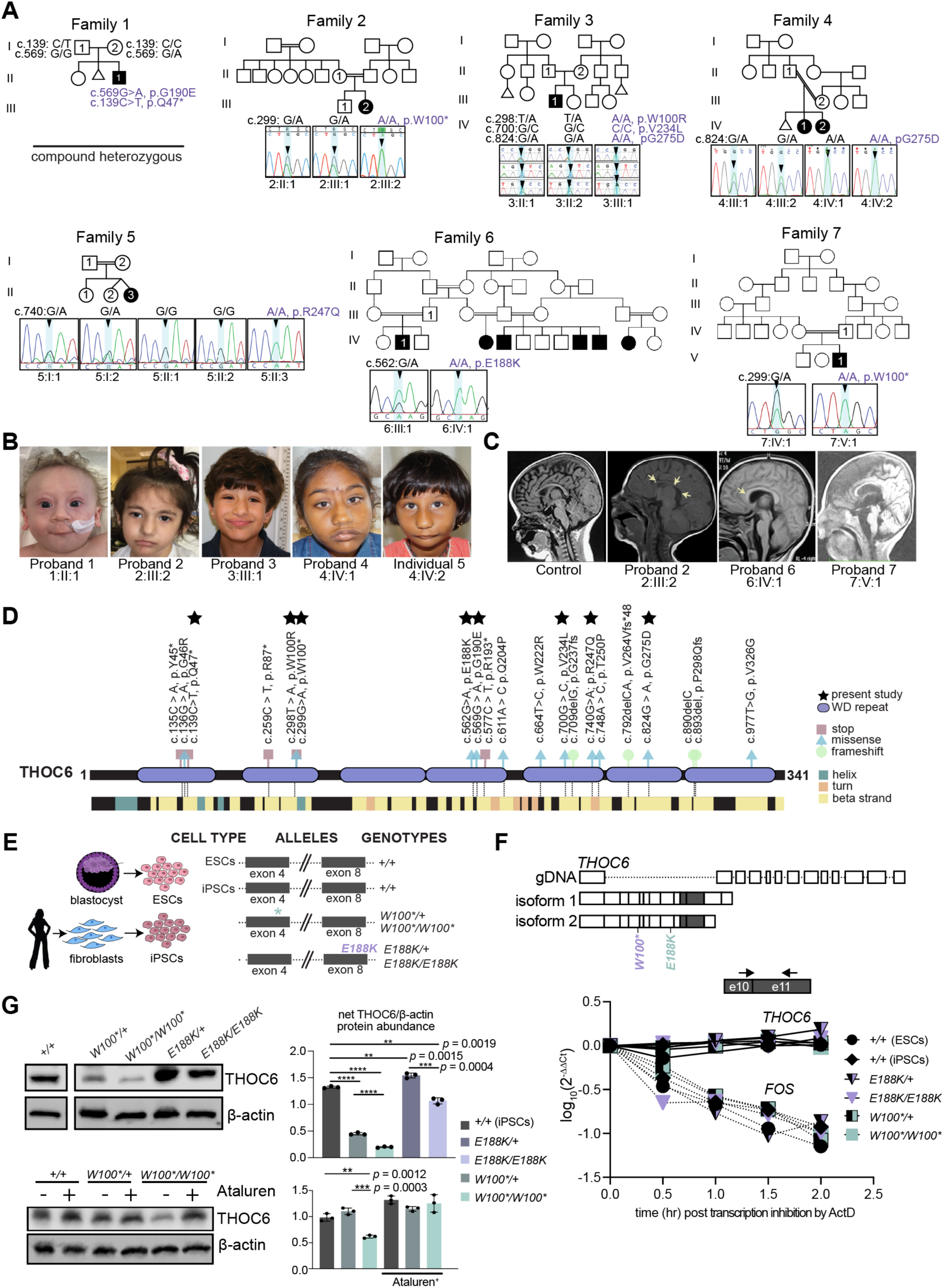
Biallelic pathogenic variants in *THOC6* cause syndromic intellectual disability. (A) Pedigree drawings of segregating TIDS phenotypes in families 1-7, with generations listed on the left-hand side. Females are represented as circles and males are denoted by squares. Miscarriages are denoted by small triangles. Affected family members are indicated by solid black coloring while unaffected are unfilled. Consanguineous partnerships are represented by double lines. Chromatograms from Sanger sequencing of THOC6 confirmation of genotypes are provided for each tested family member in families 1-7. (B) Facial photographs of Probands 1-4 and Individual 5. (C) Sagittal brain MRI showing corpus callosum dysgenesis (Probands 2, 6, & 7) and cortical and cerebellar atrophy (Proband 6) compared to control (left). (D) Canonical THOC6 protein map consisting of 341 amino acids. WD40 repeat domains 1-7 are denoted by purple rectangles. Location of known pathogenic variants are annotated relative to linear protein map (top) and secondary structure (below). Variants reported in present study are distinguished by a black star. Missense (blue triangle), nonsense (red square), and frameshift (green circle). (E) Schematic of patient and control-derived human cell types and respective genotypes. (F) Decay of *THOC6* mRNA (solid line) following ActD transcriptional inhibition compared to *FOS* mRNA decay (dotted line) in human ESC/iPSCs across genotypes. Values calculated relative to *GAPDH* reference mRNA. (G) Western blot of human ESC/iPSCs indicating reduced THOC6 protein expression in *THOC6^W100*/W100*^* iPSCs compared to unaffected controls. Confirmation of readthrough by ataluren treatment (30 μM). Abundance quantifications relative to β-actin control (right). Data represented as mean ±SEM. P-value, two-tailed unpaired *t* test. ****, p-value <0.0001.

The novel *THOC6* variants are representative of previously described nonsense and missense variants that contribute equally to the severity of TIDS phenotypes. We describe a nonsense *THOC6* c.139C>T, (p.Q47*) variant in exon 2 of proband 1, a missense c.740G>A, (p.R247Q) variant in exon 11 of proband 5, and a missense c.562G>A, (p.E188K) variant in proband 6 (Figures 1A and 1C). THOC6 is comprised of seven WD40 repeat domains (Figure 1D) that form a β-propeller structure when folded. These novel variants, like other clinically relevant *THOC6* variants, map to the WD40 repeats that comprise the beta strand structural regions of THOC6 (Figure 1D). The p.Q47* nonsense variant represents a cluster of three *THOC6* variants in the first WD40 repeat that exhibit a consistent genotype-phenotype correlation for both recessive nonsense and missense variants. The same trend is observed for the novel missense variants that are localized to subsequent THOC6 WD40 domains, which support a loss-of-function (LOF) pathogenic mechanism for both *THOC6* variant types.

A LOF mechanism is also implicated by the clinical consistency observed between biallelic inheritance of pathogenic *THOC6* haplotypes and biallelic inheritance of a single haplotype variant alone. Biallelic inheritance of a triple-variant haplotype (TVH), *THOC6* c.[298T>A;700G>C;824G>A], (p.[W100R;V234L;G275D]) has been reported in seven individuals with clinical features of TIDS (Casey *et al.*, 2016; Gupta *et al.*, 2020; Mattioli et al., 2019; Ruaud *et al.*, 2022). The TVH segregates as a founder haplotype in individuals of European ancestry (Mattioli et al., 2019). In comparison, a homozygous *THOC6* c.824G>A; p.G275D variant was identified in siblings with classic TIDS in family 4 who are of South Asian ancestry. This finding provides additional evidence for the pathogenicity of the TVH *THOC6* c.824G>A; p.G275D variant but does not negate the predicted pathogenicity of the corresponding W100R or V234L TVH variants. Pathogenicity of p.W100R or p.V234L are supported by variants detected in WD40 repeats 2 and 4 (Figure 1D). Comparing biallelic inheritance of TVH and *THOC6* c.824G>A; p.G275D suggests a single THOC6 variant in both alleles are sufficient to comprehensively disrupt THOC6, a baseline deficiency not exacerbated by accumulation of additional LOF variants.

### *THOC6* variants show mRNA stability with differential effect on protein abundance

To investigate the genetic mechanism of *THOC6*, the impact of on mRNA nonsense mediated decay (NMD) and protein expression on pathogenicity was tested in embryonic stem cells (ESCs) and iPSCs, collectively referred to as human pluripotent stem cells (hPSCs). hPSCs were reprogrammed from two individuals with TIDS (6:IV:1*, THOC6^E188K/E188K^* and 7:V:2, *THOC6^W100*/W100*^*) and their respective unaffected heterozygous parent (6:IV:2, *THOC6^E188K/+^* and 7:V:1, *THOC6^W100*/+^*) (Figure 1E), preserving the shared genetic background between affected and unaffected conditions. Consistent with a LOF mechanism, reduction in protein expression due to mRNA nonsense mediated decay was predicted. However, *THOC6* mRNA transcripts remain relatively stable between genotypes, as assessed by Actinomycin D treatment where transcription is inhibited, compared to the unstable mRNA, *FOS*, that is quickly degraded (Figure 1F) (Moon et al., 2012). This finding is consistent regardless of *THOC6* variant, with the c.299G>A, (p.W100*) and c.562G>A, (p.E188K) variants exhibiting similar decay rates as wildtype transcripts (Figure S1C and S1D). This finding could reflect defective NMD from failure of *THOC6-*affected mRNA to be exported to the cytoplasm. Nevertheless, the impact on protein expression is divergent between nonsense and missense variants. Significant reductions in THOC6 abundance were detected in *THOC6^W100*/W100*^* and *THOC6^E188K/E188K^* iPSCs relative to the *THOC6^+/+^* control, with the most remarkable reduction for *THOC6^W100*/W100*^*. Significant abundance differences were also noted for heterozygous unaffected iPSCs relative to wildtype controls (Figure 1G). Full-length THOC6 in *THOC6^W100*/W100*^* samples represent a minority readthrough product. Increased frequency of this rare event was promoted by treatment with 30 μM Ataluren, which extends translation by skipping premature termination codons, leading to an increase in THOC6 detectable by Western blot in *THOC6^W100*/W100*^* iPSCs (Figure 1G). No truncated product was observed by Western blotting in *THOC6^W100*/W100*^* and *THOC6^W100*/+^* iPSCs, suggesting THOC6 reduction is due to rapid degradation of an unstable, truncated protein. Stable expression from missense *THOC6* alleles suggests variant THOC6 is likely functionally inactive.

### *THOC6* variants interfere with TREX functions

Based on the solved crystal structure (Pühringer *et al.*, 2020), LOF *THOC6* variants are predicted to impair TREX tetramer formation. The WD40 repeat domains of THOC6 form beta strands predicted to provide the structural interface for TREX core tetramer formation. Pathogenic *THOC6* variants disrupt residues conserved in mammals, though several are not conserved across other metazoan species, mirroring TREX composition variability across species and the evolving function for THOC6 in TREX (Figure 2A). Evaluation of the TREX crystal structure indicates that pathogenic THOC6 variants are positioned at the TREX core tetrameric interface, where THOC5-THOC7 interaction is responsible for dimerization and THOC6-THOC5 interaction tetramerizes the complex (Figure 2B, 2C, and 2D). The allelic series of *THOC6* LOF variants implicate a pathogenic mechanism where LOF missense variants are predicted to perturb THOC6 β-propeller folding and/or interactions with THOC5 and THOC7 that are required for TREX tetramer assembly, and nonsense variants would produce a similar outcome due to low protein abundance (Pühringer *et al.*, 2020). Likewise, *THOC6* variants do not alter the protein abundance of other THO/TREX members (Figure 2E). Consistent with normal abundance of functional THO protein, subcellular localizations at nuclear speckle domain were observed by immunohistochemistry (Figures S1E and S1F). Conversely, ALYREF association with the THO subcomplex is diminished in *THOC6^E188K/E188K^* and *THOC6^W100*/W100*^* as assessed by co-immunoprecipitation with THOC5 and THOC6 in patient-derived iPSC lines (Figure 2F). These findings suggest a THOC6-dependent association of ALYREF to THO, with implications for the affinity of other adaptors due to the potential disruption of TREX tetramer formation.

**Figure 2.**
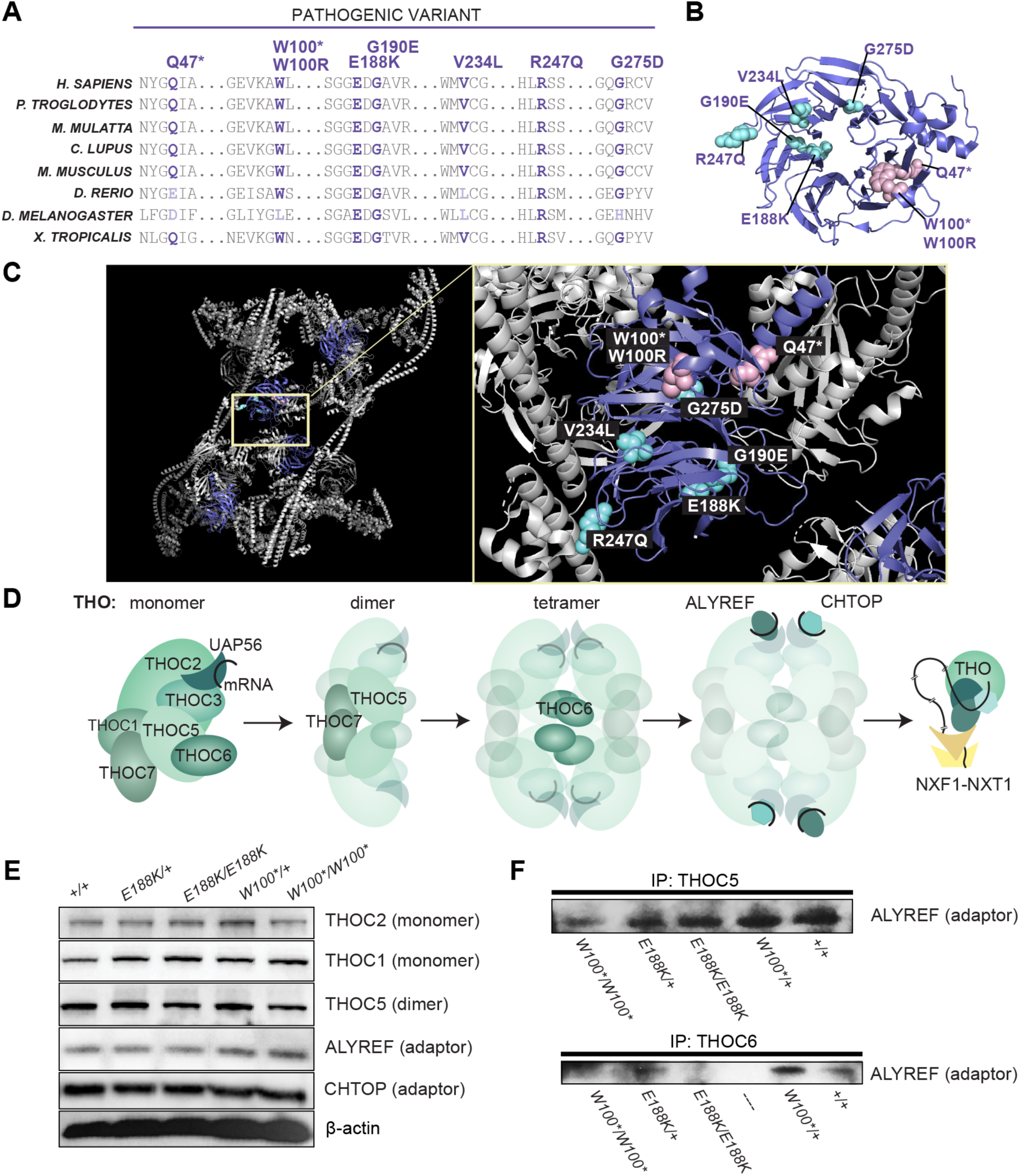
Genetic mechanism of biallelic pathogenic *THOC6* variants. (A) Amino acid alignment showing conservation of affected residues for pathogenic variants in present clinical study. Variants mapped to THOC6 folded β-propeller structure (B) and THO/TREX complex (C). (D) Schematic of TREX core tetrameric assembly mediated by THOC6 with functional implications for mRNA processing and export based on published crystal structure (Pühringer *et al.*, 2020). (E) Steady-state protein abundance for THO/TREX complex members and (F) ALYREF abundance following co-immunoprecipitation with THOC5 and THOC6 across genotypes.

### *Thoc6* is required for mouse embryogenesis

To investigate *Thoc6* pathogenic mechanisms in mammalian neural development *in vivo*, we used CRISPR/Cas9 genome editing to introduce an insertion variant in mouse *Thoc6* exon 1 that resulted in a premature termination codon predicted to ablate Thoc6 expression (p.P6Lfs*8, herein referred to as *Thoc6^fs^*) (Figure 3A). *Thoc6^+/fs^* male and female mice do not display phenotypic abnormalities, but their intercrosses yielded no homozygous offspring. Analysis at selected embryonic days (E) of gestation confirmed *Thoc6^fs/fs^* littermates die *in utero*. Differences in embryonic morphology between wildtype (WT) and *Thoc6^fs/fs^* embryos were noted starting at E7.5, the earliest day of analysis. By E9.5, *Thoc6^fs/fs^* embryos were smaller with delayed development; however, the difference in the developing neocortex was particularly pronounced (Figure 3B). No body turning differences were observed. Consistent with embryonic lethality, THOC6 was undetectable in E8.5 *Thoc6^fs/fs^* mouse embryos relative to control littermates (Figure 3C). In E9.5 *Thoc6^fs/fs^* embryos, a developmental timepoint with high Thoc6 expression, Thoc6 is detectable by Western blot at greatly diminished levels (Figure 3C). Embryonic lethality was confirmed by E11.5, indicating one functional allele of *Thoc6* is essential for mouse embryonic development (Figures 3C and 3D).

**Figure 3.**
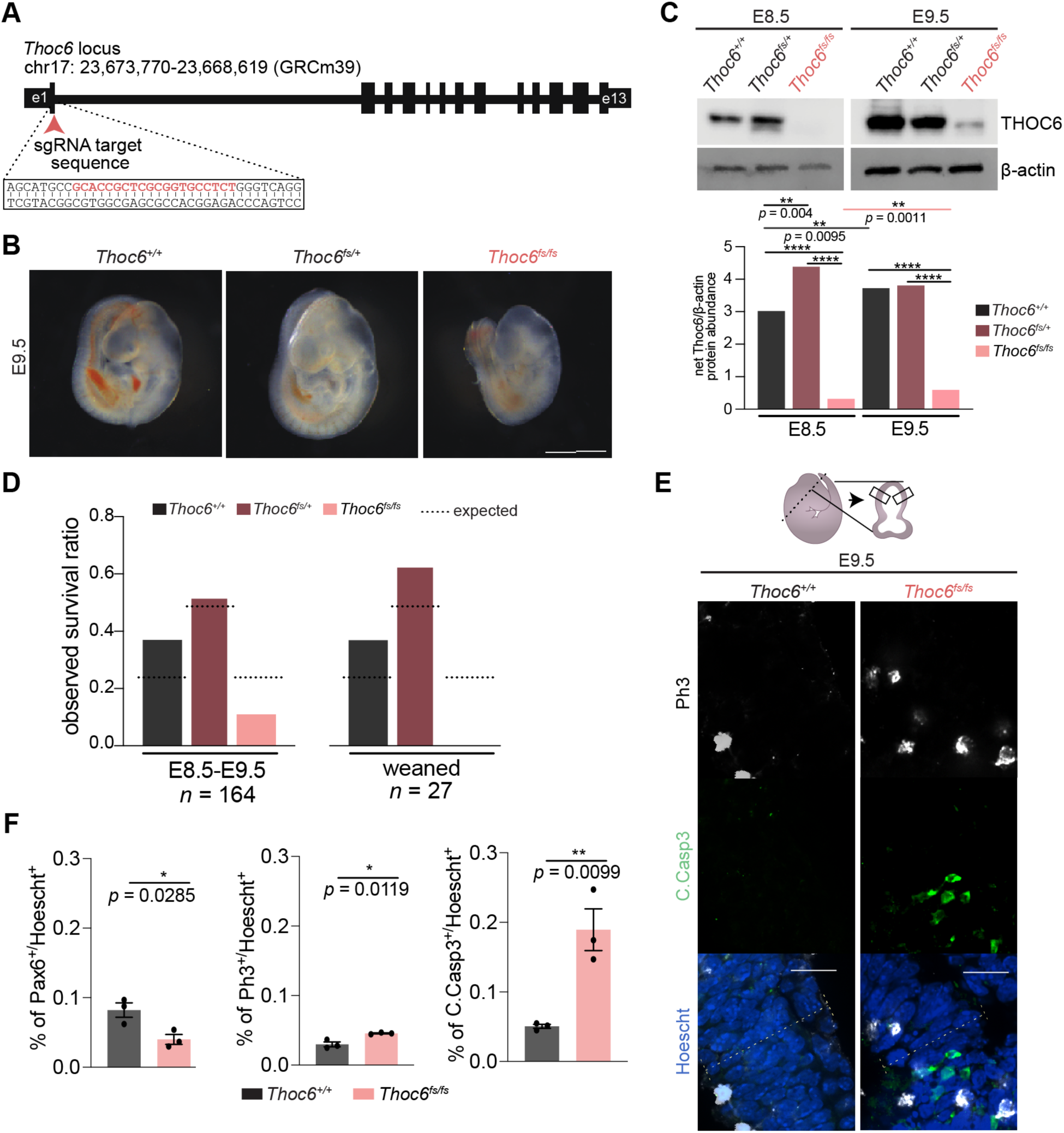
Generation of *Thoc6^fs/fs^* mouse model. (A) CRISPR/Cas9 editing strategy to introduce frameshift variants in mouse *Thoc6*. (B) Representative images of isolated *Thoc6^+/+^*, *Thoc6^fs/+^*, and *Thoc6^fs/fs^* whole embryos at E9.5 prior. Scale bar: 50 μm. (C) Western blot analysis with quantifications of E8.5 and E9.5 mouse embryos showing increased expression during development. Ablation of Thoc6 protein in *Thoc6^fs/fs^* is observed at E8.5, with presence of band suggesting read-through product at E9.5. β-actin, loading control. (D) Litter ratio analysis for E8.5-9.5 (left) and weaned (right) *Thoc6^fs/fs^* mice. Ratios are consistent with embryonic lethality of homozygous frameshift mice. n = 164 mice (left); n = 27 mice (right). (E) Immunostaining of markers Ph3 and C.Casp3 in E9.5 mouse forebrain. Illustration highlights sectioning and quantification approach. (F) Quantifications of fractions of Pax6, Ph3, and C.Casp3-expressing cells in E9.5 neuroepithelium. Measurements were combined from one rostral and one caudal section (from two lateral segments depicted by solid black boxes in E) per three embryo replicates per genotype. Data shown as mean ±SEM. Significance, two-tailed *t* test.

Forebrain tissues of E8.5-E10.5 *Thoc6* littermates were characterized to identify neurodevelopmental changes in *Thoc6^fs/fs^* pups. Immunohistochemistry of the telencephalic vesicles revealed a consistently thinner neuroepithelium (PAX6) in *Thoc6^fs/fs^* compared to *Thoc6^+/+^* littermate controls (Figures 3E,3F, S2B, and S2D). While E9.5 neuroepithelium showed a relative increase in mitotically active cells (PH3), widespread apoptosis (Cleaved Caspase-3 (C.CASP3)) was noted (Figures 3E and 3F). These findings are consistent with proliferative defects noted in *Thoc6^fs/fs^* tissue, suggesting THOC6 is important for corticogenesis.

### Global mRNA export is not altered in THOC6 models of human *in vitro* neural development

Given the prominent link between the THO subcomplex and RNA export in current literature, we first sought to investigate the impact of THOC6 variants on RNA nuclear export functions in human neural progenitor cells (hNPCs) differentiated from hPSCs (Figure 4A). NPCs are the embryonic cell population frequently implicated in the developmental mechanism of primary microcephaly, a TIDS clinical feature. Defects in mRNA export are typically observed as differential accumulation of polyadenylated (polyA+) mRNA in the nucleus, enriched at nuclear speckle domains (Bahar Halpern et al., 2015). Standard oligo-dT fluorescent *in situ* hybridization (FISH) was performed on hNPCs to visualize polyA+ mRNA signal in nuclear and cytoplasmic cellular fractions (Figures S3A, S3B, and S3C). Comparison of the nuclear-to-cytoplasmic (N/C) polyA+ signal intensity ratios across genotypes indicates a slight reduction in *THOC6* affected samples, suggesting a trend towards nuclear reduction in affected hNPCs relative to *THOC6^+/+^* and heterozygous unaffected control hNPCs (Figure S3C). The significance of this modest export finding is evident when the N/C polyA+ signal intensity ratios are compared to *THOC6^+/+^* hNPCs treated with wheat germ agglutinin (WGA), a potent inhibitor of all nuclear pore transport and the positive control for export changes (Mor et al., 2010) (Figure S3A), Relative to all *THOC6* genotypes, WGA treated hNPCs have a significantly higher N/C ratio (Figure S3C) attributed to strong polyA+ mRNA accumulation in the nucleus (Figure S3B). Although bulk mRNA export is largely unaltered in *THOC6*-affected hNPCs, a THOC6 dependent tetramer TREX export cannot be ruled out for specific polyA+ or in mRNP processing functions upstream of export—TREX functions that have not been explored in hNPCs.

**Figure 4.**
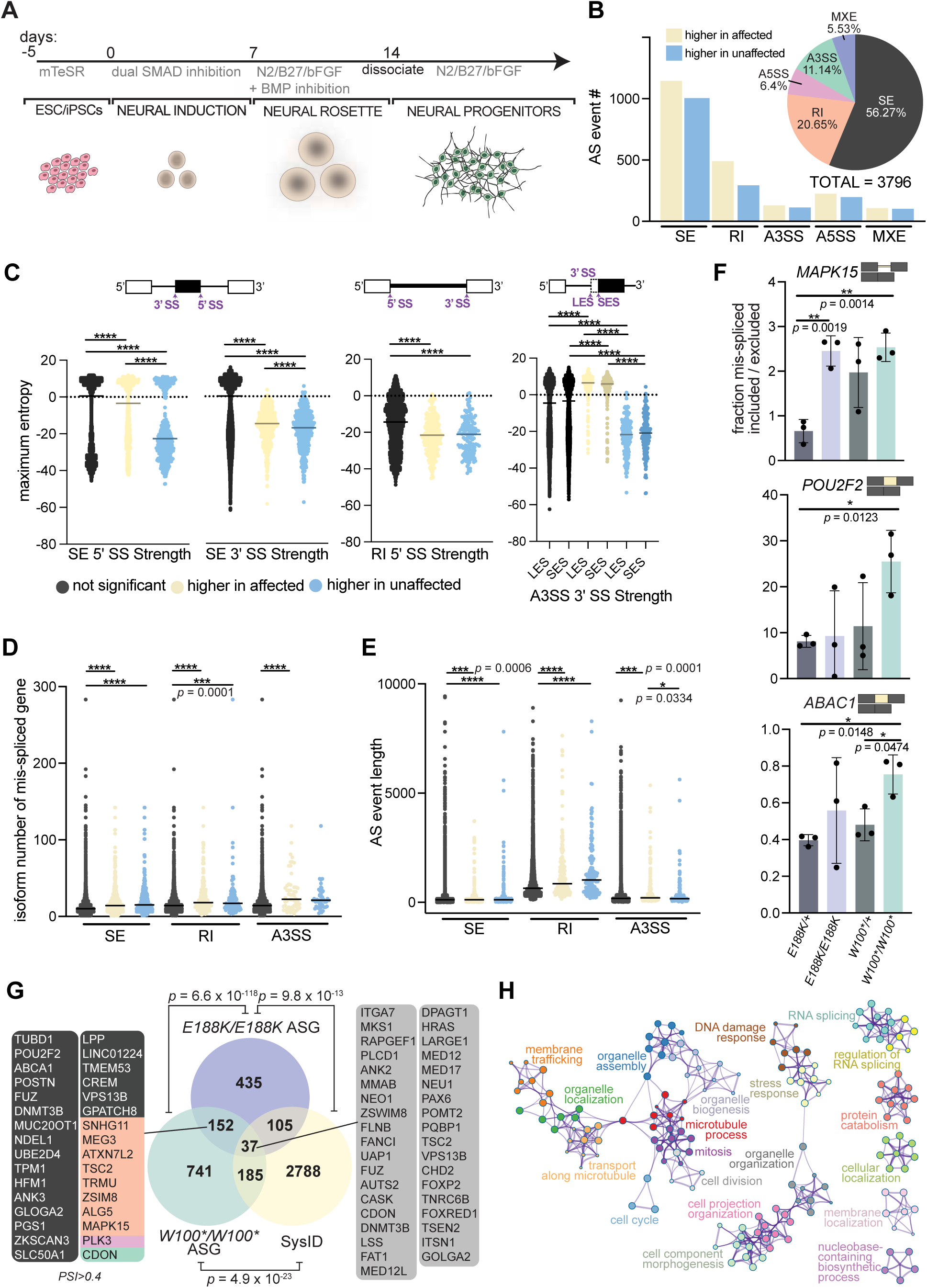
Characterization of alternative splicing events in *THOC6* affected hNPCs. (A) Differentiation protocol to derive human neural progenitor cells from affected and unaffected ESC/iPSCs. (B) Combined rMATS summary results for AS events in *THOC6^W100*/W100*^* and *THOC6^E188K/E188K^* hNPCs relative to *THOC6^W100*/+^* control hNPCs. Event type (pie chart) and inclusion status (bar chart). Yellow, higher inclusion in affected. Blue, higher inclusion in unaffected. (C) Significant splice site strength score differences at mis-spliced events in affected hNPCs based on maximum entropy model. (D) transcript number per AS gene and (E) AS event length in *THOC6^W100*/W100^* and *THOC6^E188K/E188K^* vs. *THOC6^W100*/+^* NPCs. (F) RT-PCR AS validations of SE (*ABAC1*, *POU2F2*) and RI (*MAPK15*) events in three additional biological replicates of hNPCs per genotype with quantified mis-spliced ratios. Data shown as mean ±SEM. P-values (C-F), two-tailed unpaired *t* test. ****, *p* = < 0.0001. (G) Venn diagram of overlap of *THOC6 ^W100*/W100*^* and *THOC6^E188K/E188K^* AS genes and all syndromic intellectual disability genes included in the SysID database. Overlap significance tested by Fisher’s exact test. ASG, alternatively spliced genes. Metascape analysis on combined significant mis-spliced events (FDR <0.05) in *THOC6^E188K/E188K^* and *THOC6^W100*/W100*^* NPCs (H).

### THOC6 depletion reveals TREX function in pre-mRNA splicing in hNPCs

Proper mRNP processing, including mRNA splicing, is required for TREX-dependent RNA nuclear export, linking these steps in mRNA biogenesis. Co-transcriptional recruitment of TREX to the 5’ end of maturing mRNPs coupled with described TREX associations with splicing factors (Viphakone *et al.*, 2019) led us to investigate features of mRNA processing for vulnerability to loss of THOC6. To capture RNA processing differences caused by loss of THOC6 function, we performed RNA-sequencing (RNAseq) on ribosomal (r)RNA-depleted RNA extracted from wildtype and heterozygous unaffected and homozygous affected hNPCs (Table S4). Principal components analysis (PCA) of RNAseq data demonstrates reproducibility across, and distinct transcriptomic differences between the affected hNPC replicates compared to the unaffected control hNPC replicates (Figure S4A). Genotype driven differential expression and splicing changes were assessed. To investigate a THOC6-dependent role for TREX in splicing, comparative splicing analysis was carried out using the rMATS pipeline on biallelic *THOC6^E188K/E188K^* and *THOC6^W100*/W100*^* samples versus heterozygous controls (Table S5) (Shen et al., 2014). A combined total of 3,796 significant alternative splicing (AS) events were detected in affected hNPCs, representing the major AS types: skipped/cassette exon (SE), alternative 5’ splice site (A5SS), alternative 3’ splice site (A3SS), retained intron (RI), and mutually exclusive exon (MXE). The most overrepresented AS events observed in affected cells were SE (56%, 2136 of 3796) and RIs (21%, 784 of 3796). The high frequency of RIs is notable and unique relative to splicing defects identified in LOF models of other splicing factors, as well as for normal splicing patterns in neural development (Figure 4B) (Chai et al., 2021; Ellis et al., 2012; Jin et al., 2020; Licatalosi et al., 2008; Llorian et al., 2010; Weyn-Vanhentenryck et al., 2018). AS events in affected hNPCs show comparable inclusion and exclusion of AS junctions, with a slight trend towards inclusion due in part to the high frequency of RIs (Figure 4B). SE and RI splicing events occur by distinct molecular mechanisms, mediated by EJC pathways. Detection of defects in both splicing categories suggests the THO tetramer serves as a molecular platform for coordinating complex splicing events, as opposed to regulation of a specific subset of splicing events controlled by association with and function of individual RNA splicing factors. In agreement with this finding, we did not find consistent motif enrichment for specific RNA-binding proteins at AS junctions. These findings implicate a novel role for THOC6-dependent TREX splicing in mRNP processing in hNPCs.

Since AS motif enrichment analysis of THOC6-affected and control hNPCs transcriptomic data did not reveal trans-regulatory elements responsible for the differential splicing patterns, it was posited that cis-elements may underlie these differences. A maximum entropy model that assesses short sequence motif distributions was used to test the strength of the donor (5’) and acceptor (3’) of AS events (Yeo and Burge, 2004). A general trend towards weaker splice sites were detected at differential SE, RI, and A3SS events in affected cells (unpaired two-tailed *t* test, Figure 4C). The SE, RI, and A3SS events were enriched in genes with a disproportionately high number of isoforms that show dependence on weak, alternative/cryptic splice sites to facilitate isoform diversity (Figure 4D) (Wang et al., 2015). RI events in affected hNPCs also had weaker splice sites compared to controls, suggesting that THOC6 deficiency induces mis-splicing at weak splice sites. In addition, the AS SE, RI, and A3SS events in affected *THOC6* hNPCs impacted exons/introns that are significantly longer than nonsignificant events (Figure 4E). Likewise, the length of introns retained in RI events were significantly longer, with a 1.4-fold increase in length quantified for significant RI events (P=<0.0001, unpaired two-tailed *t* test, Figure 4E). Lastly, no positional bias was observed for AS events (Figure S4D and S4F). To validate our bioinformatic analysis, AS inclusion trends in select, top, shared events were validated by qRT-PCR, demonstrating a high correlation (unpaired two-tailed *t* test, Figure 4F). Together, the detected RNA processing signature across diverse SE and RI events at weak splices sites suggest impaired splicing fidelity from loss of THOC6.

To investigate the role of RNA misprocessing in ID pathology, we intersected our AS events with the genes that are known to cause syndromic ID, deposited in the SysID database (SysIDdb). 152 genes with significant AS events included or excluded in >10% of transcripts in hNPCS were detected in nonsense and missense affected genotypes (Figure 4G). 185 AS genes in *THOC6^W100*/W100*^* and 105 AS genes in *THOC6^E188K/E188K^* hNPCs are known genes causative for syndromic ID represented in the SysIDdb (Figure 4G). Aberrantly spliced ID genes were identified in both THOC6 affected genotypes, consistent with a role for THOC6 in ID. 37 ID genes (1.3% of SysIDdb) are AS in both affected genotypes, identifying genes for shared mechanisms that may preferentially contribute to TIDS pathology. To identify biological mechanisms implicated by THOC6-dependent AS, biological pathway enrichment analysis was performed on mis-spliced genes in affected cells. Genes with differential splicing were significantly enriched for functions in RNA splicing, cell projection organization, membrane trafficking, organelle organization, mitosis cell cycle, and DNA damage response (Figure 4H). RNA processing is tightly controlled by feedback loops (e.g., auto-repression by poison exons or intron retention), which would explain how effects on *cis* elements may lead to changes in *trans* factors (i.e., AS events in splicing regulatory factors).

### THOC6-dependent mRNP processing is required for hNPC proliferation and differentiation

Alternative mRNA processing events, such as SE and RI in mRNA splicing, can impact gene expression through different mechanisms; isoform ratio differences versus inclusion of premature termination codons that initiate NMD. To investigate the impact of AS events on expression, we performed differential gene expression analysis on *THOC6^E188K/E188K^* versus *THOC6^W100*/+^*, and *THOC6^W100*/W100*^* versus *THOC6^W100*/+^* dyads (Figure S5B). Among the 336 differentially expressed genes (DEGs) in *THOC6^E188K/E188K^* hNPCs, only 13 had splicing defects (*p* = 5.3×10^−3^, Fisher’s exact test) compared to 46 mis-spliced DEGs (of 661 DEGs, *p* = 4.2×10^−7^, Fisher’s exact test) in *THOC6^W100*/W100*^* hNPCs, indicating a subtle effect of mis-splicing on expression (Figure 5A). Notably, there are nearly double the number of AS genes (ASGs) (435 E188K; 741 W100*) and DEGs (336 E188K; 661 W100*) in *THOC6^W100*/W100*^* hNPCs, suggesting that the nonsense genotype has a greater impact on splicing and gene expression (Figure 5A). Relevant for TIDS pathology, 20% (68 of 336; *p* = 1.5×10^−5^, Fisher’s exact test) of *THOC6^E188K/E188K^* DEGs and 18% (118 of 661; *p* = 9.7×10^−6^, Fisher’s exact test) of *THOC6^W100*/W100*^* DEGs are syndromic ID genes, which conveys important information pertinent for understanding the pathogenic mechanisms of TIDS (Figure 5A).

**Figure 5.**
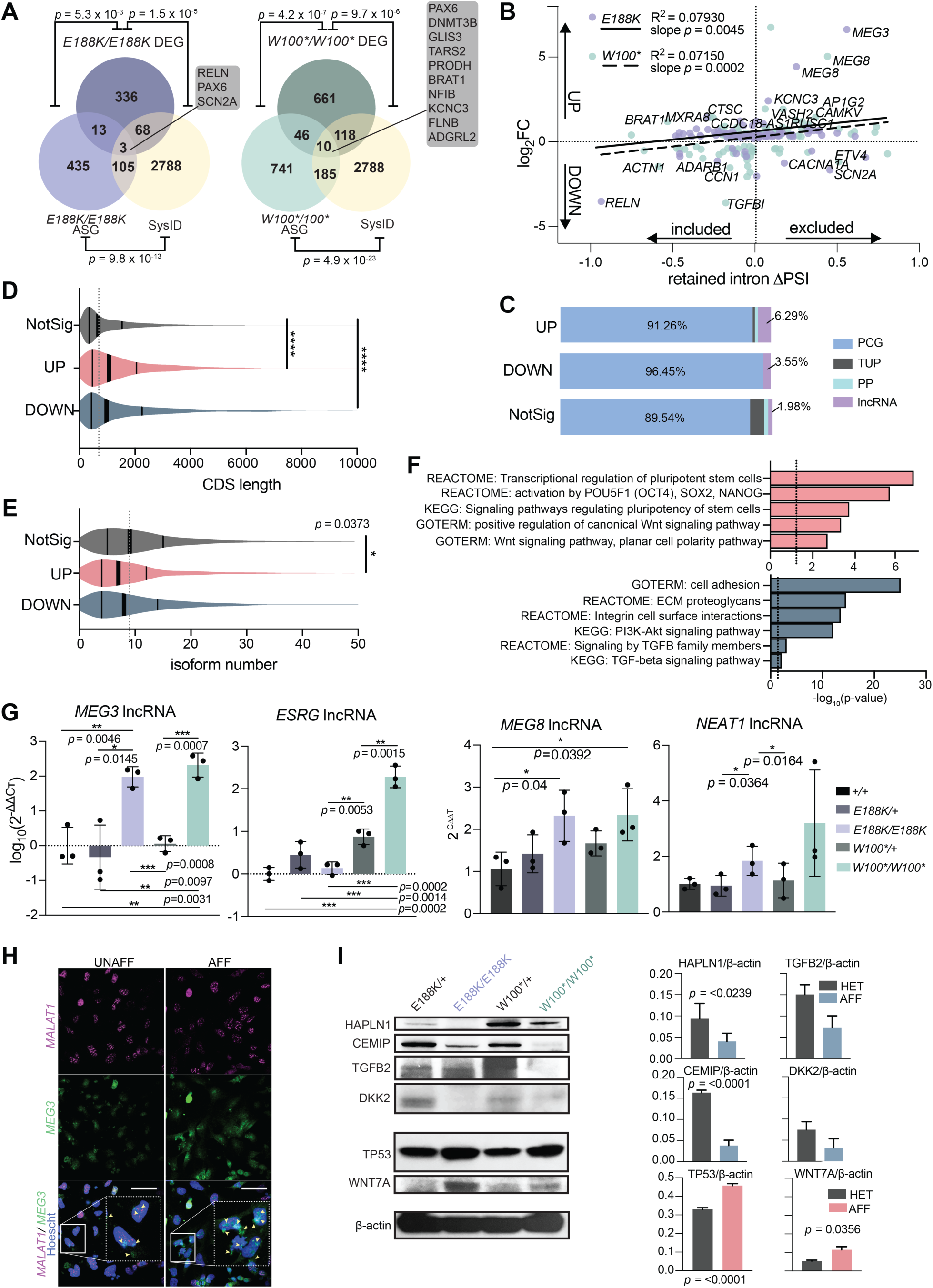
Differential expression analysis in affected hNPCs. (A) Venn diagram of gene overlap of *THOC6 ^W100*/W100*^* and *THOC6^E188K/E188K^* affected genes and all syndromic intellectual disability genes included in the SysID database. Overlap significance tested by Fisher’s exact test. DEG, differentially expressed genes; ASG, alternatively spliced genes. (B) Linear regression analysis of log_2_foldchange and Δ percent transcripts spliced in (PSI) for significant retained intron events in affected cells. Purple dots indicate *THOC6^E188K/E188K^* hits and green dots indicate *THOC6 ^W100*/W100*^*. Best fit line, R^2^, and slope p-value for *THOC6^E188K/E188K^* (solid line) and *THOC6 ^W100*/W100*^*(dotted line). (C) Percentage of gene type by condition for combined DEGs in affected hNPCs. NotSig, not significant; Up, upregulated. Down, downregulated; lncRNA, long non-coding RNA; PP, processed pseudogene; TUP, transcribed unprocessed pseudogene; PCG, protein coding gene. Violin plots of coding sequence (CDS) length (D) and isoform number (E) of combined DEGs in affected cells compared to non-significant genes. (F) DAVID biological pathway enrichment analysis of combined upregulated genes (top, red) and downregulated genes (bottom, blue) in *THOC6* affected hNPCs. (G) qPCR relative - abundance quantifications (2^−ΔΔCt^) for *MEG3*, *ESRG*, *MEG8*, and *NEAT1* in hNPCs. Three technical replicates of three biological replicates per genotype. (H) RNA FISH probing for *MEG3* and *MALAT1* in affected and unaffected hNPCs. Cell inset showing *MEG3* expression and localization differences with yellow arrows in merged image. Scale bar 50 μm. UNAFF, unaffected (W100*/+); AFF, affected (W100*/W100*). (I) Protein abundance of top downregulated and upregulated genes across genotypes. Genes labeled with log_2_FC > 1 or < −1 and PSI > 0.1 or < −0.1. Data shown as mean ±SEM. Significance, two-tailed unpaired *t* test.

Retained intron AS events are often subject to NMD in protein-coding genes (PCGs), while intron inclusion in lncRNAs alters nuclear export and conformation. Given the high number of RI events detected in *THOC6* affected hNPCs, we tested the correlation between differential expression and RI events. For this analysis, gene-level mRNA abundance fold-change in affected hNPCs was correlated to the change in intron inclusion within transcripts from PCGs or non-coding RNA loci, referred to as percentage spliced-in (ΔPSI). We observed a trend in both *THOC6-*affected genotypes that lower gene expression correlates with greater intron inclusion (slope, *p* = 0.0045 for *THOC6^W100*/W100*^* and *p* = 0.0002 for *THOC6^E188K/E188K^*, simple linear regression) (Figure 5B). Conversely, the quadrant representing differential intron exclusion and elevated expression was prominently represented by lncRNAs, where intron exclusion is implicated in impaired lncRNA function. The three significantly dysregulated ASGs represented in both affected genotypes were *MEG3*, *PAX6*, and *POSTN*. Consistent with the observed trend between RI events and expression, analysis of all DEGs revealed that PCGs make up the largest portion of DEGs (with a greater portion, 96.45% affected in downregulated genes), while the portion of lncRNAs is highest in upregulated genes (6.29%; compared to 1.98% of non-significant genes), reflecting the molecular differences between these distinct mRNA subtypes.

Additional mRNA characteristics that may account for a portion of the observed differential expression are gene length and isoform number. In affected hNPCs, significantly more transcripts from long genes with on average of less than 10 annotated isoforms were identified compared to non-significant genes (Figures 5D and 5E). The trend towards DEGs with fewer transcript isoforms in affected hNPCs suggest alternatively spliced transcripts are more stable in affected cells (Figure 5C). These findings again reflect a requirement for the larger THOC6-dependent TREX tetramer complex function in facilitating mRNP processing of long mRNAs with high expression in brain.

To identify biological pathways predicted to contribute to *THOC6* neuropathology, DAVID analysis was performed to identify biological categories defined by DEGs in*THOC6* affected hNPCs (Figure 5D). Downregulated genes are enriched in integrin cell adhesion, extracellular matrix interactions, PI3K-AKT signaling, and TGF-β signaling pathways, which are critical for brain development (Figure 5F). PI3K-AKT/mTOR signaling regulates cortical NPC proliferation, differentiation, and apoptosis (Andrews et al., 2020; Li et al., 2017). Over 30 genes attributed to the PI3K-AKT/mTOR signaling pathway were downregulated in affected cells, accounting for the significant enrichment (*p* = <1 x 10^−13^). *HAPLN1*, *MYC*, *BMPR1B*, *DCN*, *FBN1*, *INHBA*, *ID4*, *THBS1*, *TGFB2,* DEGs enriched in the TGF-β signaling pathway (*p* = <0.001), have direct implications for TGF-β signaling in neural induction, differentiation, and NPC fate specification in TIDS developmental mechanisms (Meyers and Kessler, 2017; Vogel et al., 2010). Complementary pathways enriched with upregulated DEGs implicate multipotency (*OCT4*, *PAX6*), proliferation, neuron differentiation and WNT signaling pathways (*p* = <1 x 10^−6^, *p* = <0.001, *p* = <0.001, and *p* = <0.001, respectively). WNT signaling is known to promote NPC self-renewal expansion during corticogenesis (Harrison-Uy and Pleasure, 2012; Qu et al., 2013). Shared dysregulation of mTOR, TGF-β, and WNT signaling, coupled with upregulation of multipotency factors in affected genotypes, suggests defects in hNPC multipotency and neural differentiation underlie TIDS pathogenesis.

To identify transcription factor networks dysregulated in *THOC6*-affected hNPCs, transcription factor motif enrichment analyses was performed. Significant enrichment of MEF2, LHX3, and SRF target genes was observed in heterozygous controls compared to both affected genotypes (Figure S5D). Using a second analysis tool, ChEA3, differential expression of SOX, FEZF, FOX, and GLI target genes, and downregulation of HEYL, TWIST, FOX, MEOX2, PRRX2, and MKX target genes were enriched in affected hNPCs (Figure S5E). These transcription factor networks are important for neuronal differentiation and fate specification (Jalali et al., 2011; Tsui et al., 2013; Wang et al., 2011), concordant with GSEA findings. Together, these results suggest that gene expression programs that modulate timing of the switch from neural proliferation to differentiation are altered in TIDS.

To refine specific candidate genes implicated in shared TIDS neuropathology, DEGs between affected *THOC6* genotypes were intersected. 12 genes were upregulated and 117 were downregulated in affected hNPCs, with notable lncRNAs represented. Significant enrichment was detected in Integrin 1 pathway and extracellular matrix protein interaction networks (Figure S5C). Using mRNA obtained from three additional replicate differentiations of hNPCs per genotype, significant upregulation of *MEG3*, *MEG8*, *ESRG*, and *NEAT1* lncRNAs was confirmed by qRT-PCR (Figures 5G). RNA FISH confirmed increased expression of *MEG3* in affected hNPCs compared to controls, with elevated signal observed in both nuclear and cytoplasmic fractions (Figure 5H). Upregulation of functional lncRNAs *NEAT1* and *MEG3* has been linked to activation of WNT activation and suppression of TGF-β signaling, respectively (Cui et al., 2019; Mondal et al., 2015). Concordant with these findings, the protein level of WNT and TGF-β signaling components in *THOC6-*affected hNPCs exhibit a corresponding differential up- and down-expression relative to controls. Specifically, WNT signaling components WNT7A and TP53 showed increased protein expression, with higher abundance detected in affected hNPCs (Figure 5I). TGF-β pathway proteins HAPLN1 and TGFB2 also showed reduced protein expression in affected hNPCs together with high CEMIP and DKK2 (Figure 5I). We propose that loss of THOC6 leads to lncRNA-mediated dysregulation of key developmental signaling pathways which has implications for the balance of proliferation and differentiation during neural development.

### Apoptotic upregulation and retained intron enrichment in *Thoc6^fs/fs^* E9.5 mouse forebrain

To investigate the conserved THOC6-dependent TREX functions that account for divergent phenotypic outcomes between mammalian models, mRNP processing was assessed in E9.5 mouse brain using complementary RNAseq experiments to those performed in hNPCs (Figures 6A and S6A). Three biological replicates were analyzed per genotype (*Thoc6^+/+^*, *Thoc6^fs/+^*, *Thoc6^fs/fs^*). Fewer significant AS events (FDR <0.05) were detected in *Thoc6^fs/fs^* E9.5 brain than in affected hNPCs, but the pattern of AS events was recapitulated, with the majority of AS events categorized as SE (45%) and RI (26%) (Figure 6B). Greater than 40 PSI was quantified in the *Thoc6^fs/fs^* transcriptome and retained and excluded intron events in *Cenpt*, *Admts6*, and *Fam214b* were validated (Figures 6C and S6B). Maximum entropy model analysis of splice junctions revealed significantly weaker 3’ splice site strengths for SE events and weaker 5’ splice sites associated with RI events in the *Thoc6^fs/fs^* mouse model. While this signature of splice site weakness is more modest in mouse than in human THOC6 models, these findings suggest a conserved role of THOC6-dependent TREX tetramer in coordinating mRNA processing that precedes TREX export functions (Figures 6D and S6C).

**Figure 6.**
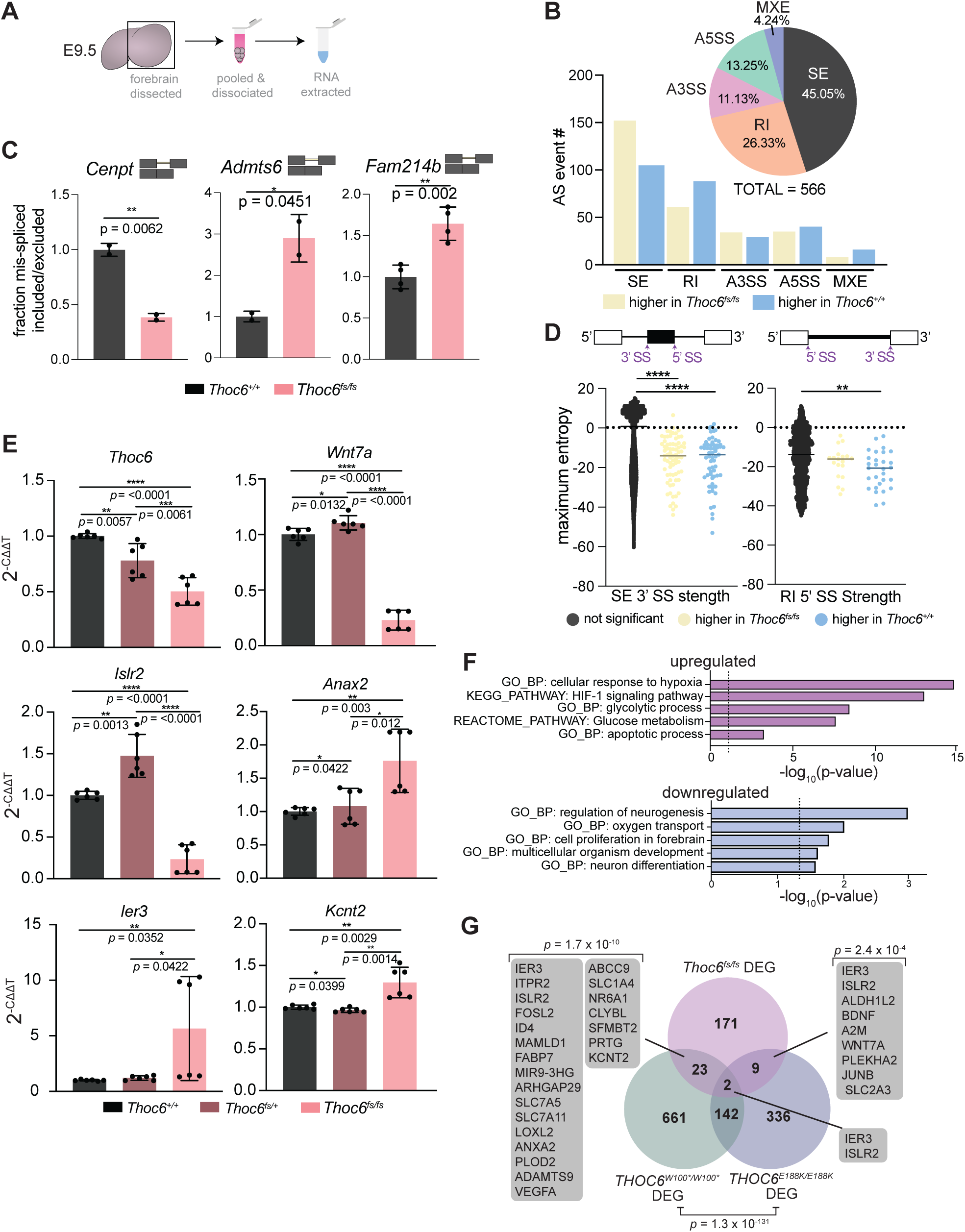
Characterization of mRNA processing defects in *Thoc6^fs/fs^* mouse E9.5 forebrain. (A) Cartoon of E9.5 mouse forebrain total RNA sample preparation. (B) rMATS summary results for AS events in *Thoc6^fs/fs^* E9.5 forebrain. Event type (pie chart) and inclusion status (bar chart). Yellow, higher inclusion in *Thoc6^fs/fs^*. Blue, higher inclusion in *Thoc6^+/+^*. (C) Quantifications from RT-PCR validating top AS events *Cenpt*, *Admts6*, *Fam214b* in 2-4 biological replicates. Significance, two-tailed unpaired *t* test. Data are mean ±SEM. (D) Significant splice site strength score differences at mis-spliced events in *Thoc6^fs/fs^* samples based on maximum entropy model. (E) RT-qPCR validations of *Thoc6*, *Wnt7a*, *Islr2*, *Ier3*, *Kcnt2*, *Anax2* mRNA abundance on two additional biological replicates of E9.5 forebrain per genotype. Three technical replicates analyzed per sample. Significance, two-tailed unpaired *t* test. Data are mean ±SEM. (F) DAVID analysis showing significantly enriched biological pathways among upregulated genes (top, magenta) and downregulated genes (bottom, blue) in *Thoc6^fs/fs^* E9.5 forebrain. (G) Venn diagram of overlap of DEG in affected hNPCs and *Thoc6^fs/fs^* mouse E9.5 forebrain. Overlap significance tested by Fisher’s exact test.

Notably, biological pathway and network enrichment analysis of AS genes identified mRNA processing, pre-miRNA processing, de-adenylation of mRNA, central nervous system development, forebrain development, multicellular growth, response to oxidative stress, cytoskeletal organization, and neuron projection (Figure S6D) — several of the biological categories associated with hNPCs ASGs. These shared findings suggest selective conservation of mRNP processing mechanisms by *THOC6* in mouse and human forebrain.

To assess the correlation between THOC6 mRNP processing defects and expression, differential expression analysis of *Thoc6^fs/fs^* forebrain mRNA sequencing data was performed compared to *Thoc6^+/+^* controls. Of note, *Thoc6* mRNA is downregulated (two-fold, *p*=<0.0001), consistent with NMD (Figures 6E and S6F). In this model, 5x more genes were upregulated (144 genes) than downregulated (27 genes). Nevertheless, downregulated genes may covey important pathology. First, downregulated genes functionally converge on neurogenesis, proliferation, and differentiation pathways (Figure 6F). Upregulated genes are implicated in the hypoxic response, HIF-1 signaling pathway, and glycolysis—biological categories indicative of increased apoptosis in affected cells (Figure 6F). To investigate if altered transcription factor networks contribute to pathway dysregulation, we performed GSEA transcription factor motif enrichment analysis. *HIF1*, *NRSF*, *SMAD3*, and *STAT3* target genes were enriched in *Thoc6^fs/fs^* E9.5 forebrain (Figure S6G). HIF-1 and STAT3 can induce apoptosis in response to hypoxia (Greijer and van der Wall, 2004; Zhou et al., 2021), which is consistent the observed elevation of apoptosis in *Thoc6^fs/fs^* E9.5 neuroepithelium (Figures 3E and 3F). SMAD3 signaling is activated by TGF-β to promote cortical differentiation (Vogel *et al.*, 2010), suggesting shared disruption of TGF-β signaling in both human and mouse model systems.

DEGs shared between mouse and human model systems are consistent with conserved TIDS molecular pathology (Figure 6G). More *Thoc6^fs/fs^* DEGs overlapped with *THOC6^W100*/W100*^* (23 genes) than *THOC6^E188K/E188K^* (9 genes) samples, and include genes involved in neurogenesis, hypoxic response, and synapse regulation. Validation of *Ier3*, *Islr2, Wnt7a*, *Kcnt2*, *Anax2*, and *Vegfa* DEGs shared across affected models were confirmed by qRT-PCR in three additional E9.5 forebrain biological replicates for *Thoc6^+/+^*, *Thoc6^fs/+^*, and *Thoc6^fs/fs^* samples (Figures 6E and S6H). Overlapping affected human and mouse molecular mechanisms suggest shared pathology. However, the extent of upregulation of genes in response to increased apoptosis is exacerbated in mouse, highlighting species-specific phenotypic differences due to loss of THOC6.

### Delayed differentiation and elevated apoptosis in *THOC6*-affected forebrain organoids

*THOC6* pathogenesis in human cortical development was investigated using dorsal forebrain-fated organoids, neural differentiated from iPSC lines (Figure 7A). Forebrain organoids recapitulate the cellular heterogeneity and developmental dynamics of early corticogenesis (Qian et al., 2016). Within each organoid, several neural rosette (NR) structures develop stochastically to recapitulate features of *in vivo* ventricular zone development, including hNPC proliferation and differentiation to cortical neuron fates (Figure 7A). NR morphology was evaluated in cortical organoids at 28 days of neural differentiation (ND) from three independent differentiations per genotype. To minimize the effect of inter-cell line NR variability, the following analyses focus on heterozygous unaffected and homozygous affected comparisons. To characterize the NR proliferative niche, the maximum thickness of the NR neuroepithelial center was measured as defined by N-cadherin immunostaining and pseudostratified NR cytoarchitecture by Hoescht staining (Figures 7B, 7C, S7A, and S7B). *THOC6*-affected organoids show significantly thinner pseudostratified neuroepithelium (*p* = <0.0001, two-tailed *t* test), concordant with reduced NR size composed of fewer cells (*p* = <0.0001, two-tailed *t* test) (Figures 7B, 7C, S7A, and S7B). Given the stark upregulation of apoptosis observed in *Thoc6^fs/fs^* E9.5 mouse forebrain, we investigated the contribution of apoptosis as a mechanism of reduced NR size in the *THOC6*-affected organoids (Figures 7B, 7D, S7A, and S7B). A significantly higher proportion of affected NR cells expressed the apoptotic marker C.CASP3, providing evidence for shared mechanism of altered corticogenesis (*p* = <0.0001, two-tailed *t* test) (Figure 7D).

**Figure 7.**
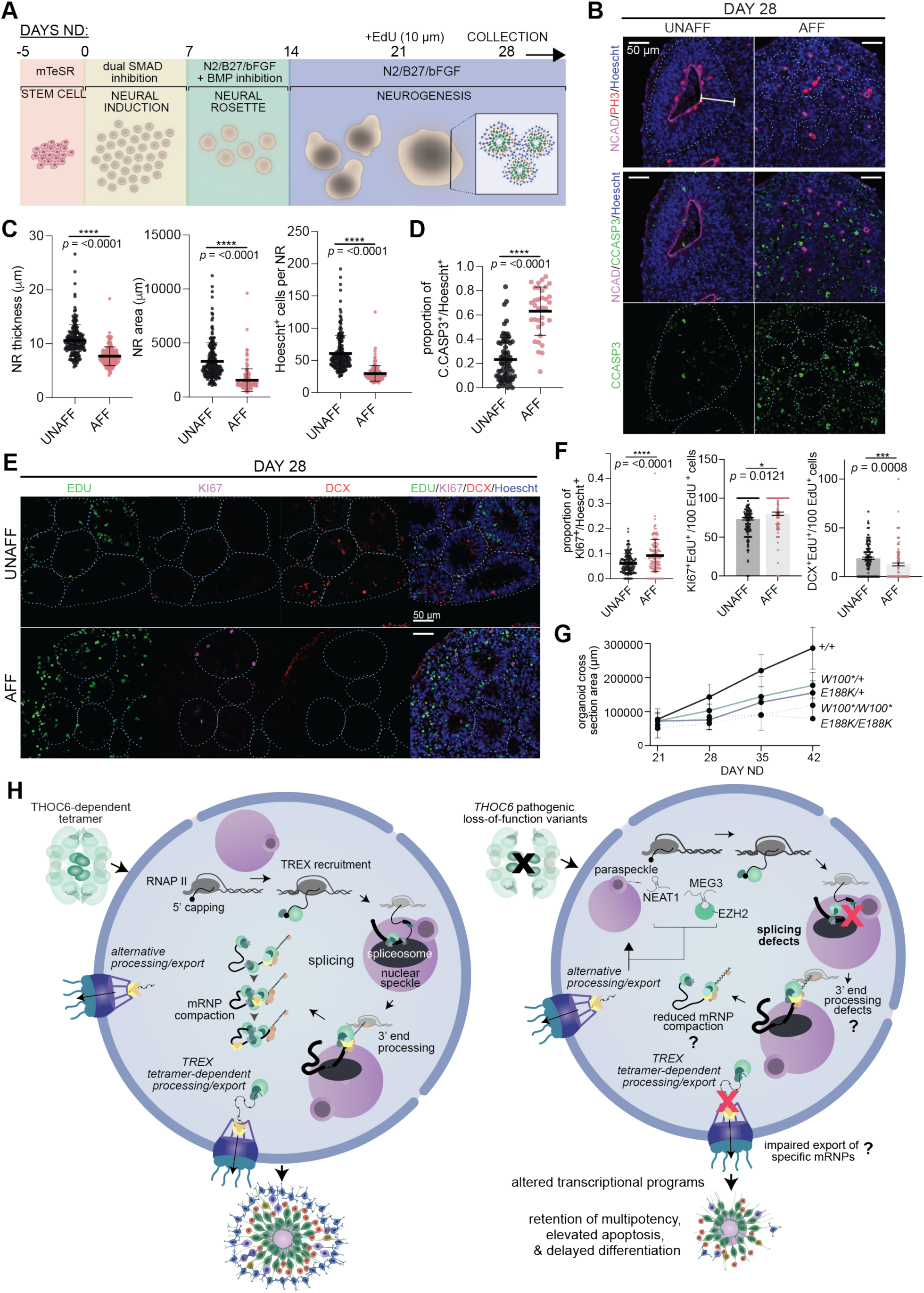
Modeling of *THOC6* variant pathogenesis in human cerebral organoids. (A) Cerebral organoid differentiation protocol. ND, neural differentiation. (B) Immunostaining of PH3, N-Cadherin, apoptosis marker cleaved caspase3 (C.CASP3), and Hoescht in day 28 human cerebral organoids differentiated from unaffected and affected iPSCs, highlighting differences in neural rosette morphology. 40x magnification; Scale bar: 50 μm. UNAFF, unaffected; AFF, affected. (C-D) Quantifications of area, thickness, Hoescht+ cells, and fraction of C.CASP3+ cells per NR for *THOC6^W100*/+^* and *THOC6^E188K/+^* controls and *THOC6^W100*/W100*^* and *THOC6^E188K/E188K^* affected organoids. NR (organoid) number analyzed across one differentiation replicate per genotype: unaffected, *n* = 67 (15); affected, *n* = 34 (10). (E) Immunostaining of EDU, KI67, DCX to assess timing of differentiation in day 28 organoids with quantifications (F). NR (organoid) number analyzed across three differentiation replicates per genotype: unaffected, *n* = 187 (87); affected, *n* = 157 (67). (G) Growth rate of organoids across genotypes measured by cross section area (μm) from days 21-42. (C-D, F-G) Data shown as mean ±SEM. Significance, two-tailed unpaired *t* test. Representative image for unaffected and affected are *THOC6^E188K/+^* and *THOC6^E188K/E188K^*, respectively. (H) Schematic of proposed model of *THOC6* pathogenesis.

To assess alterations in the timing of differentiation in affected NRs, we performed EdU-pulse labeling at day 21ND for 24 hours to label mitotically active cells, followed by organoid immunohistochemistry analysis at day 28ND (Figures 7E, 7F, S7C, and S7D). To assess the balance of multipotency and differentiation EdU, KI67, and DCX co-labeled cells per NR were quantified. A significant increase in cells co-stained with the proliferation marker KI67 and EdU per affected NR were detected at day 28ND (*p* = <0.0001, two-tailed *t* test), indicating affected NPCs remain mitotically active longer than control NPCs (Figure 7F). This finding paired with elevated mRNA and protein expression of OCT4 in affected hNPCs (data not shown) supports retention of multipotency model. Consistent with this finding, we observed a significant reduction in the fraction of EdU cells co-labeled with the migrating neuron marker doublecortin (DCX) in affected NRs (*p* = <0.0001, two-tailed *t* test) (Figure 7F). Paired with the prolonged proliferation dynamics, this suggests a disruption to the differentiation timeline in affected organoids.

To investigate effects of reduced NR growth on organoid size, we measured whole organoid cross section areas weekly from day 21 to day 42. Compared to the steady size increase of *THOC6^+/+^* organoids, affected organoids showed a slower growth rate (E188K/E188K: *p* = 0.0122; W100*/W100*: *p* = 0.0362) (Figure 7G). Together, our findings implicate a pathogenic mechanism of delayed differentiation due to reduced NPC proliferative capacity and elevated apoptosis with subsequent cortical growth impairment in affected organoids.

## DISCUSSION

There is a growing cohort of individuals with TIDS due to variant discovery resulting from exome based genetic testing (Accogli *et al.*, 2018; Amos *et al.*, 2017; Anazi *et al.*, 2016; Anazi *et al.*, 2017; Beaulieu *et al.*, 2013; Boycott *et al.*, 2010; Casey *et al.*, 2016; Gupta *et al.*, 2020; Hassanvand Amouzadeh *et al.*, 2020; Kiraz *et al.*, 2022; Mattioli *et al.*, 2018; 2019; Ruaud *et al.*, 2022; Zhang *et al.*, 2020). However, our understanding of the molecular functions of THOC6 are limited, despite robust research on the TREX complex (Chi *et al.*, 2013; Dias et al., 2010; Guria *et al.*, 2011; Heath *et al.*, 2016; Katahira et al., 2009; Katahira et al., 2013; Luna *et al.*, 2012; Maeder *et al.*, 2018; Mancini et al., 2010; Masuda *et al.*, 2005; Peña *et al.*, 2012; Pühringer *et al.*, 2020; Rondón *et al.*, 2010; Tran et al., 2014; Viphakone *et al.*, 2019; Wickramasinghe and Laskey, 2015; Zuckerman et al., 2020). Information implicating important species differences in TREX composition and function that are THOC6-dependent serve to highlight this gap in knowledge (Chávez et al., 2000; Jimeno et al., 2002; Meinel et al., 2013; Strässer et al., 2002). Our findings support a novel *THOC6* LOF model of TIDS pathogenesis whereby pathogenic variants impair the formation of the THO/TREX tetramer complex that facilitates multivalent protein-mRNA interactions to support coordination of mRNP processing and export functions (Figure 7H). The resulting molecular impact preferentially disrupts processing of long mRNAs, with complex splicing patterns that are compounded by genetic feedback loops that regulate splicing. The neurological features of TIDS and the enrichment of long genes expressed in the brain further implicate a critical role for the THOC6-dependent TREX tetramer in providing a platform for enhanced coalescence of mRNP processing cofactors and maintenance of mRNA structural integrity during splicing of long mRNA transcripts important for brain development. Cell-type and organism specific requirements for splicing and gene length are predicted to inform the variation in tolerance that underlies distinct human phenotypic presentations and interspecific differences.

The crystal structure of human THO-UAP56 was recently solved, implicating several putative consequences of THOC6-dependent TREX tetramer disruption (Figure 7H) (Pühringer *et al.*, 2020). Defects in TREX tetramer assembly are not predicted to disrupt formation of stable functional dimers, allowing THOC6-depleted models to discriminate between dimer and tetramer functions. The tetramer affords greater surface area for mRNP processing and permits enhanced co-adaptor loading of known and potentially species-specific TREX cofactors in species with a substantial splicing burden. Certainly, we see evidence for reduced association of ALYREF with THO complexes by THOC6 co-immunoprecipitation in *THOC6*-affected iPSCs, providing indirect evidence for altered TREX tetramer formation with impaired binding of tetramer-associated mRNP processing cofactors (Figure 2F). Tetramer formation juxtaposes two UAP56 helicases on each end of dimers (Chang *et al.*, 2013) to support a greater number of cofactors with broad mRNP processing capabilities (e.g., CHTOP in 3’ end processing and ALYREF in splicing) (Viphakone *et al.*, 2019) known to compete for the limited number of UAP56 binding sites (Hautbergue et al., 2009; Heath *et al.*, 2016; Izumikawa et al., 2018). The tetramer also affords a greater surface area to maintain the structural integrity of long mRNA transcripts during mRNP processing and export. Through this process, TREX serves as a mRNA chaperone to prevent formation of DNA-RNA hybrid or R-loop structures that can promote genome instability (Luna *et al.*, 2019; Pérez-Calero *et al.*, 2020). Thus, the TREX tetramer helps ensure mRNP quality control and evasion of degradation by the nuclear exosome (Fan et al., 2017). The tetramer may also enable multiple TREX complexes to simultaneously bind several mRNP regions (Pühringer *et al.*, 2020) to facilitate compaction and/or protection of longer transcripts with elevated splicing.

Additional insights into dimer versus tetramer functions come from functional investigation of THO monomer components. For instance, THOC2 and THOC5 are required for the maintenance of ESC pluripotency, whereas TREX tetramer cofactors ALYREF and UAP56 impact differentiation (Wang et al., 2013). This phenotypic difference may reflect cell- and/or species-specific TREX dimer versus tetramer mRNP processing and export functions, with THO monomers being essential for viability and tetramer cofactors critical for differentiation and development. Enhanced mRNA processing efficiency provided by UAP56, ALYREF, CHTOP, and other export adaptors associated with the TREX tetramer are susceptible to *THOC6* pathogenic variants. Likewise, the diversity of expressed isoforms increases phylogenetically (yeast versus mammals), as well as during mammalian differentiation (Chen *et al.*, 2006; Mazin et al., 2021; Nagasaki *et al.*, 2005), especially in the brain, highlighting the vulnerability of neural development to *THOC6* loss. (Mauger *et al.*, 2016; Wang et al., 2008). Neuronal expression of long genes is imperative for proper neuronal differentiation and synaptogenesis (Zylka et al., 2015). ALYREF is predicted to interact with other UAP56/TREX-bound adaptors associated to the same mRNP (Pühringer *et al.*, 2020), suggesting that the tetramer facilitates increased mRNP compaction required for proper export, especially of longer mRNAs. It is possible that enhanced selectivity of mRNP processing and export co-evolved with TREX conformation, contributing to organismal complexity and distinguishing mammalian and yeast cells.

In addition to its prominent role in nuclear RNA export, TREX has been implicated in splicing functions that correlate to increased export efficiency of spliced mRNAs (Masuda *et al.*, 2005; Viphakone *et al.*, 2019). As such, a significant reduction in global polyA+ RNA nuclear export was predicted to result from loss of THOC6-mediated TREX function. However, in the absence of a global mRNA export defect in *THOC6*-affected hNPCs, alternative pathogenic mechanisms were investigated. Significant splicing changes implicate a pathogenic mechanism where THOC6-dependent disruption of TREX tetramer formation indirectly disrupts coordination of multiple steps of mRNP processing, including splicing, upstream of polyA+ mRNA packaging and export. This interpretation is supported by the diversity of mRNP processing functions attributed to tetramer-associated cofactors. UAP56 and ALYREF play important roles in mediating pre-mRNA splicing decisions (Shen et al., 2007; Viphakone *et al.*, 2019). This finding does not rule out the possibility that THOC6 plays a direct role in pre-mRNA splicing outside of mediating TREX core tetrameric assembly on the EJC. That THO member THOC5 can interact with unspliced transcripts (Chi *et al.*, 2013), and WD40-repeat domains facilitate splicing factor interactions with pre-mRNA are evidence in support of this possibility (Jin *et al.*, 2020).

Our findings implicating THOC6-dependent TREX tetramers as indirect facilitators of splicing by coordinating the mechanics of mRNP processing is also supported by enrichment of aberrant splicing events at weaker splice sites. Weak splice sites are most often utilized by transcripts during alternative splicing, and genes with elevated isoform diversity from alternative splicing are more susceptible to disruption of the overall integrity of mRNP processing in *THOC6*-affected hNPCs. Long genes that are highly expressed in the brain particularly rely on such infrastructure to ensure pro-neural gene expression. This may also be supported by the observed enrichment of retained introns in THOC6-deficient cells. Indeed, retained introns in unaffected tissues typically have weaker 5’ and 3’ splice sites compared to other splice junctions (Monteuuis et al., 2019). In addition, intron retention increases during mammalian differentiation (Braunschweig et al., 2014), again suggesting differentiated cells may be more susceptible to loss of THOC6-mediated TREX tetramer functions by miscoordination affecting splicing outcomes.

Given the conserved splicing and nuclear export functions in mammals, the phenotypic discrepancy between *Thoc6^fs/fs^* mouse embryonic lethality and human biallelic pathogenic *THOC6* variants is notable. Superficially, this finding suggests that humans are more tolerant to *THOC6* variants. Alternatively, this may represent differences in the pathogenic mechanisms of each variant or interspecies sensitivities to splicing defects. In addition, the downstream genetic mechanisms of *Thoc6* and *THOC6* variants are divergent. Human pathogenic *THOC6* variants reside in the WD40 repeat domains whereas the mouse *Thoc6^fs^* variant is located upstream of the first WD40 repeat domain. Nonsense *THOC6* variant transcripts are stable and readthrough permits limited THOC6 expression. Conversely, *Thoc6^fs^* transcripts are subject to NMD, impairing THOC6 expression. This suggests that minimal THOC6 expression may be sufficient for human embryogenesis. Splicing requirements may also account for interspecific phenotypic differences. While SE and RI events were the predominant splicing defects in both model systems, AS events affect more protein coding transcripts and lncRNAs in THOC6-affected hNPCs. Despite less AS events in *Thoc6^fs/fs^* cells, lethality in the mouse could be attributed to aberrant alternative splicing of specific transcripts important for mouse embryogenesis. Additionally, previous findings indicate that RI events account for a substantial portion of splicing variation in the primate prefrontal cortex, a trend that is most pronounced in humans (Mazin et al., 2018). Although intron retention is a known mechanism of mouse neuronal gene regulation by initiating RNA exosome-mediated degradation (Yap et al., 2012), it is possible that human cells are more tolerant than mouse cells to elevated intron retention. Further investigation of these interspecific differences is important for generating translationally relevant discoveries.

The number of ID genes that are mis-spliced in *THOC6* affected hNPCs relative to controls implicate shared underlying developmental mechanisms of ID pathology. However, developmental impact of individual defects on TIDS neuropathology is complicated by the compounding effects of constitutive THOC6 LOF models. In addition to trends shared with the mouse model, we show that biallelic THOC6 LOF is responsible for disruption of key TGF-β and Wnt signaling pathways via a mechanism that involves dysregulation of signaling components and lncRNAs resulting in delayed hNPC differentiation, prolonged retention of multipotency, and enhanced apoptosis. This is exemplified by intron retention and upregulation of *MEG3* in affected hNPCs. *MEG3* is linked to the regulation of TGF-β signaling and other EZH2 common target genes (Mondal *et al.*, 2015). Our findings suggest that RI events alter *MEG3* subcellular localization, expression, and downstream WNT signaling that increases multipotency and disrupts the balance of proliferation and differentiation in affected hNPCs. A shift towards cytoplasmic localization of lncRNAs has evolved in human cells, which is important for the maintenance of stem cell pluripotency (e.g., cytoplasmic *FAST* binds E3 ubiquitin ligase β-TrCP to block its interaction with β-catenin and enable activation of Wnt signaling) (Azam et al., 2019; Guo et al., 2020). Given the increased diversity of lncRNA functions in human developmental biology, mouse cells may be less tolerant to lncRNA dysregulation than human cells. In addition, *MEG3* is also upregulated by CREB (Zhao et al., 2006) whose target genes are affected in *Thoc5* conditional knockout mouse cortical neurons (Maeder *et al.*, 2018), potentially reflecting a shared mechanism of THO dysregulation in neural cells. While our analyses from mouse and human organoid models of *Thoc6/THOC6* disruption provide insight into the molecular pathology of early neural development, later analysis of synaptic physiology will be important to elucidate mechanisms of neuronal dysfunction in TIDS.

Altogether, our findings expand the TIDS clinical population and provide novel functional insight into the pathogenic mechanisms of biallelic LOF variants in *THOC6* using comparative mammalian model systems. Functional studies with THOC6 enable us to assess TREX tetramer function while retaining THO subcomplex formation, and our findings provide novel insight into TREX splicing functions separate from export. Future work is needed to dissect the direct and indirect effects of THOC6 loss and confirm endogenous tetramer disruption under native protein conditions. In addition, the well-known role of several TREX members in determination of polyadenylation site choice necessitates research focused on characterizing global aberrant alternative polyadenylation changes that could be contributing to dysregulation. Follow-up investigation at later-stage cortical organoids, and with use of unbiased single-cell RNAseq profiling, will allow for more detailed assessment of the developmental consequences of observed defects for cortical lamination and cell type composition. Lastly, it will be important to determine if alterations in mRNA processing/export also underlie synaptic defects—a morphological basis of ID, and a prominent clinical feature shared between *THOC2* and *THOC6*-associated neurodevelopmental disorders.

## ACKNOWLEDGEMENTS

The authors would like to especially thank all the family members who participated in this study and gave permission to use genetic and clinical information. We thank Stephanie Moon for guidance on mRNA stability assay, Sumin Kim for reagents and protocols for oligo-dT FISH, and the University of Michigan Advanced Genomics, Transgenic Animal Model, and Microscopy Cores for sequencing and equipment. This research was supported by NSF Doctoral Dissertation Research Improvement Grant 1919671: F055138, Wenner-Gren Foundation for Anthropological Research N02820: 9967, Leakey Foundation Dissertation Fieldwork Grant, University of Michigan Rackham Candidate Research Grant, the Joan B. Kessler Award, and the University of Michigan Pandemic Research Recovery Award to E.A.W, as well as the National Institutes of Health award numbers R01AWD010411 to S.L.B, A.S., and K.M.G, R00HD099403 to A.E.S, F31NS122207 and T32NS077888 to G.R.L, and T32GM008056 to K.J.

## AUTHOR CONTRIBUTIONS

A.E.S. generated the mouse model and human cell lines. E.A.W. performed human cell culture and organoid experiments and all bioinformatics analyses. G.R.L., K.J., B.B., S.L.R, C.D.P, and A.E.S. performed mouse experiments. D.P., S.L.R., and C.D.P. contributed to immunohistochemistry analyses. E.A.W. and G.R.L. performed the molecular biology experiments. E.A.W. created the figures. E.A.W., G.L., S.L.B, and A.E.S. wrote the manuscript. A.E.S., S.L.B, A.S., S.B., S.M., R.P., M.H, R.J.H, E.K, A.O, J.D., A.K., K.C., E.J.P., R.J.L., R.R.L., T.L.E., C.G., K.M.G., K.B., and A.S. contributed to the clinical work.

## METHODS

### Human subjects

All subjects or parents/guardians provided informed consent and were enrolled in institutional review board-approved research studies. In all cases, the procedures followed were in accordance with the ethical standards of the respective institution’s committee on human research and were in keeping with international standards. Probands 1-3 and 5 were identified through GeneMatcher (Sobreira et al., 2015). Details for all subjects are provided in Table S1.

### Animal models

All mice were maintained according with the National Institutes of Health Guidelines for the Care and Use of Laboratory Animals and were approved by the Case Western Reserve Institutional Animal Care and Use Committee. CRISPR genome editing was performed in the University of California, San Diego Transgenic and Knockout Mouse Core. C57BL/6JN hybrid mice (Jackson Laboratory, 005304) were used for CRISPR editing of the *Thoc6* locus. Founder mice with the *Thoc6^fs/+^* allele were intercrossed with C57BL/6JN mice (Jackson Laboratory, 005304) for line maintenance. All *ex vivo* analyses were performed on tissue collected from mice of both sexes at embryonic day (E) 8.5-10.5. Sex-dependent differences were not assessed.

Litters were genotyped by allele-specific polymerase chain reaction (AS-PCR). Genomic DNA was prepared from mouse tissue samples as previously described (Truett et al., 2000). AS-PCR for each allele was assembled using the standard GoTaq DNA polymerase (Promega) protocol. Reaction conditions were executed as recommended by the manufacturer. Primers and sgRNA sequences are provided in Table S3.

### Human ESC/iPSC culture

Human ESC and iPSC lines were cultured using feeder free conditions on Matrigel (Corning) with mTeSR-1 (STEMCELL Technologies). Lines used in this manuscript include: H9 (*THOC6^+/+^*, 46XX, ESCs, WA09, WiCell), AS0035 (*THOC6^+/+^*, 46XX, iPSCs, New York Stem Cell Foundation (NYSCF) Diabetes iPSC Panel), AS0041 (*THOC6^+/+^*, 46XY, iPSCs, NYSCF Diabetes iPSC Panel), KMC6002 (*THOC6^E188K/+^*, 46XY, iPSCs), KMC6003 (*THOC6^E188K/E188K^*, 46XY, iPSCs), KMC7001 (*THOC6^W100*/+^*, 46XY, iPSCs), KMC7002 (*THOC6^W100*/W100*^*, 46XY, iPSCs). Passaging was performed using mTeSR-1 supplemented 1 nM ROCK inhibitor (BD Biosciences 562822) to prevent differentiation. Both manual and chemical dissociation with Versene (Gibco 15040066) were performed for splitting. Sanger sequencing validation of genotypes (Figure S1B and S1C) (primers listed in Table S3) and CNV microarray analysis (Illumina Bead Array, analysis with Genome Studio v2.0) were performed on all lines to ensure no pathogenic changes were acquired during culturing. No further validation of iPSCs lines was performed in our lab.

## METHOD DETAILS

### Whole exome sequencing and analysis

Exome libraries from genomic DNA of all BBIS-affected probands were prepared and captured with the Agilent SureSelectXT Human All Exon 50Mb Kit for Probands 1 & 4-7 and Individual 5, the Agilent SureSelectXT Clinical Research Exome kit for Proband 3, and the TrueSeq Rapid Exome Kit for Proband 2. Further, exome libraries were sequenced on an Illumina HiSeq or NextSeq instrument as described previously (Srivastava et al., 2018).

Reads were aligned to the human reference genome NCBI builds 37 (GRCh37) and 38 (GRCh38) and 38 using Burrows-Wheeler Aligner (BWA) (Li and Durbin, 2009). Variant calling of single nucleotide variants (SNVs) and copy number variants (CNVs) was performed using GATK (Van der Auwera et al., 2013), VEP, and CoNIFER (Krumm et al., 2012). Average depth of coverage was calculated across all targeted regions. The data were filtered and annotated from the canonical *THOC6* transcript (ENST00000326266.8 and ENSP00000326531.8) using in-house bioinformatics software (Müller et al., 2017). Variants were also filtered against public databases including the 1000 Genomes Project phase 311, Exome Aggregation Consortium (ExAC) v.0.3.1, Genome Aggregate Database (gnomAD), National Heart, Lung, and Blood Institute Exome Sequencing Project Exome Variant Server (ESP6500SI-V2). Those with a minor allele frequency >3.3% were excluded. Additionally, variants flagged as low quality or putative false positives (Phred quality score 14; 15 < 20, low quality by depth <20) were excluded from the analysis. Variants in genes known to be associated with ID were selected and prioritized based on predicted pathogenicity.

### Sanger sequencing

All variants discovered by WES were confirmed with Sanger sequencing of *THOC6* for each individual and respective family members who submitted samples except Proband 1 where high-coverage WES of *THOC6* in the proband and parents was deemed sufficient to report without Sanger confirmation. Chromatograms were analyzed using NextGENe, Sequencer, and Geneious Prime Software (v.2022.1.1).

### Cerebral organoid generation

Telencephalic cerebral organoids were generated based on previously published protocols (Lancaster et al., 2013), with few modifications to start with low cell density in order to generate smaller and more consistent embryoid bodies (EBs). Briefly, human and chimpanzee iPSCs were passaged into 96-well V-shaped bottom ultra-low attachment cell culture plates (PrimeSurface® 3D culture, MS-9096VZ) to achieve a starting cell density of 600-1,000 cells per well in 30 µl of mTesR^TM^1 with 1 nM ROCK inhibitor. After 36 hours, 150 µl of N-2/SMAD inhibition media (cocktail of 1X N-2 supplement (Invitrogen 17502048), 2 μM A-83-01 inhibitor (Tocris Bioscience 2939), and 1 mM dorsomorphin (Tocris Bioscience 309350) in DMEM-F12 (Gibco 11330032)) was added for neural induction. On day 7, EBs were transferred to Matrigel-coated plates to enrich for neural rosettes at a density of 20-30 EBs per well of a 6-well plate, and media was changed to neural differentiation media (0.5X N-2 supplement, 0.5X B-27 supplement (Invitrogen 17504044) with 20 pg/μl bFGF and 1mM dorsomorphin inhibitor in DMEM/F-12). For organoid differentiation EBs were outlined on day 14 using a pipet tip and uplifted carefully with a cell scraper to minimize organoid fusion and tissue ripping. Media was changed once more to N-2/B-27 with bFGF only and plates with uplifted organoids were placed on a shaker in the incubator set at a rotation speed of 90. On day 14, media was changed once more to N-2/B-27 with bFGF only. Prior to day 14, media changes were performed every 48 hours. After day 14, daily media changes were performed until collection. For monolayer NPC differentiation, neural rosettes were scored and uplifted on day 14, dissociated in Accutase (Gibco A1110501), and re-plated on poly-L-ornithine (PLO)/Laminin-coated plates for NPC expansion, selection, and passaging. 15 μg/mL PLO (Sigma-Aldrich P4957) diluted in DPBS (Gibco 14040-133); 10 μg/mL laminin (Sigma-Aldrich L2020) diluted in DMEM/F-12.

### Western blot analysis and immunoprecipitation

ESCs, iPSCs, and NPCs used for western blot analysis were pelleted and lysed in RIPA buffer supplemented with 1:50 protease inhibitor cocktail (Sigma-Aldrich P8340) and 1:100 phosphatase inhibitor cocktail 3 (Sigma-Aldrich P0044) using mortar and pestle coupled with end-over-end rotation for 30 minutes to 1 hr at 4°C. Protein concentration was quantified by BCA (Thermo Scientific Pierce A53227). Lysis samples were then incubated at a 1:3 ratio with 4x Laemmli sample buffer (Bio-Rad) supplemented with 10% BME and incubated at 95°C on a heat block for 5 minutes for denaturation. For co-immunoprecipitation, primary antibodies anti-THOC5 and anti-THOC6 (1:50 dilution in 1x PBS with Tween-20) were incubated overnight at 4°C with Dynabeads Protein G (Invitrogen, 10003D). Beads were washed and cell lysis (35 μg of protein) was added for incubation overnight at 4°C with rotation. IP samples were prepared according to manufacturer’s instructions with elution in Laemmli sample buffer with 10% BME. For promotion of readthrough of premature termination codons, ataluren (eMolecules NC1485023) was dissolved in DMSO added to ESC/iPSC media at a final concentration of 30 μM for 48 hours as previously described (Roy et al., 2016). Protein was then extracted as described above.

Samples were loaded into 4-20% SDS-polyacrylamide gels (Bio-Rad) and proteins were separated by electrophoresis at 30V for ∼4 hours room temperature. Separated proteins were then transferred to PVDF membranes (Millipore) overnight using a wet transfer system (Bio-Rad) at 4°C. For immunoblotting, membranes were incubated in 5% milk blocking buffer (1x TBS-T) followed by primary antibody incubation overnight at 4°C with rotation. Membranes were washed 3 times for 5 minutes in 1x TBS-T and then incubated with secondary antibodies for 1-2 hours at room temperature. Membranes underwent final washes before developing using West Femto Substrate (ThermoFisher 34095) with film exposure. Primary antibodies used: mouse anti-THOC6 (1:1000, Abnova H00079228-A01), rabbit anti-THOC1 (1:200, Bethyl Laboratories A302-839A), rabbit anti-THOC2 (1:200, Bethyl Laboratories A303-630A), rabbit anti-THOC5 (1:200, Bethyl Laboratories A302-120A), mouse anti-ALYREF (1:200, Sigma Aldrich A9979), rabbit anti-CHTOP (1:200, Invitrogen PA5-55929), mouse anti-β-actin (1:250, Abcam ab6276). Secondary antibodies used: donkey anti-rabbit HRP-conjugated (1:5000, Cytiva NA9340V) and goat anti-mouse HRP-conjugated (1:1000, Invitrogen 32430).

### Immunofluorescence and Single-molecule Fluorescence *in situ* Hybridization

Human NPCs were fixed in 4% paraformaldehyde (PFA) for 20 minutes. Human cortical organoids and mouse embryos were fixed in 4% PFA for 24 hours at 4°C, cryoprotected in 15% and 30% sucrose in 1x DPBS for 24 hours at 4°C, then embedded in OCT with quick freezing in −50°C 2-methylbutane, followed by cryosectioning for immunostaining. Mouse embryos were sectioned at 13 µm and organoids at 16 µm. Samples for immunostaining were incubated for 1 hour with blocking buffer (5% NDS (Jackson ImmunoResearch) 0.1% Triton X-100, 5% BSA) at room temperature, then overnight with primary antibodies diluted in blocking buffer at 4°C, and for 1-2 hours in secondary dilution at room temperature. Washes performed in PBS. For nuclear staining, samples were incubated at room temperature for 10 minutes in Hoescht or DAPI (1:1000 dilution in PBS) prior to final washes. For EdU labeling detection, the Click-IT EdU imaging kit (Invitrogen C10337) was used according to the manufacturer’s instructions. After incubation with the Click-IT reaction cocktail, sections were blocked and immunostained as described above. Some antibodies required antigen retrieval via incubation in heated 10 mM sodium citrate solution (95-100°C) for 20 minutes prior to immunostaining. Primary antibodies used: mouse anti-PAX6 (1:250, Abcam MA-109), rabbit anti-KI67 (1:200, Abcam ab16667), rat anti-PH3 (1:250, Abcam ab10543), rabbit anti-cleaved caspase3 (1:100-1:400, Cell Signaling 9661), mouse anti-N-Cadherin (BD Biosciences 610920), goat anti-DCX (1:400, Santa Cruz Biotechnology A1313), rat anti-CTIP2 (1:500, Abcam ab18465), rabbit anti-PAX6 (1:100, BioLegend PRB-278P) and goat anti-SOX1 (1:100, R&D Biosystems AF3369). AlexaFluor-conjugated secondaries used: donkey anti-mouse 647 (1:400, Invitrogen A31571), donkey anti-rat 555 (1:400, Invitrogen A48270), and donkey anti-rabbit 488 (1:400, Invitrogen A21206).

Embryos and organoids used for RNA Fluorescence *in situ* Hybridization (FISH) were fixed and cryoprotected as indicated above using RNAse-free PBS. RNAse-zap treatment of sectioning equipment was performed prior to cryosectioning. NPCs for RNA FISH were fixed in RNAse-free 4% PFA then permeabilized in PBS-TritonX (0.1%) for 15 minutes. Hybridizations were then performed overnight at 37°C with a final concentration of 2 ng/µl of Cy3-conjugated oligo-dT(30-mer) probe, *MALAT1* (Quasar-670, Stellaris VSMF-2211-5), and/or *MEG3* (Quasar-570, Stellaris VSMF-20346-5). Saline-sodium citrate washes were performed before and after hybridization, followed by nuclear staining with RNAse-free Hoescht-PBS wash (1:1000 dilution) and final wash in RNAse-free PBS.

Glass covers were mounted onto all slides with Prolong Gold (Molecular Probes S36972) and incubated for 24 hours at room temperature prior to imaging. Imaging was performed with a Nikon A1ss inverted confocal microscope using NIS-Elements Advanced Research software. Image analysis was performed using Fiji (ImageJ) software (Schindelin et al., 2012). For oligo-dT FISH, Z-series images were taken every 0.2 μm across entire width of cells for each genotype using same laser intensity settings and collapsed by max intensity using Z project tool in Fiji for quantification of nuclear and cytoplasmic fractions of polyA intensity by automated quantitation with CellProfiler (v4.2.1). Hoechst signal was used to segment nuclei and the oligo-dT signal to segment cell body. Three differentiation replicates per genotype. 3D surface plots were made in Fiji.

### WGA inhibition of nuclear export

Confluent NPCs were incubated with digitonin at 30 μg/mL diluted in DMSO and WGA conjugated to Alexa Fluor 488 (Invitrogen, W11261) at 5 μg/mL diluted in DPBS for 5 minutes, as previously described (Mor *et al.*, 2010). Cells were washed to remove digitonin and WGA only was added to media at 5 μg/mL for 1 hour. Control NPCs were only treated with digitonin. Cells were fixed and prepped for oligo-dT FISH as described above.

### RNA sequencing and bioinformatics analysis

Total RNA was extracted from cultured hNPCs (two biological replicates per genotype) using TRIzol Reagent (Invitrogen 15596026) followed by DNAse column treatment using PureLink RNA extraction kit (Invitrogen 12183018A). Total RNA from dissected E9.5 mouse forebrain tissue (three biological replicates per genotype) was extracted using Picopure RNA isolation kit (Applied Biosystems KIT0204) according to manufacturer’s recommendations. hNPC and E9.5 mouse forebrain RNA samples were ribo-depleted followed by 151 bp paired-end sequencing on the Illumina NovaSeq 300 cycle, ∼20-30 million reads per sample. Library preparation and sequencing was conducted by the Advanced Genomics Core (AGC) at the University of Michigan. ERCC spike-ins (Invitrogen 4456740) were added for sequencing controls at starting concentrations according to the manufacturer’s instructions. FASTQ files were trimmed with Cutadapt v4.1 using default parameters (Martin, 2011). Read quality was assessed by FASTQC v0.11.9 (Wingett and Andrews, 2018). MultiQC v1.7 (Ewels et al., 2016) was used to visualize FASTQC outputs and compare samples. ERCC spike-in FASTA and GTF annotation files were merged with human GRCh38.p13 reference genome FASTA with GTF release 39 or mouse GRCm39 reference genome FASTA with GTF release M28. FASTQ reads were then mapped to merged files using STAR alignment with parameter ‘--outSAMtype BAM SortedByCoordinate’ (Dobin et al., 2013). Count analysis was performed on sorted BAM files using RSEM with paired-end alignment specified (Li and Dewey, 2011). Differential expression analysis was carried out using DESeq2 v1.34.0 (Love et al., 2014) in R v4.1.2 (Team, 2018). ERCC spike-in counts were used to estimate size factors for each sample for DESEq2 analysis. Genes were considered dysregulated if FDR < 0.05 and fold-change > 2 or < −2. Volcano and PCA plots were made using ggplot2 and pcaExplorer packages in R.

Alternative splicing analysis was performed on sorted BAM files using rMATS v4.1.2 (Park et al., 2013) with the following parameters: ‘-t paired --readLength 150 --variable-read-length --nthread 4’ (Shen *et al.*, 2014). AS events were called if FDR < 0.05 and ΔPSI > 10%. Events with less than 5 average reads were filtered out using the MASER package in R (Veiga, 2022). To calculate splice site strength at 5’ and 3’ splice sites in AS transcripts identified by rMATS, maximum entropy modeling was carried out using MaxEntScan (Yeo and Burge, 2004). The required input is a 9-mer sequence at 5’ splice sites (3 bases in exon and 6 bases in downstream intron) and a 23-mer at 3’ splice site (20 bases of intron and 3 bases of downstream exon). Scores were plotted in GraphPad Prism (v9.3.1).

DAVID (david.ncifcrf.gov/tools) (Subramanian et al., 2005) and Metascape (metascape.org) (Zhou et al., 2019) analyses were performed to identify enriched biological pathways based on Benjamini-Hochberg multiple hypothesis corrections of the *p*-values. To identify potential transcription factors responsible for expression differences, Gene Set Enrichment Analysis (GSEA v4.2.3) against the MSigDB transcription factor motif gene set (c4.tftv7.5.1.symbols.gmt) and ChIP-X Enrichment Analysis v3 (ChEA3) were performed (Keenan et al., 2019). Ensembl BioMart tool (http://useast.ensembl.org/biomart) was used to obtain coding sequence length, transcript number per gene, gene type, and sequences for AS events. The GeneOverlap v1.32 R package was used to identify overlapping DE and AS hits between affected genotypes. Primary and candidate syndromic ID genes were obtained from the SysID database (https://www.sysid.dbmr.unibe.ch).

### qRT-PCR and mRNA half-live analysis

Reverse transcription for cDNA synthesis was performed using 1 μg of total RNA with Superscript III first-strand synthesis kit (Invitrogen 18080051) according to manufacturer’s instructions. For validation of AS events, standard PCR was performed as described above. Abundances of *THOC6*, *FOS*, *GAPDH*, *MEG3*, *MEG8*, *ESRG*, and *NEAT1* mRNA was determined by quantitative real-time PCR (qPCR) using the Applied Biosystems 7500 system with 7500 Software v2.3 and Radiant Green 2X qPCR Mix Lo-ROX 2X qPCR Mix (Alkali Scientific inc., QS1020) according to manufacturer’s instructions. Cycler parameters used: cDNA activation (1 cycle at 95°C for 2 minutes), denaturation (40 cycles 95°C for 5 seconds) and annealing/extension (40 cycles at 60°C for 30 seconds). The ΔΔCt method was used to analyze data with *GAPDH* as a reference gene. ΔΔCt values obtained by subtracting mean *THOC6^+/+^* ΔCt values for each sample. Data shown represent mean values of three qPCR technical replicates per sample for three biological replicates per genotype (independent differentiations for NPCs). Melt curve analysis was performed on all primers to ensure temperature peaks at ∼80-90°C. *GAPDH* and *FOS* primer sequences were obtained from (Moon *et al.*, 2012). *NEAT1* was obtained from (Cui *et al.*, 2019) and *MEG3* was obtained from (Mondal *et al.*, 2015). All others were designed using NCBI primer blast. Primer sequences provided in Table S3.

mRNA decay analysis was performed using transcription inhibition by Actinomycin D (ActD) based on (Moon *et al.*, 2012). Human ESCs/iPSCs were first passaged into five 12-well plates. Each plate had the following lines: *THOC6^+/+^* (H9 ESCs), *THOC6^+/+^* (AS0041 iPSCs), *THOC6^W100*/W100*^*, *THOC6^W100*/+^*, *THOC6^E188K/E188K^, THOC6^E188K/+^*. Once confluent, ActD was added to media of all four plates at 10 μg/mL (Sigma-Aldrich A9415). After 30 minutes, media was removed from one plate and 1 mL of TRIzol Reagent was added directly to each well (*t* = 0). Cells were uplifted in TRIzol by pipetting and transferred to a fresh tube. Tubes were immediately frozen in TRIzol at −80°C. This was repeated every 30 minutes to obtain the following time points 30 minutes post-ActD treatment: *t* = 0.5, 1, 1.5, and 2 hrs. Extractions were performed in batches per time point based on protocol described above. Standard curve analysis was performed to validate primers (Figure S1C). This experiment was repeated to capture longer decay window using the following time points: *t* = 0, 2, 4, 8 (Figure S1D). ΔΔCt values obtained by subtracting mean *t* = 0 ΔCt for each genotype. Abundances of *THOC6*, *GAPDH*, and *FOS* (positive control for rapid decay) mRNA was determined.

## QUANTIFICATION AND STATISTICAL ANALYSIS

Statistical significance of all quantifications from microscopy images, western blot images, gel images, and qRT-PCR abundances was tested using a student’s two-tailed *t*-test and data was plotted using GraphPad Prism (v9.3.1) as mean ±SEM or mean ±SD, as specified in figure legends. Simple linear regression was performed in qRT-PCR standard curve analysis, organoid growth curves, and intron retention analysis. Statistical significance of gene overlaps were tested using Fisher’s exact test via GeneOverlap package function testGeneOverlap() in R. Benjamini-Hochberg multiple hypothesis corrections were performed in pathway enrichment analyses.

## REFERENCES

Accogli, A., Scala, M., Calcagno, A., Castello, R., Torella, A., Musacchia, F., Allegri, A.M.E., Mancardi, M.M., Maghnie, M., Severino, M., et al. (2018). Novel CNS malformations and skeletal anomalies in a patient with Beaulieu-boycott-Innes syndrome. Am J Med Genet A 176, 2835–2840. 10.1002/ajmg.a.40534.

Amos, J.S., Huang, L., Thevenon, J., Kariminedjad, A., Beaulieu, C.L., Masurel-Paulet, A., Najmabadi, H., Fattahi, Z., Beheshtian, M., Tonekaboni, S.H., et al. (2017). Autosomal recessive mutations in THOC6 cause intellectual disability: syndrome delineation requiring forward and reverse phenotyping. Clin Genet 91, 92–99. 10.1111/cge.12793.

Anazi, S., Alshammari, M., Moneis, D., Abouelhoda, M., Ibrahim, N., and Alkuraya, F.S. (2016). Confirming the candidacy of THOC6 in the etiology of intellectual disability. Am J Med Genet A 170A, 1367-1369. 10.1002/ajmg.a.37549.

Anazi, S., Maddirevula, S., Faqeih, E., Alsedairy, H., Alzahrani, F., Shamseldin, H.E., Patel, N., Hashem, M., Ibrahim, N., Abdulwahab, F., et al. (2017). Clinical genomics expands the morbid genome of intellectual disability and offers a high diagnostic yield. Mol Psychiatry 22, 615–624. 10.1038/mp.2016.113.

Andrews, M.G., Subramanian, L., and Kriegstein, A.R. (2020). mTOR signaling regulates the morphology and migration of outer radial glia in developing human cortex. Elife 9. 10.7554/eLife.58737.

Azam, S., Hou, S., Zhu, B., Wang, W., Hao, T., Bu, X., Khan, M., and Lei, H. (2019). Nuclear retention element recruits U1 snRNP components to restrain spliced lncRNAs in the nucleus. RNA Biol 16, 1001–1009. 10.1080/15476286.2019.1620061.

Bahar Halpern, K., Caspi, I., Lemze, D., Levy, M., Landen, S., Elinav, E., Ulitsky, I., and Itzkovitz, S. (2015). Nuclear Retention of mRNA in Mammalian Tissues. Cell Rep 13, 2653–2662. 10.1016/j.celrep.2015.11.036.

Beaulieu, C.L., Huang, L., Innes, A.M., Akimenko, M.A., Puffenberger, E.G., Schwartz, C., Jerry, P., Ober, C., Hegele, R.A., McLeod, D.R., et al. (2013). Intellectual disability associated with a homozygous missense mutation in THOC6. Orphanet J Rare Dis 8, 62. 10.1186/1750-1172-8-62.

Boycott, K.M., Beaulieu, C., Puffenberger, E.G., McLeod, D.R., Parboosingh, J.S., and Innes, A.M. (2010). A novel autosomal recessive malformation syndrome associated with developmental delay and distinctive facies maps to 16ptel in the Hutterite population. Am J Med Genet A 152A, 1349-1356. 10.1002/ajmg.a.33379.

Braunschweig, U., Barbosa-Morais, N.L., Pan, Q., Nachman, E.N., Alipanahi, B., Gonatopoulos-Pournatzis, T., Frey, B., Irimia, M., and Blencowe, B.J. (2014). Widespread intron retention in mammals functionally tunes transcriptomes. Genome Res 24, 1774–1786. 10.1101/gr.177790.114.

Casey, J., Jenkinson, A., Magee, A., Ennis, S., Monavari, A., Green, A., Lynch, S.A., Crushell, E., and Hughes, J. (2016). Beaulieu-Boycott-Innes syndrome: an intellectual disability syndrome with characteristic facies. Clin Dysmorphol 25, 146–151. 10.1097/MCD.0000000000000134.

Chai, G., Webb, A., Li, C., Antaki, D., Lee, S., Breuss, M.W., Lang, N., Stanley, V., Anzenberg, P., Yang, X., et al. (2021). Mutations in Spliceosomal Genes PPIL1 and PRP17 Cause Neurodegenerative Pontocerebellar Hypoplasia with Microcephaly. Neuron 109, 241–256.e249. 10.1016/j.neuron.2020.10.035.

Chang, C.T., Hautbergue, G.M., Walsh, M.J., Viphakone, N., van Dijk, T.B., Philipsen, S., and Wilson, S.A. (2013). Chtop is a component of the dynamic TREX mRNA export complex. EMBO J 32, 473–486. 10.1038/emboj.2012.342.

Chen, F.C., Chen, C.J., Ho, J.Y., and Chuang, T.J. (2006). Identification and evolutionary analysis of novel exons and alternative splicing events using cross-species EST-to-genome comparisons in human, mouse and rat. BMC Bioinformatics 7, 136. 10.1186/1471-2105-7-136.

Cheng, H., Dufu, K., Lee, C.S., Hsu, J.L., Dias, A., and Reed, R. (2006). Human mRNA export machinery recruited to the 5’ end of mRNA. Cell 127, 1389–1400. 10.1016/j.cell.2006.10.044.

Chervitz, S.A., Hester, E.T., Ball, C.A., Dolinski, K., Dwight, S.S., Harris, M.A., Juvik, G., Malekian, A., Roberts, S., Roe, T., et al. (1999). Using the Saccharomyces Genome Database (SGD) for analysis of protein similarities and structure. Nucleic Acids Res 27, 74–78. 10.1093/nar/27.1.74.

Chi, B., Wang, Q., Wu, G., Tan, M., Wang, L., Shi, M., Chang, X., and Cheng, H. (2013). Aly and THO are required for assembly of the human TREX complex and association of TREX components with the spliced mRNA. Nucleic Acids Res 41, 1294–1306. 10.1093/nar/gks1188.

Chávez, S., Beilharz, T., Rondón, A.G., Erdjument-Bromage, H., Tempst, P., Svejstrup, J.Q., Lithgow, T., and Aguilera, A. (2000). A protein complex containing Tho2, Hpr1, Mft1 and a novel protein, Thp2, connects transcription elongation with mitotic recombination in Saccharomyces cerevisiae. EMBO J 19, 5824-5834. 10.1093/emboj/19.21.5824.

Cui, Y., Yin, Y., Xiao, Z., Zhao, Y., Chen, B., Yang, B., Xu, B., Song, H., Zou, Y., Ma, X., and Dai, J. (2019). LncRNA Neat1 mediates miR-124-induced activation of Wnt/β-catenin signaling in spinal cord neural progenitor cells. Stem Cell Res Ther 10, 400. 10.1186/s13287-019-1487-3.

Dias, A.P., Dufu, K., Lei, H., and Reed, R. (2010). A role for TREX components in the release of spliced mRNA from nuclear speckle domains. Nat Commun 1, 97. 10.1038/ncomms1103.

Dobin, A., Davis, C.A., Schlesinger, F., Drenkow, J., Zaleski, C., Jha, S., Batut, P., Chaisson, M., and Gingeras, T.R. (2013). STAR: ultrafast universal RNA-seq aligner. Bioinformatics 29, 15–21. 10.1093/bioinformatics/bts635.

Dufu, K., Livingstone, M.J., Seebacher, J., Gygi, S.P., Wilson, S.A., and Reed, R. (2010). ATP is required for interactions between UAP56 and two conserved mRNA export proteins, Aly and CIP29, to assemble the TREX complex. Genes Dev 24, 2043-2053. 10.1101/gad.1898610.

Ellis, J.D., Barrios-Rodiles, M., Colak, R., Irimia, M., Kim, T., Calarco, J.A., Wang, X., Pan, Q., O’Hanlon, D., Kim, P.M., et al. (2012). Tissue-specific alternative splicing remodels protein-protein interaction networks. Mol Cell 46, 884–892. 10.1016/j.molcel.2012.05.037.

Ewels, P., Magnusson, M., Lundin, S., and Käller, M. (2016). MultiQC: summarize analysis results for multiple tools and samples in a single report. Bioinformatics 32, 3047–3048. 10.1093/bioinformatics/btw354.

Gieldon, L., Mackenroth, L., Kahlert, A.K., Lemke, J.R., Porrmann, J., Schallner, J., von der Hagen, M., Markus, S., Weidensee, S., Novotna, B., et al. (2018). Diagnostic value of partial exome sequencing in developmental disorders. PLoS One 13, e0201041. 10.1371/journal.pone.0201041.

Greijer, A.E., and van der Wall, E. (2004). The role of hypoxia inducible factor 1 (HIF-1) in hypoxia induced apoptosis. J Clin Pathol 57, 1009–1014. 10.1136/jcp.2003.015032.

Gromadzka, A.M., Steckelberg, A.L., Singh, K.K., Hofmann, K., and Gehring, N.H. (2016). A short conserved motif in ALYREF directs cap- and EJC-dependent assembly of export complexes on spliced mRNAs. Nucleic Acids Res 44, 2348–2361. 10.1093/nar/gkw009.

Guo, C.J., Ma, X.K., Xing, Y.H., Zheng, C.C., Xu, Y.F., Shan, L., Zhang, J., Wang, S., Wang, Y., Carmichael, G.G., et al. (2020). Distinct Processing of lncRNAs Contributes to Non-conserved Functions in Stem Cells. Cell 181, 621–636.e622. 10.1016/j.cell.2020.03.006.

Gupta, N., Yadav, S., Gurramkonda, V.B., Vl, R., Sg, T., and Kabra, M. (2020). First report of THOC6 related intellectual disability (Beaulieu Boycott Innes syndrome) in two siblings from India. Eur J Med Genet 63, 103742. 10.1016/j.ejmg.2019.103742.

Guria, A., Tran, D.D., Ramachandran, S., Koch, A., El Bounkari, O., Dutta, P., Hauser, H., and Tamura, T. (2011). Identification of mRNAs that are spliced but not exported to the cytoplasm in the absence of THOC5 in mouse embryo fibroblasts. RNA 17, 1048–1056. 10.1261/rna.2607011.

Harrison-Uy, S.J., and Pleasure, S.J. (2012). Wnt signaling and forebrain development. Cold Spring Harb Perspect Biol 4, a008094. 10.1101/cshperspect.a008094.

Hassanvand Amouzadeh, M., Akhavan Sepahi, M., and Abasi, E. (2020). Proteinuria in Two Sisters with Beaulieu-Boycott-Innes Syndrome, A Case Report. Iran J Kidney Dis 14, 312–314.

Hautbergue, G.M., Hung, M.L., Golovanov, A.P., Lian, L.Y., and Wilson, S.A. (2008). Mutually exclusive interactions drive handover of mRNA from export adaptors to TAP. Proc Natl Acad Sci U S A 105, 5154–5159. 10.1073/pnas.0709167105.

Hautbergue, G.M., Hung, M.L., Walsh, M.J., Snijders, A.P., Chang, C.T., Jones, R., Ponting, C.P., Dickman, M.J., and Wilson, S.A. (2009). UIF, a New mRNA export adaptor that works together with REF/ALY, requires FACT for recruitment to mRNA. Curr Biol 19, 1918–1924. 10.1016/j.cub.2009.09.041.

Heath, C.G., Viphakone, N., and Wilson, S.A. (2016). The role of TREX in gene expression and disease. The Biochemical journal 473, 2911–2935. 10.1042/BCJ20160010.

Izumikawa, K., Ishikawa, H., Simpson, R.J., and Takahashi, N. (2018). Modulating the expression of Chtop, a versatile regulator of gene-specific transcription and mRNA export. RNA Biol 15, 849–855. 10.1080/15476286.2018.1465795.

Jalali, A., Bassuk, A.G., Kan, L., Israsena, N., Mukhopadhyay, A., McGuire, T., and Kessler, J.A. (2011). HeyL promotes neuronal differentiation of neural progenitor cells. J Neurosci Res 89, 299–309. 10.1002/jnr.22562.

Jimeno, S., and Aguilera, A. (2010). The THO complex as a key mRNP biogenesis factor in development and cell differentiation. J Biol 9, 6. 10.1186/jbiol217.

Jimeno, S., Rondón, A.G., Luna, R., and Aguilera, A. (2002). The yeast THO complex and mRNA export factors link RNA metabolism with transcription and genome instability. EMBO J 21, 3526–3535. 10.1093/emboj/cdf335.

Jin, L., Chen, Y., Crossman, D.K., Datta, A., Vu, T., Mobley, J.A., Basu, M.K., Scarduzio, M., Wang, H., Chang, C., and Datta, P.K. (2020). STRAP regulates alternative splicing fidelity during lineage commitment of mouse embryonic stem cells. Nat Commun 11, 5941. 10.1038/s41467-020-19698-6.

Juneau, K., Miranda, M., Hillenmeyer, M.E., Nislow, C., and Davis, R.W. (2006). Introns regulate RNA and protein abundance in yeast. Genetics 174, 511–518. 10.1534/genetics.106.058560.

Katahira, J., Inoue, H., Hurt, E., and Yoneda, Y. (2009). Adaptor Aly and co-adaptor Thoc5 function in the Tap-p15-mediated nuclear export of HSP70 mRNA. EMBO J 28, 556–567. 10.1038/emboj.2009.5.

Katahira, J., Okuzaki, D., Inoue, H., Yoneda, Y., Maehara, K., and Ohkawa, Y. (2013). Human TREX component Thoc5 affects alternative polyadenylation site choice by recruiting mammalian cleavage factor I. Nucleic Acids Res 41, 7060–7072. 10.1093/nar/gkt414.

Keenan, A.B., Torre, D., Lachmann, A., Leong, A.K., Wojciechowicz, M.L., Utti, V., Jagodnik, K.M., Kropiwnicki, E., Wang, Z., and Ma’ayan, A. (2019). ChEA3: transcription factor enrichment analysis by orthogonal omics integration. Nucleic Acids Res 47, W212–W224. 10.1093/nar/gkz446.

Kiraz, A., Tubaş, F., and Seber, T. (2022). A truncating variant in the THOC6 gene with new findings in a patient with Beaulieu-Boycott-Innes syndrome. Am J Med Genet A 188, 1568–1571. 10.1002/ajmg.a.62667.

Kochinke, K., Zweier, C., Nijhof, B., Fenckova, M., Cizek, P., Honti, F., Keerthikumar, S., Oortveld, M.A., Kleefstra, T., Kramer, J.M., et al. (2016). Systematic Phenomics Analysis Deconvolutes Genes Mutated in Intellectual Disability into Biologically Coherent Modules. Am J Hum Genet 98, 149–164. 10.1016/j.ajhg.2015.11.024.

Krumm, N., Sudmant, P.H., Ko, A., O’Roak, B.J., Malig, M., Coe, B.P., Quinlan, A.R., Nickerson, D.A., Eichler, E.E., and Project, N.E.S. (2012). Copy number variation detection and genotyping from exome sequence data. Genome Res 22, 1525–1532. 10.1101/gr.138115.112.

Kumar, R., Corbett, M.A., van Bon, B.W., Woenig, J.A., Weir, L., Douglas, E., Friend, K.L., Gardner, A., Shaw, M., Jolly, L.A., et al. (2015). THOC2 Mutations Implicate mRNA-Export Pathway in X-Linked Intellectual Disability. Am J Hum Genet 97, 302–310. 10.1016/j.ajhg.2015.05.021.

Köhler, A., and Hurt, E. (2007). Exporting RNA from the nucleus to the cytoplasm. Nat Rev Mol Cell Biol 8, 761–773. 10.1038/nrm2255.

Lancaster, M.A., Renner, M., Martin, C.A., Wenzel, D., Bicknell, L.S., Hurles, M.E., Homfray, T., Penninger, J.M., Jackson, A.P., and Knoblich, J.A. (2013). Cerebral organoids model human brain development and microcephaly. Nature 501, 373–379. 10.1038/nature12517.

Lander, E.S., Linton, L.M., Birren, B., Nusbaum, C., Zody, M.C., Baldwin, J., Devon, K., Dewar, K., Doyle, M., FitzHugh, W., et al. (2001). Initial sequencing and analysis of the human genome. Nature 409, 860–921. 10.1038/35057062.

Lemire, G., Innes, A.M., and Boycott, K.M. (2020). THOC6 intellectual disability syndrome. GeneReviews®[Internet].

Li, B., and Dewey, C.N. (2011). RSEM: accurate transcript quantification from RNA-Seq data with or without a reference genome. BMC Bioinformatics 12, 323. 10.1186/1471-2105-12-323.

Li, H., and Durbin, R. (2009). Fast and accurate short read alignment with Burrows–Wheeler transform. bioinformatics 25, 1754–1760.

Li, Y., Muffat, J., Omer, A., Bosch, I., Lancaster, M.A., Sur, M., Gehrke, L., Knoblich, J.A., and Jaenisch, R. (2017). Induction of Expansion and Folding in Human Cerebral Organoids. Cell Stem Cell 20, 385–396.e383. 10.1016/j.stem.2016.11.017.

Licatalosi, D.D., Mele, A., Fak, J.J., Ule, J., Kayikci, M., Chi, S.W., Clark, T.A., Schweitzer, A.C., Blume, J.E., Wang, X., et al. (2008). HITS-CLIP yields genome-wide insights into brain alternative RNA processing. Nature 456, 464–469. 10.1038/nature07488.

Llorian, M., Schwartz, S., Clark, T.A., Hollander, D., Tan, L.Y., Spellman, R., Gordon, A., Schweitzer, A.C., de la Grange, P., Ast, G., and Smith, C.W. (2010). Position-dependent alternative splicing activity revealed by global profiling of alternative splicing events regulated by PTB. Nat Struct Mol Biol 17, 1114–1123. 10.1038/nsmb.1881.

Love, M.I., Huber, W., and Anders, S. (2014). Moderated estimation of fold change and dispersion for RNA-seq data with DESeq2. Genome Biol 15, 550. 10.1186/s13059-014-0550-8.

Luna, R., Rondón, A.G., and Aguilera, A. (2012). New clues to understand the role of THO and other functionally related factors in mRNP biogenesis. Biochim Biophys Acta 1819, 514–520. 10.1016/j.bbagrm.2011.11.012.

Luna, R., Rondón, A.G., Pérez-Calero, C., Salas-Armenteros, I., and Aguilera, A. (2019). The THO Complex as a Paradigm for the Prevention of Cotranscriptional R-Loops. Cold Spring Harb Symp Quant Biol 84, 105–114. 10.1101/sqb.2019.84.039594.

Luo, M.L., Zhou, Z., Magni, K., Christoforides, C., Rappsilber, J., Mann, M., and Reed, R. (2001). Pre-mRNA splicing and mRNA export linked by direct interactions between UAP56 and Aly. Nature 413, 644–647. 10.1038/35098106.

Maeder, C.I., Kim, J.I., Liang, X., Kaganovsky, K., Shen, A., Li, Q., Li, Z., Wang, S., Xu, X.Z.S., Li, J.B., et al. (2018). The THO Complex Coordinates Transcripts for Synapse Development and Dopamine Neuron Survival. Cell 174, 1436–1449.e1420. 10.1016/j.cell.2018.07.046.

Mancini, A., Niemann-Seyde, S.C., Pankow, R., El Bounkari, O., Klebba-Färber, S., Koch, A., Jaworska, E., Spooncer, E., Gruber, A.D., Whetton, A.D., and Tamura, T. (2010). THOC5/FMIP, an mRNA export TREX complex protein, is essential for hematopoietic primitive cell survival in vivo. BMC Biol 8, 1. 10.1186/1741-7007-8-1.

Martin, M. (2011). Cutadapt removes adapter sequences from high-throughput sequencing reads. EMBnet. journal 17, 10–12.

Masuda, S., Das, R., Cheng, H., Hurt, E., Dorman, N., and Reed, R. (2005). Recruitment of the human TREX complex to mRNA during splicing. Genes Dev 19, 1512–1517. 10.1101/gad.1302205.

Mattioli, F., Isidor, B., Abdul-Rahman, O., Gunter, A., Huang, L., Kumar, R., Beaulieu, C., Gecz, J., Innes, M., Mandel, J.L., and Piton, A. (2018). Clinical and functional characterization of recurrent missense variants implicated in THOC6-related intellectual disability. Hum Mol Genet. 10.1093/hmg/ddy391.

Mattioli, F., Isidor, B., Abdul-Rahman, O., Gunter, A., Huang, L., Kumar, R., Beaulieu, C., Gecz, J., Innes, M., Mandel, J.L., and Piton, A. (2019). Clinical and functional characterization of recurrent missense variants implicated in THOC6-related intellectual disability. Hum Mol Genet 28, 952–960. 10.1093/hmg/ddy391.

Mauger, O., Lemoine, F., and Scheiffele, P. (2016). Targeted Intron Retention and Excision for Rapid Gene Regulation in Response to Neuronal Activity. Neuron 92, 1266–1278. 10.1016/j.neuron.2016.11.032.

Mazin, P.V., Jiang, X., Fu, N., Han, D., Guo, M., Gelfand, M.S., and Khaitovich, P. (2018). Conservation, evolution, and regulation of splicing during prefrontal cortex development in humans, chimpanzees, and macaques. RNA 24, 585–596. 10.1261/rna.064931.117.

Mazin, P.V., Khaitovich, P., Cardoso-Moreira, M., and Kaessmann, H. (2021). Alternative splicing during mammalian organ development. Nat Genet 53, 925–934. 10.1038/s41588-021-00851-w.

Meinel, D.M., Burkert-Kautzsch, C., Kieser, A., O’Duibhir, E., Siebert, M., Mayer, A., Cramer, P., Söding, J., Holstege, F.C., and Sträßer, K. (2013). Recruitment of TREX to the transcription machinery by its direct binding to the phospho-CTD of RNA polymerase II. PLoS Genet 9, e1003914. 10.1371/journal.pgen.1003914.

Merz, C., Urlaub, H., Will, C.L., and Lührmann, R. (2007). Protein composition of human mRNPs spliced in vitro and differential requirements for mRNP protein recruitment. RNA 13, 116–128. 10.1261/rna.336807.

Meyers, E.A., and Kessler, J.A. (2017). TGF-β Family Signaling in Neural and Neuronal Differentiation, Development, and Function. Cold Spring Harb Perspect Biol 9. 10.1101/cshperspect.a022244.

Mondal, T., Subhash, S., Vaid, R., Enroth, S., Uday, S., Reinius, B., Mitra, S., Mohammed, A., James, A.R., Hoberg, E., et al. (2015). MEG3 long noncoding RNA regulates the TGF-β pathway genes through formation of RNA-DNA triplex structures. Nat Commun 6, 7743. 10.1038/ncomms8743.

Monteuuis, G., Wong, J.J.L., Bailey, C.G., Schmitz, U., and Rasko, J.E.J. (2019). The changing paradigm of intron retention: regulation, ramifications and recipes. Nucleic Acids Res 47, 11497–11513. 10.1093/nar/gkz1068.

Moon, S.L., Anderson, J.R., Kumagai, Y., Wilusz, C.J., Akira, S., Khromykh, A.A., and Wilusz, J. (2012). A noncoding RNA produced by arthropod-borne flaviviruses inhibits the cellular exoribonuclease XRN1 and alters host mRNA stability. RNA 18, 2029–2040. 10.1261/rna.034330.112.

Mor, A., Suliman, S., Ben-Yishay, R., Yunger, S., Brody, Y., and Shav-Tal, Y. (2010). Dynamics of single mRNP nucleocytoplasmic transport and export through the nuclear pore in living cells. Nat Cell Biol 12, 543–552. 10.1038/ncb2056.

Müller, H., Jimenez-Heredia, R., Krolo, A., Hirschmugl, T., Dmytrus, J., Boztug, K., and Bock, C. (2017). VCF.Filter: interactive prioritization of disease-linked genetic variants from sequencing data. Nucleic Acids Res 45, W567–W572. 10.1093/nar/gkx425.

Nagasaki, H., Arita, M., Nishizawa, T., Suwa, M., and Gotoh, O. (2005). Species-specific variation of alternative splicing and transcriptional initiation in six eukaryotes. Gene 364, 53–62. 10.1016/j.gene.2005.07.027.

Park, J.W., Tokheim, C., Shen, S., and Xing, Y. (2013). Identifying differential alternative splicing events from RNA sequencing data using RNASeq-MATS. Methods Mol Biol 1038, 171–179. 10.1007/978-1-62703-514-9_10.

Peña, A., Gewartowski, K., Mroczek, S., Cuéllar, J., Szykowska, A., Prokop, A., Czarnocki-Cieciura, M., Piwowarski, J., Tous, C., Aguilera, A., et al. (2012). Architecture and nucleic acids recognition mechanism of the THO complex, an mRNP assembly factor. EMBO J 31, 1605–1616. 10.1038/emboj.2012.10.

Pérez-Calero, C., Bayona-Feliu, A., Xue, X., Barroso, S.I., Muñoz, S., González-Basallote, V.M., Sung, P., and Aguilera, A. (2020). UAP56/DDX39B is a major cotranscriptional RNA-DNA helicase that unwinds harmful R loops genome-wide. Genes Dev 34, 898–912. 10.1101/gad.336024.119.

Pühringer, T., Hohmann, U., Fin, L., Pacheco-Fiallos, B., Schellhaas, U., Brennecke, J., and Plaschka, C. (2020). Structure of the human core transcription-export complex reveals a hub for multivalent interactions. Elife 9. 10.7554/eLife.61503.

Qian, X., Nguyen, H.N., Song, M.M., Hadiono, C., Ogden, S.C., Hammack, C., Yao, B., Hamersky, G.R., Jacob, F., Zhong, C., et al. (2016). Brain-Region-Specific Organoids Using Mini-bioreactors for Modeling ZIKV Exposure. Cell 165, 1238–1254. 10.1016/j.cell.2016.04.032.

Qu, Q., Sun, G., Murai, K., Ye, P., Li, W., Asuelime, G., Cheung, Y.T., and Shi, Y. (2013). Wnt7a regulates multiple steps of neurogenesis. Mol Cell Biol 33, 2551–2559. 10.1128/MCB.00325-13.

Retterer, K., Juusola, J., Cho, M.T., Vitazka, P., Millan, F., Gibellini, F., Vertino-Bell, A., Smaoui, N., Neidich, J., Monaghan, K.G., et al. (2016). Clinical application of whole-exome sequencing across clinical indications. Genet Med 18, 696–704. 10.1038/gim.2015.148.

Rondón, A.G., Jimeno, S., and Aguilera, A. (2010). The interface between transcription and mRNP export: from THO to THSC/TREX-2. Biochim Biophys Acta 1799, 533–538. 10.1016/j.bbagrm.2010.06.002.

Roy, B., Friesen, W.J., Tomizawa, Y., Leszyk, J.D., Zhuo, J., Johnson, B., Dakka, J., Trotta, C.R., Xue, X., Mutyam, V., et al. (2016). Ataluren stimulates ribosomal selection of near-cognate tRNAs to promote nonsense suppression. Proc Natl Acad Sci U S A 113, 12508–12513. 10.1073/pnas.1605336113.

Ruaud, L., Roux, N., Boutaud, L., Bessières, B., Ageorges, F., Achaiaa, A., Bole, C., Nitschke, P., Masson, C., Vekemans, M., et al. (2022). Biallelic THOC6 pathogenic variants: Prenatal phenotype and review of the literature. Birth Defects Res 114, 499–504. 10.1002/bdr2.2011.

Schalock, R.L., Luckasson, R., and Tassé, M.J. (2021). An Overview of Intellectual Disability: Definition, Diagnosis, Classification, and Systems of Supports (12th ed.). Am J Intellect Dev Disabil 126, 439-442. 10.1352/1944-7558-126.6.439.

Schindelin, J., Arganda-Carreras, I., Frise, E., Kaynig, V., Longair, M., Pietzsch, T., Preibisch, S., Rueden, C., Saalfeld, S., Schmid, B., et al. (2012). Fiji: an open-source platform for biological-image analysis. Nat Methods 9, 676–682. 10.1038/nmeth.2019.

Shen, J., Zhang, L., and Zhao, R. (2007). Biochemical characterization of the ATPase and helicase activity of UAP56, an essential pre-mRNA splicing and mRNA export factor. J Biol Chem 282, 22544–22550. 10.1074/jbc.M702304200.

Shen, S., Park, J.W., Lu, Z.X., Lin, L., Henry, M.D., Wu, Y.N., Zhou, Q., and Xing, Y. (2014). rMATS: robust and flexible detection of differential alternative splicing from replicate RNA-Seq data. Proc Natl Acad Sci U S A 111, E5593–5601. 10.1073/pnas.1419161111.

Shi, M., Zhang, H., Wu, X., He, Z., Wang, L., Yin, S., Tian, B., Li, G., and Cheng, H. (2017). ALYREF mainly binds to the 5’ and the 3’ regions of the mRNA in vivo. Nucleic Acids Res 45, 9640–9653. 10.1093/nar/gkx597.

Sobreira, N., Schiettecatte, F., Valle, D., and Hamosh, A. (2015). GeneMatcher: a matching tool for connecting investigators with an interest in the same gene. Hum Mutat 36, 928–930. 10.1002/humu.22844.

Srivastava, A., Srivastava, K.R., Hebbar, M., Galada, C., Kadavigrere, R., Su, F., Cao, X., Chinnaiyan, A.M., Girisha, K.M., Shukla, A., and Bielas, S.L. (2018). Genetic diversity of NDUFV1-dependent mitochondrial complex I deficiency. Eur J Hum Genet 26, 1582–1587. 10.1038/s41431-018-0209-0.

Stirling, D.R., Swain-Bowden, M.J., Lucas, A.M., Carpenter, A.E., Cimini, B.A., and Goodman, A. (2021). CellProfiler 4: improvements in speed, utility and usability. BMC Bioinformatics 22, 433. 10.1186/s12859-021-04344-9.

Strässer, K., and Hurt, E. (2001). Splicing factor Sub2p is required for nuclear mRNA export through its interaction with Yra1p. Nature 413, 648–652. 10.1038/35098113.

Strässer, K., Masuda, S., Mason, P., Pfannstiel, J., Oppizzi, M., Rodriguez-Navarro, S., Rondón, A.G., Aguilera, A., Struhl, K., Reed, R., and Hurt, E. (2002). TREX is a conserved complex coupling transcription with messenger RNA export. Nature 417, 304–308. 10.1038/nature746.

Subramanian, A., Tamayo, P., Mootha, V.K., Mukherjee, S., Ebert, B.L., Gillette, M.A., Paulovich, A., Pomeroy, S.L., Golub, T.R., Lander, E.S., and Mesirov, J.P. (2005). Gene set enrichment analysis: a knowledge-based approach for interpreting genome-wide expression profiles. Proc Natl Acad Sci U S A 102, 15545–15550. 10.1073/pnas.0506580102.

Taniguchi, I., and Ohno, M. (2008). ATP-dependent recruitment of export factor Aly/REF onto intronless mRNAs by RNA helicase UAP56. Mol Cell Biol 28, 601–608. 10.1128/MCB.01341-07.

Team, R.C. (2018). R: A language and environment for statistical computing. https://www.R-project.org/.

Tran, D.D., Saran, S., Williamson, A.J., Pierce, A., Dittrich-Breiholz, O., Wiehlmann, L., Koch, A., Whetton, A.D., and Tamura, T. (2014). THOC5 controls 3’end-processing of immediate early genes via interaction with polyadenylation specific factor 100 (CPSF100). Nucleic Acids Res 42, 12249–12260. 10.1093/nar/gku911.

Truett, G.E., Heeger, P., Mynatt, R.L., Truett, A.A., Walker, J.A., and Warman, M.L. (2000). Preparation of PCR-quality mouse genomic DNA with hot sodium hydroxide and tris (HotSHOT). Biotechniques 29, 52, 54. 10.2144/00291bm09.

Tsui, D., Vessey, J.P., Tomita, H., Kaplan, D.R., and Miller, F.D. (2013). FoxP2 regulates neurogenesis during embryonic cortical development. J Neurosci 33, 244–258. 10.1523/JNEUROSCI.1665-12.2013.

Van der Auwera, G.A., Carneiro, M.O., Hartl, C., Poplin, R., Del Angel, G., Levy-Moonshine, A., Jordan, T., Shakir, K., Roazen, D., Thibault, J., et al. (2013). From FastQ data to high confidence variant calls: the Genome Analysis Toolkit best practices pipeline. Curr Protoc Bioinformatics 11, 11.10.11-11.10.33.10.1002/0471250953.bi1110s43.

Vasudevan, P., and Suri, M. (2017). A clinical approach to developmental delay and intellectual disability. Clin Med (Lond) 17, 558–561. 10.7861/clinmedicine.17-6-558.

Veiga, D.F.T. (2022). maser: Mapping Alternative Splicing Events to pRoteins.

Venter, J.C., Adams, M.D., Myers, E.W., Li, P.W., Mural, R.J., Sutton, G.G., Smith, H.O., Yandell, M., Evans, C.A., Holt, R.A., et al. (2001). The sequence of the human genome. Science 291, 1304–1351. 10.1126/science.1058040.

Viphakone, N., Sudbery, I., Griffith, L., Heath, C.G., Sims, D., and Wilson, S.A. (2019). Co-transcriptional Loading of RNA Export Factors Shapes the Human Transcriptome. Mol Cell 75, 310–323.e318. 10.1016/j.molcel.2019.04.034.

Vogel, T., Ahrens, S., Büttner, N., and Krieglstein, K. (2010). Transforming growth factor beta promotes neuronal cell fate of mouse cortical and hippocampal progenitors in vitro and in vivo: identification of Nedd9 as an essential signaling component. Cereb Cortex 20, 661–671. 10.1093/cercor/bhp134.

Wang, E.T., Sandberg, R., Luo, S., Khrebtukova, I., Zhang, L., Mayr, C., Kingsmore, S.F., Schroth, G.P., and Burge, C.B. (2008). Alternative isoform regulation in human tissue transcriptomes. Nature 456, 470–476. 10.1038/nature07509.

Wang, H., Ge, G., Uchida, Y., Luu, B., and Ahn, S. (2011). Gli3 is required for maintenance and fate specification of cortical progenitors. J Neurosci 31, 6440–6448. 10.1523/JNEUROSCI.4892-10.2011.

Wang, L., Miao, Y.L., Zheng, X., Lackford, B., Zhou, B., Han, L., Yao, C., Ward, J.M., Burkholder, A., Lipchina, I., et al. (2013). The THO complex regulates pluripotency gene mRNA export and controls embryonic stem cell self-renewal and somatic cell reprogramming. Cell Stem Cell 13, 676–690. 10.1016/j.stem.2013.10.008.

Wang, Y., Liu, J., Huang, B.O., Xu, Y.M., Li, J., Huang, L.F., Lin, J., Zhang, J., Min, Q.H., Yang, W.M., and Wang, X.Z. (2015). Mechanism of alternative splicing and its regulation. Biomed Rep 3, 152–158. 10.3892/br.2014.407.

Weyn-Vanhentenryck, S.M., Feng, H., Ustianenko, D., Duffié, R., Yan, Q., Jacko, M., Martinez, J.C., Goodwin, M., Zhang, X., Hengst, U., et al. (2018). Precise temporal regulation of alternative splicing during neural development. Nat Commun 9, 2189. 10.1038/s41467-018-04559-0.

Wickramasinghe, V.O., and Laskey, R.A. (2015). Control of mammalian gene expression by selective mRNA export. Nat Rev Mol Cell Biol 16, 431–442. 10.1038/nrm4010.

Wingett, S.W., and Andrews, S. (2018). FastQ Screen: A tool for multi-genome mapping and quality control. F1000Res 7, 1338. 10.12688/f1000research.15931.2.

Xie, Y., and Ren, Y. (2019). Mechanisms of nuclear mRNA export: A structural perspective. Traffic 20, 829–840. 10.1111/tra.12691.

Yang, Y., Muzny, D.M., Xia, F., Niu, Z., Person, R., Ding, Y., Ward, P., Braxton, A., Wang, M., Buhay, C., et al. (2014). Molecular findings among patients referred for clinical whole-exome sequencing. JAMA 312, 1870–1879. 10.1001/jama.2014.14601.

Yap, K., Lim, Z.Q., Khandelia, P., Friedman, B., and Makeyev, E.V. (2012). Coordinated regulation of neuronal mRNA steady-state levels through developmentally controlled intron retention. Genes Dev 26, 1209–1223. 10.1101/gad.188037.112.

Yeo, G., and Burge, C.B. (2004). Maximum entropy modeling of short sequence motifs with applications to RNA splicing signals. J Comput Biol 11, 377–394. 10.1089/1066527041410418.

Zhang, Q., Chen, S., Qin, Z., Zheng, H., and Fan, X. (2020). The first reported case of Beaulieu-Boycott-Innes syndrome caused by two novel mutations in THOC6 gene in a Chinese infant. Medicine (Baltimore) 99, e19751. 10.1097/MD.0000000000019751.

Zhao, J., Zhang, X., Zhou, Y., Ansell, P.J., and Klibanski, A. (2006). Cyclic AMP stimulates MEG3 gene expression in cells through a cAMP-response element (CRE) in the MEG3 proximal promoter region. Int J Biochem Cell Biol 38, 1808–1820. 10.1016/j.biocel.2006.05.004.

Zhou, S., Zhong, Z., Huang, P., Xiang, B., Li, X., Dong, H., Zhang, G., Wu, Y., and Li, P. (2021). IL-6/STAT3 Induced Neuron Apoptosis in Hypoxia by Downregulating ATF6 Expression. Front Physiol 12, 729925. 10.3389/fphys.2021.729925.

Zhou, Y., Zhou, B., Pache, L., Chang, M., Khodabakhshi, A.H., Tanaseichuk, O., Benner, C., and Chanda, S.K. (2019). Metascape provides a biologist-oriented resource for the analysis of systems-level datasets. Nat Commun 10, 1523. 10.1038/s41467-019-09234-6.

Zuckerman, B., Ron, M., Mikl, M., Segal, E., and Ulitsky, I. (2020). Gene Architecture and Sequence Composition Underpin Selective Dependency of Nuclear Export of Long RNAs on NXF1 and the TREX Complex. Mol Cell 79, 251–267.e256. 10.1016/j.molcel.2020.05.013.

Zylka, M.J., Simon, J.M., and Philpot, B.D. (2015). Gene length matters in neurons. Neuron 86, 353–355. 10.1016/j.neuron.2015.03.059.

