## Supplementary material for "Mechanisms of mRNA processing defects in inherited *THOC6* intellectual disability syndrome": Figure S1

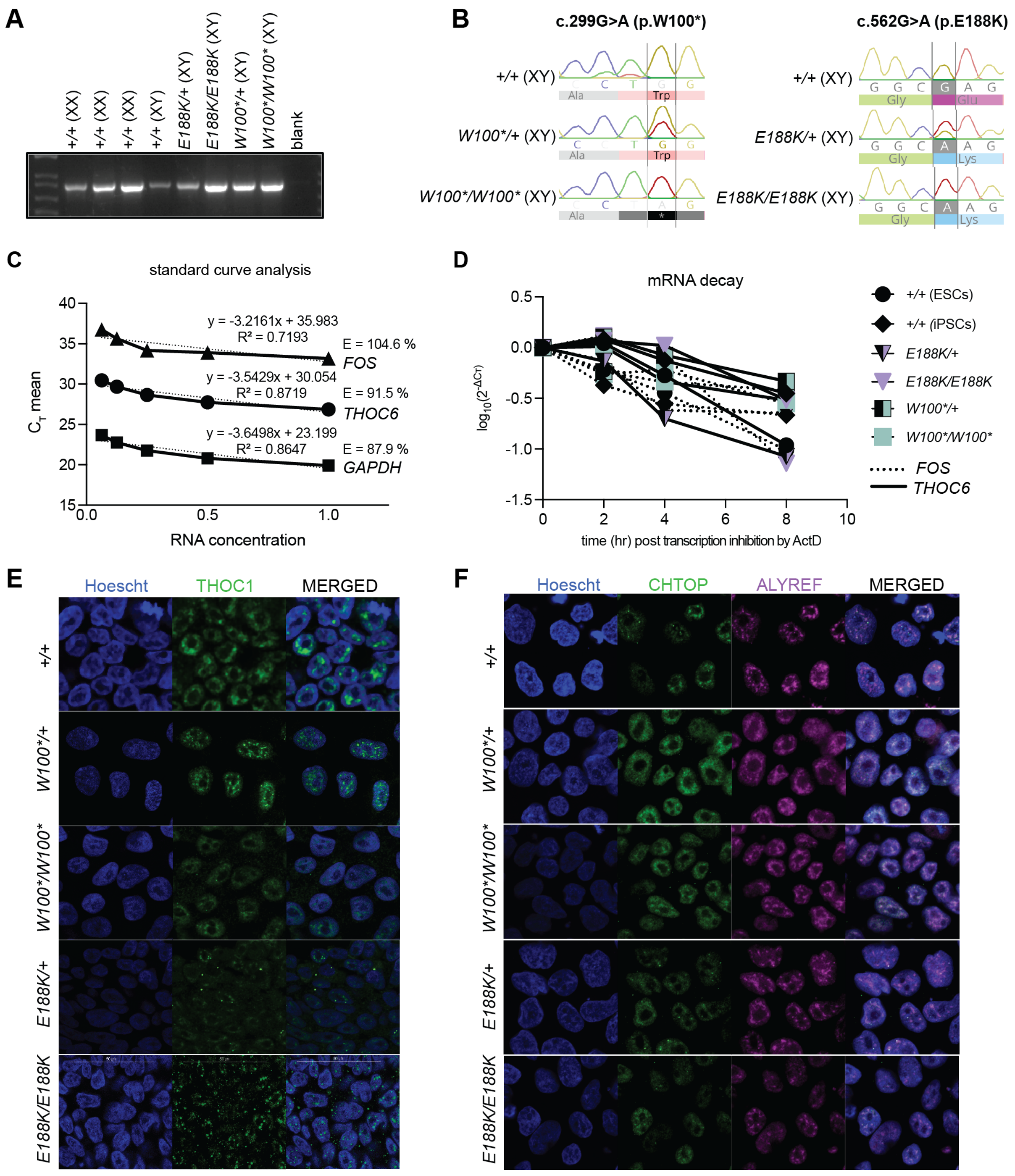
A
+/+ (XX)
+/+ (XX)
+/+ (XX)
+/+ (XY)
E188K/+ (XY)
E188K/E188K (XY)
W100*/+ (XY)
W100*/W100* (XY)
blank
B
c.299G>A (p.W100*)
c.562G>A (p.E188K)
+/+ (XY)
W100*/+ (XY)
W100*/W100* (XY)
E188K/+ (XY)
E188K/E188K (XY)
Ala
Trp
Gly
Glu
Lys
C
standard curve analysis
y = -3.2161x + 35.983
R² = 0.7193
E = 104.6 %
FOS
y = -3.5429x + 30.054
R² = 0.8719
E = 91.5 %
THOC6
y = -3.6498x + 23.199
R² = 0.8647
E = 87.9 %
GAPDH
C_T mean
RNA concentration
D
mRNA decay
log10(2^-ΔCT)
time (hr) post transcription inhibition by ActD
+/+ (ESCs)
+/+ (iPSCs)
E188K/+
E188K/E188K
W100*/+
W100*/W100*
FOS
THOC6
E
Hoescht
THOC1
MERGED
+/+
W100*/+
W100*/W100*
E188K/+
E188K/E188K
F
Hoescht
CHTOP
ALYREF
MERGED
+/+
W100*/+
W100*/W100*
E188K/+
E188K/E188K

**Figure S1. Genetic mechanism of biallelic pathogenic *THOC6* variants**. *See also Figures 1 and 2.* Confirmation of genotypes in human ESC/iPSC lines by PCR (A) and Sanger sequencing (B). (C) Standard curve analysis using 5 two-fold serial dilutions of control cDNA to confirm primer quality for *FOS, THOC6,* and *GAPDH* for mRNA stability assay*.* (D) mRNA decay curve for extended timeframe capturing *THOC6* and *FOS* RNA decay after 2.5 hrs following Actinomysin D treatment. Values were not normalized to *GAPDH* because control transcripts are also degraded after 2.5 hrs of transcription inhibition. Immunostaining to assess subcellular localization of THO/TREX complex members THOC1 (E), and CHTOP and ALYREF (F) in human ESC/iPSCs with biallelic pathogenic *THOC6* variants compared to heterozygous and wildtype unaffected controls.
