## Supplementary material for "Mechanisms of mRNA processing defects in inherited *THOC6* intellectual disability syndrome": Figure S2

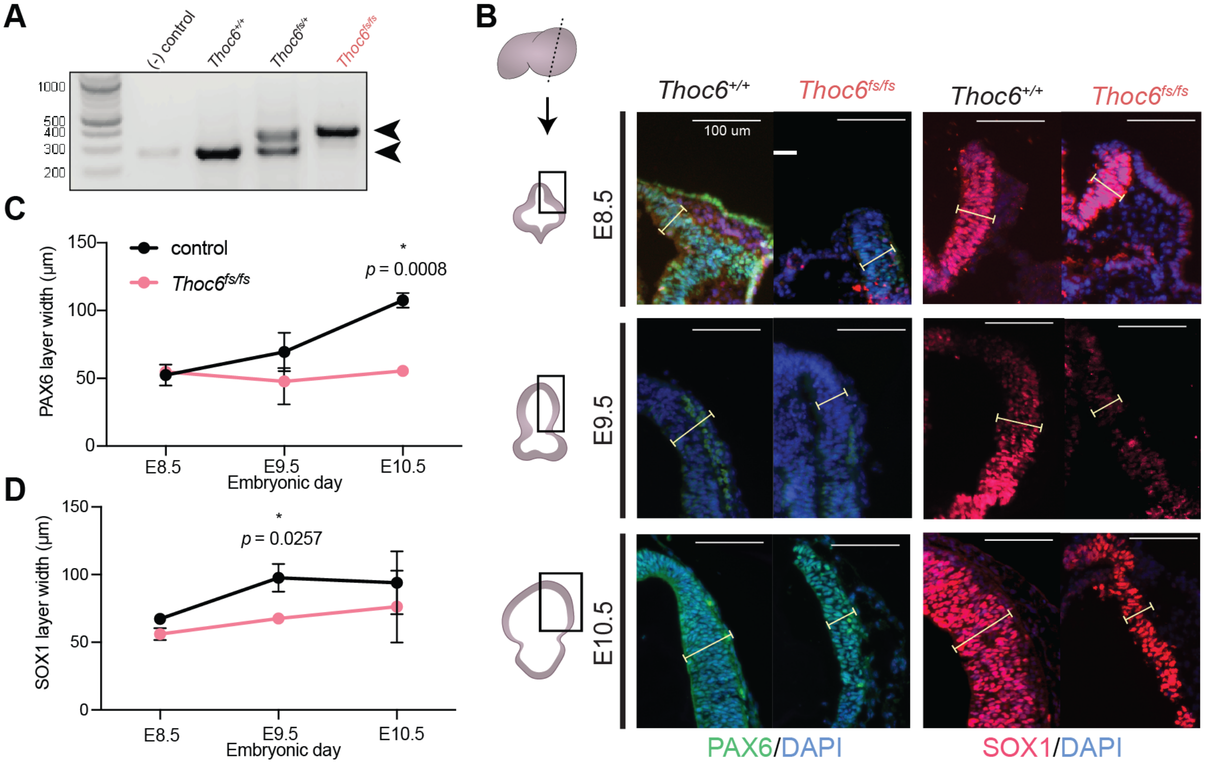

A
(-) control
Thoc6+/+
Thoc6fs/+
Thoc6fs/fs
1000
500
400
300
200
B
Thoc6+/+
Thoc6fs/fs
Thoc6+/+
Thoc6fs/fs
100 um
E8.5
E9.5
E10.5
PAX6/DAPI
SOX1/DAPI
C
control
Thoc6fs/fs
*
p = 0.0008
PAX6 layer width (μm)
150
100
50
0
E8.5
E9.5
E10.5
Embryonic day
D
*
p = 0.0257
SOX1 layer width (μm)
150
100
50
0
E8.5
E9.5
E10.5
Embryonic day

**Figure S2. Generation of *Thoc6$^{fs/fs}$* mouse model.** *See also Figure 3.* (A) Gel of allele-specific PCR confirming *Thoc6* frameshift alleles. (B) Immunostaining of the neural plate and neural tube in *Thoc6$^{fs/fs}$* mice relative to *Thoc6$^{+/+}$* control littermates at E8.5-10.5 for Pax6 (C) and Sox1 (D), with layer width quantifications (right). Two-sided paired Student's *t* test. *n.s.*, not significant. Scale bar: 100 μm.
