## Supplementary material for "Mechanisms of mRNA processing defects in inherited *THOC6* intellectual disability syndrome": Figure S3

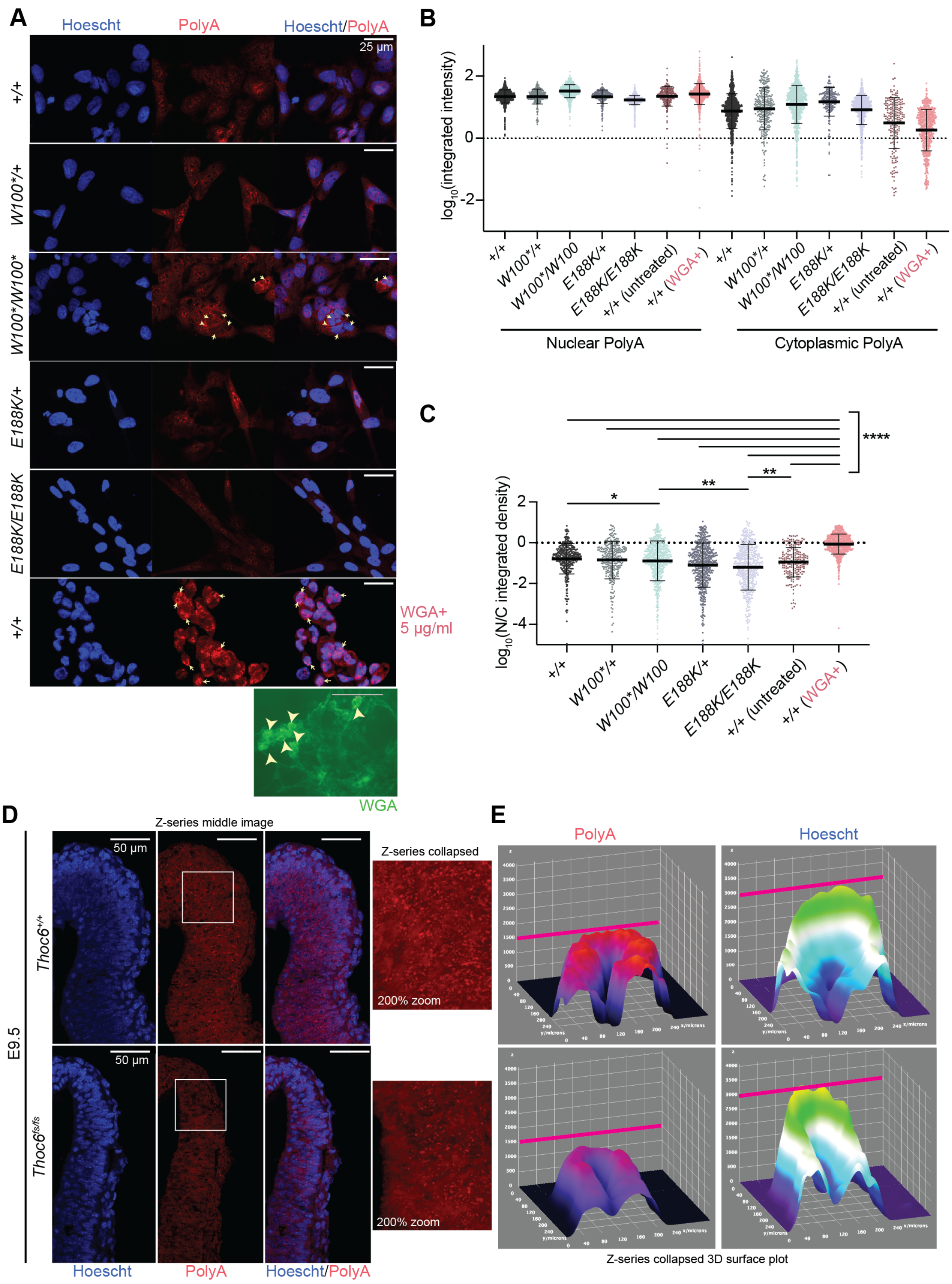
A
Hoescht
PolyA
Hoescht/PolyA
25 μm
+/+
W100*/+
W100*/W100*
E188K/+
E188K/E188K
+/+
WGA+
5 μg/ml
WGA
B
log10(integrated intensity)
2
0
-2
+/+
W100*/+
W100*/W100
E188K/+
E188K/E188K
+/+ (untreated)
+/+ (WGA+)
Nuclear PolyA
Cytoplasmic PolyA
C
****
*
**
**
log10(N/C integrated density)
2
0
-2
-4
+/+
W100*/+
W100*/W100
E188K/+
E188K/E188K
+/+ (untreated)
+/+ (WGA+)
D
Z-series middle image
Z-series collapsed
50 μm
E9.5
Thoc6+/+
Thoc6fs/fs
200% zoom
Hoescht
PolyA
Hoescht/PolyA
E
PolyA
Hoescht
Z-series collapsed 3D surface plot

**Figure S3. Characterization of bulk mRNA export across genotypes.** *Related to Figures 3-6).* (A) Oligo-dT FISH Z-collapsed confocal images (40x magnification) of NPCs differentiated from iPSCs with the following genotypes: *THOC6*$^{+/+}$, *THOC6*$^{W100*/W100*}$, *THOC6*$^{W100*/+}$, *THOC6*$^{E188K/E188K}$, *THOC6*$^{E188K/+}$. Arrows point to cells showing extreme signal differences. As a positive control for impaired nuclear export resulting in aberrant accumulation of transcripts in the nucleus, *THOC6*$^{+/+}$ were treated with WGA at 5 μg/ml which acts to block the nuclear pore complex (bottom). Scale bar: 25 μm. (B) Intensity quantifications from 200-700 cells per genotype from three replicates were performed using an automated CellProfiler (v4.2.1) pipeline (Stirling et al., 2021) that measures polyA signal in nuclear and cytoplasmic fractions. Variability in polyA+ signal intensity observed across genotypes is likely due to slight technical variation in the assay across replicate slides. (C) Ratios of nuclear to cytoplasmic polyA showing minimal differences in bulk export across genotypes relative to WGA+ positive control. (D) Oligo-dT FISH of E9.5 *Thoc6*$^{+/+}$ and *Thoc6*$^{fs/fs}$ mouse neuroepithelium; Z-series middle image (left). Scale bar: 50 μm. White box represents zoomed in area. (Right) 200% zoom of polyA signal in Z-series collapsed image by maximum intensity. (E) 3D surface plot for Z-series collapsed image of polyA intensity (left) and Hoescht intensity (right) in *Thoc6*$^{+/+}$ E9.5 neuroepithelium (top) and *Thoc6*$^{fs/fs}$ E9.5 neuroepithelium (bottom).
