## Supplementary material for "Mechanisms of mRNA processing defects in inherited *THOC6* intellectual disability syndrome": Figure S4

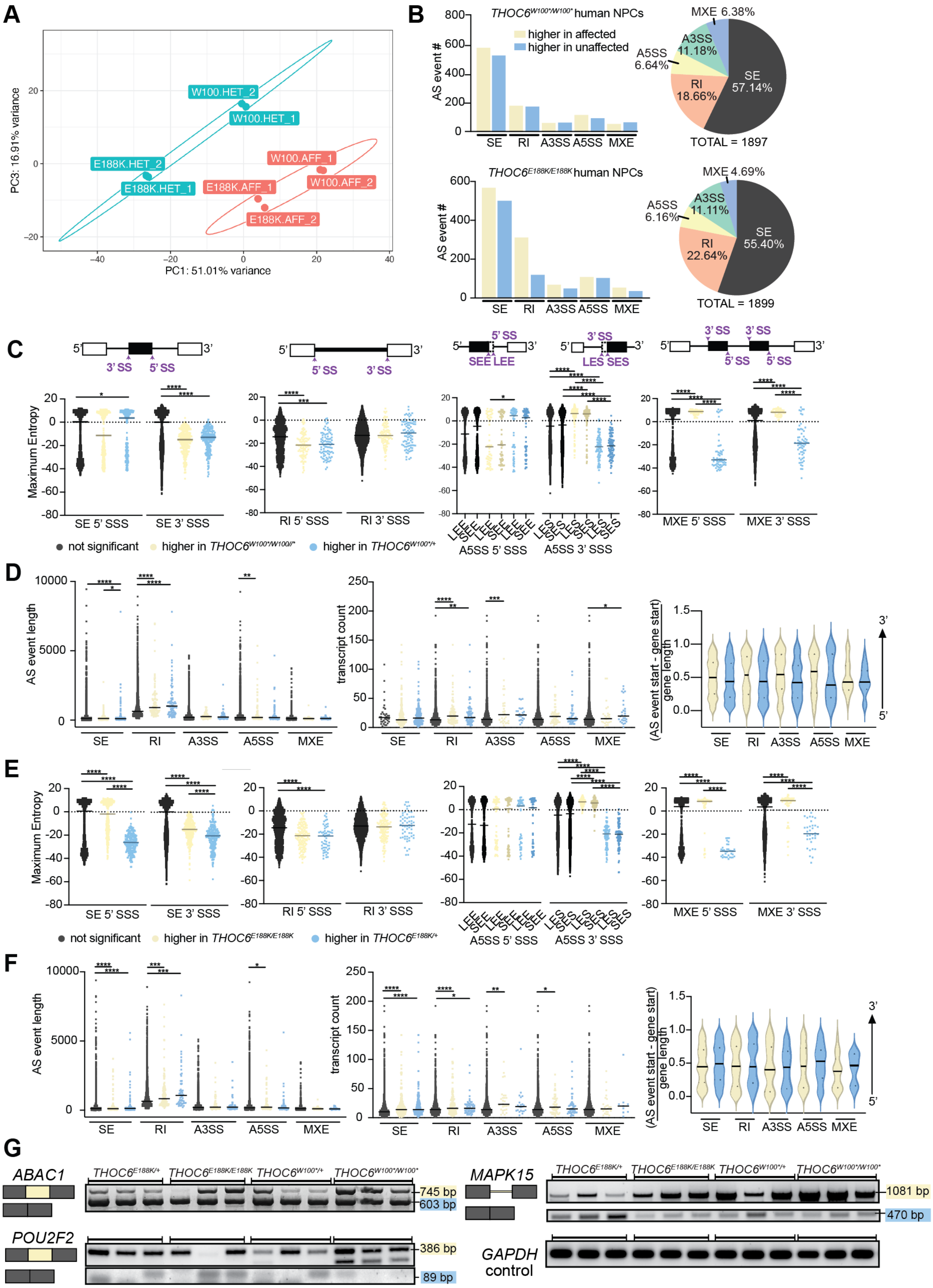

A
PC1: 51.01% variance
PC3: 16.91% variance
W100.HET_2
W100.HET_1
E188K.HET_2
E188K.HET_1
W100.AFF_1
W100.AFF_2
E188K.AFF_1
E188K.AFF_2
B
THOC6W100*/W100* human NPCs
higher in affected
higher in unaffected
AS event #
SE RI A3SS A5SS MXE
MXE 6.38%
A3SS 11.18%
A5SS 6.64%
RI 18.66%
SE 57.14%
TOTAL = 1897
THOC6E188K/E188K human NPCs
MXE 4.69%
A5SS 6.16%
A3SS 11.11%
RI 22.64%
SE 55.40%
TOTAL = 1899
C
3' SS
5' SS
SEE LEE
LES SES
Maximum Entropy
SE 5' SSS
SE 3' SSS
RI 5' SSS
RI 3' SSS
A5SS 5' SSS
A5SS 3' SSS
MXE 5' SSS
MXE 3' SSS
not significant
higher in THOC6W100*/W100/*
higher in THOC6W100*/+
D
AS event length
transcript count
(AS event start - gene start) / gene length
SE RI A3SS A5SS MXE
E
not significant
higher in THOC6E188K/E188K
higher in THOC6E188K/+
F
G
ABAC1
THOC6E188K/+
THOC6E188K/E188K
THOC6W100*/+
THOC6W100*/W100*
745 bp
603 bp
POU2F2
386 bp
89 bp
MAPK15
1081 bp
470 bp
GAPDH control

**Figure S4. Characterization of alternative splicing events in *THOC6* affected hNPCs.** *See also Figure 4.* (A) PCA of hNPCs RNAseq samples showing separation by condition (unaffected (E188K_HET, W100_HET) and affected (E188K_AFF, W100_AFF)). (B) rMATS summary analysis on *THOC6*$^{W100*/W100*}$ versus *THOC6*$^{W100*/+}$ hNPCs (top) and *THOC6*$^{E188K/E188K}$ versus *THOC6*$^{E188K/+}$ hNPCs (bottom). Differences in splicing patterns were comparable in both *THOC6*$^{E188K/E188K}$ versus *THOC6*$^{E188K/+}$ and *THOC6*$^{E188K/E188K}$ versus *THOC6*$^{W100*/+}$ comparisons, as well as compared to *THOC6*$^{W100*/W100*}$ versus *THOC6*$^{W100*/+}$ (see Table S5 and Figure 4). Blue, excluded in affected; yellow, included in affected. 5' and 3' splice site strengths (SSS) (C), and event length, transcript count, and event position (D) per AS event in *THOC6*$^{W100*/W100*}$ hNPCs. Long exon start, LES; short exon start, SES; long exon end, LEE; short exon end, SEE. 5' and 3' splice site strengths (E), and event length, transcript count, and event position (F) per AS event in *THOC6*$^{E188K/E188K}$ hNPCs. (G) RT-PCR gels across three biological hNPC replicates per genotype for *ABAC1*, *POU2F2*, and *MAPK15* AS events and *GAPDH* loading control. Some variation across replicates was noted.
