## Supplementary material for "Mechanisms of mRNA processing defects in inherited *THOC6* intellectual disability syndrome": Figure S5

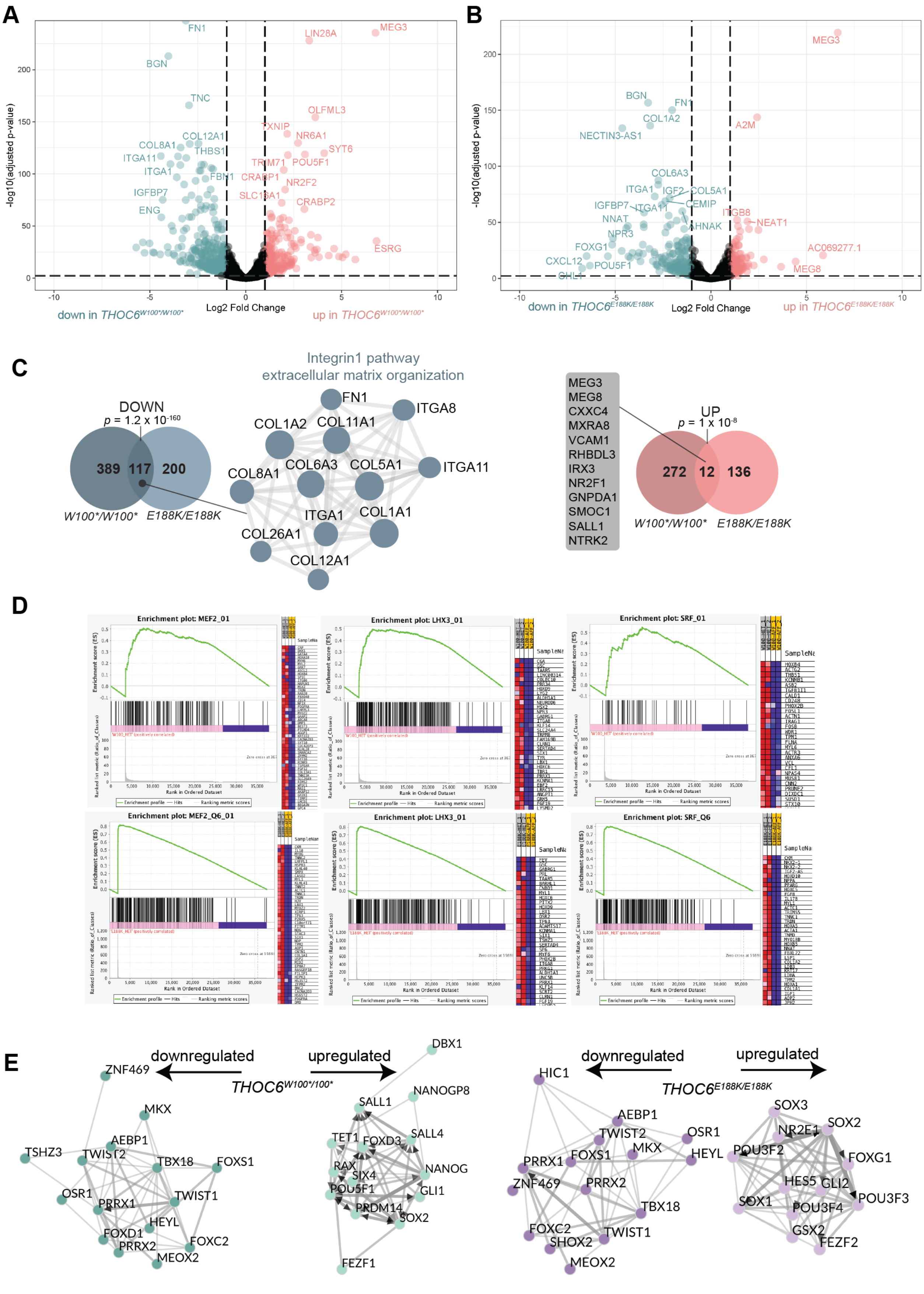
A
B
C
D
E
down in THOC6W100*/W100*
up in THOC6W100*/W100*
Log2 Fold Change
-log10(adjusted p-value)
down in THOC6E188K/E188K
up in THOC6E188K/E188K
Integrin1 pathway
extracellular matrix organization
DOWN
p = 1.2 x 10-160
389
117
200
W100*/W100*
E188K/E188K
UP
p = 1 x 10-8
272
12
136
MEG3
MEG8
CXXC4
MXRA8
VCAM1
RHBDL3
IRX3
NR2F1
GNPDA1
SMOC1
SALL1
NTRK2
Enrichment plot: MEF2_01
Enrichment plot: LHX3_01
Enrichment plot: SRF_01
Enrichment plot: MEF2_Q6_01
Enrichment plot: LHX3_01
Enrichment plot: SRF_Q6
downregulated
upregulated
THOC6W100*/100*
THOC6E188K/E188K

**Figure S5. Differential expression analysis in affected hNPCs.** *See also Figure 5.* Volcano plot of differential expression in *THOC6*$^{W100*/W100*}$ (A) and *THOC6*$^{E188K/E188K}$ (B) NPCs relative to *THOC6*$^{W100*/+}$ controls. Data represent analysis of two biological replicates per genotype. The later comparison was chosen following observation of upregulated skeletal muscle genes in *THOC6*$^{E188K/+}$ hNPC replicates, indicating issues during the differentiation of this line. (C) Gene overlap of *THOC6*$^{W100*/W100*}$ and *THOC6*$^{E188K/E188K}$ downregulated (left, blue) and upregulated (right, red) genes. Metascape protein-protein network enrichment analysis identifies integrin1 pathway and extracellular matrix modules enriched among genes downregulated in both genotypes. (D) GSEA enrichment plots for top transcription factors using the transcription factor motif gene dataset (c4.tftv7.5.1.symbols.gmt) in *THOC6*$^{W100*/W100*}$ (top) *THOC6*$^{E188K/E188K}$ (bottom). (E) ChEA3 transcription factor motif enrichment of DEGs in *THOC6*$^{W100*/W100*}$ (left, green) and *THOC6*$^{E188K/E188K}$ (right, purple) NPCs.
