## Supplementary material for "Mechanisms of mRNA processing defects in inherited *THOC6* intellectual disability syndrome": Figure S6

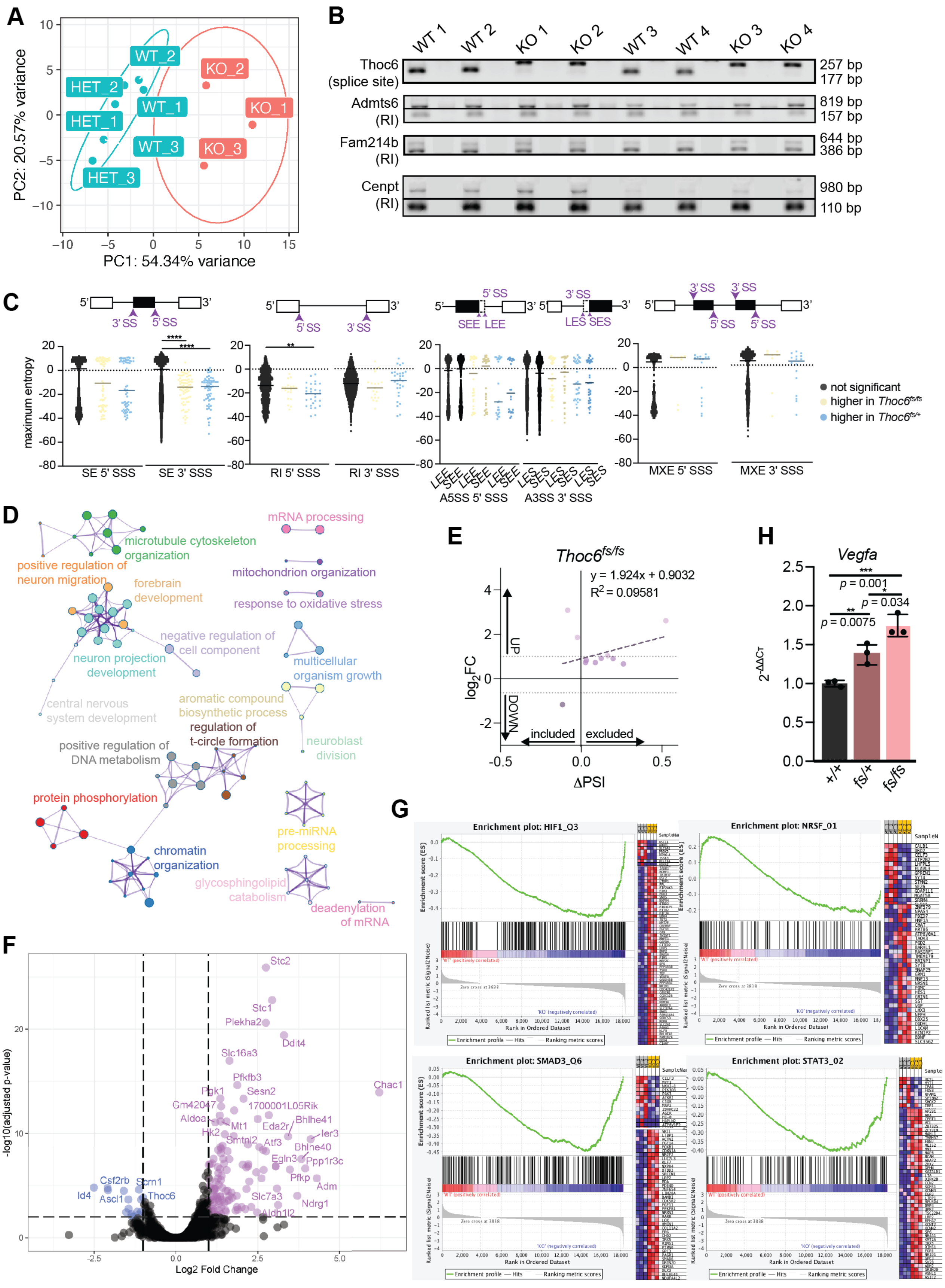

A
PC1: 54.34% variance
PC2: 20.57% variance
WT_1
WT_2
WT_3
HET_1
HET_2
HET_3
KO_1
KO_2
KO_3
B
WT 1
WT 2
KO 1
KO 2
WT 3
WT 4
KO 3
KO 4
Thoc6 (splice site)
Admts6 (RI)
Fam214b (RI)
Cenpt (RI)
257 bp
177 bp
819 bp
157 bp
644 bp
386 bp
980 bp
110 bp
C
maximum entropy
SE 5' SSS
SE 3' SSS
RI 5' SSS
RI 3' SSS
A5SS 5' SSS
A3SS 3' SSS
MXE 5' SSS
MXE 3' SSS
not significant
higher in Thoc6fs/fs
higher in Thoc6fs/+
D
mRNA processing
microtubule cytoskeleton organization
positive regulation of neuron migration
forebrain development
mitochondrion organization
response to oxidative stress
negative regulation of cell component
neuron projection development
multicellular organism growth
central nervous system development
aromatic compound biosynthetic process
regulation of t-circle formation
neuroblast division
positive regulation of DNA metabolism
protein phosphorylation
chromatin organization
pre-miRNA processing
glycosphingolipid catabolism
deadenylation of mRNA
E
Thoc6fs/fs
y = 1.924x + 0.9032
R2 = 0.09581
log2FC
ΔPSI
UP
DOWN
included
excluded
F
-log10(adjusted p-value)
Log2 Fold Change
Stc2
Stc1
Plekha2
Ddit4
Slc16a3
Chac1
Pfkfb3
Pgk1
Sesn2
Gm42047
1700001L05Rik
Aldoa
Bhlhe41
Mt1
Eda2r
Hk2
Ier3
Smtnl2
Atf3
Bhlhe40
Egln3
Ppp1r3c
Pfkp
Adm
Csf2rb
Scrn1
Id4
Ascl1
Thoc6
Slc7a3
Ndrg1
Aldh1l2
G
Enrichment plot: HIF1_Q3
Enrichment plot: NRSF_01
Enrichment plot: SMAD3_Q6
Enrichment plot: STAT3_02
Enrichment score (ES)
Rank in Ordered Dataset
Ranked list metric (Signal2Noise)
Enrichment profile
Hits
Ranking metric scores
H
Vegfa
2-ΔΔCT
p = 0.001
p = 0.034
p = 0.0075
+/+
fs/+
fs/fs

**Figure S6. Characterization of mRNA processing defects in *Thoc6*$^{fs/fs}$ mouse E9.5 forebrain.** *See also Figure 6.* (A) PCA of mouse E9.5 forebrain RNAseq samples showing separation by genotype (wildtype (WT), heterozygous (HET), homozygous for frameshift (KO). (B) RT-PCR gels across four biological replicates per genotype in two litters showing *Thoc6*, *Admts6*, *Fam214b*, and *Cenpt* AS events. (C) 5' and 3' splice site strengths (SSS) across AS events in *Thoc6*$^{fs/fs}$ mouse E9.5 forebrain. Blue, excluded in affected; yellow, included in affected. Long exon start, LES; short exon start, SES; long exon end, LEE; short exon end, SEE. (D) Metascape visualization of enriched biological categories among AS genes in *Thoc6*$^{fs/fs}$ with FDR < 0.05, ΔPSI < -0.1 or > 0.1, and average read coverage > 5. (E) Linear regression analysis of $log_2$foldchange and Δ percent transcripts spliced in (PSI) for significant retained intron events in *Thoc6*$^{fs/fs}$ cells. (F) Volcano plot of differential expression analysis in *Thoc6*$^{fs/fs}$ relative to *Thoc6*$^{+/+}$ control. Data represent three biological replicates per genotype. (G) GSEA enrichment plots for top transcription factors for in E9.5 *Thoc6*$^{fs/fs}$ forebrain. (H) qPCR relative abundance for mouse E9.5 forebrain (relative to wildtype) for *Vegfa.* Three technical replicates of two biological replicates per genotype. Significance, two-tailed unpaired *t* test. Data shown as mean ±SEM.
