## Supplementary material for "Mechanisms of mRNA processing defects in inherited *THOC6* intellectual disability syndrome": Figure S7

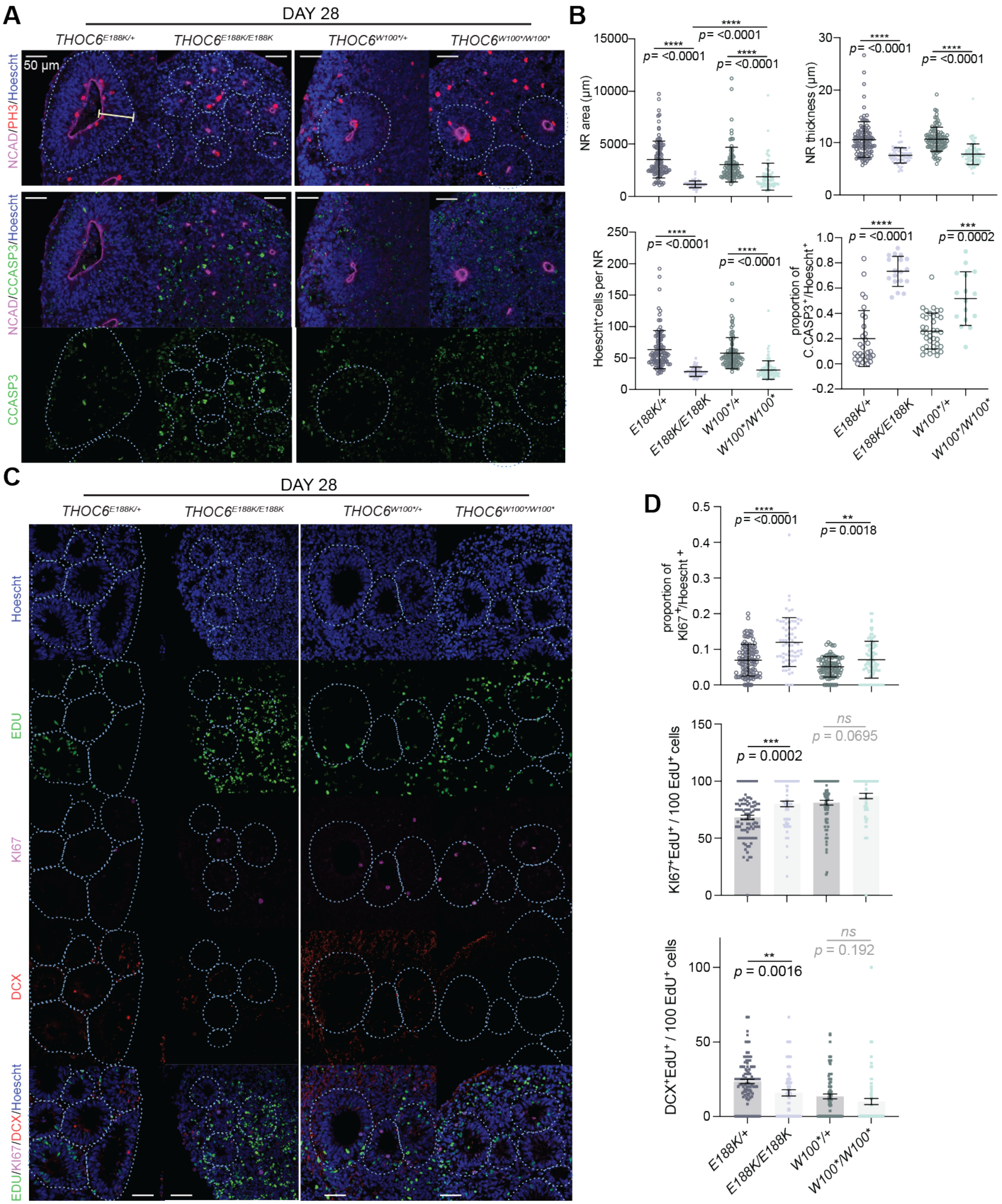
A
DAY 28
THOC6E188K/+
THOC6E188K/E188K
THOC6W100*/+
THOC6W100*/W100*
50 µm
NCAD/PH3/Hoescht
NCAD/CCASP3/Hoescht
CCASP3
B
NR area (µm)
NR thickness (µm)
Hoescht+cells per NR
proportion of C. CASP3+/Hoescht+
p= <0.0001
p= 0.0002
E188K/+
E188K/E188K
W100*/+
W100*/W100*
C
DAY 28
Hoescht
EDU
KI67
DCX
EDU/KI67/DCX/Hoescht
D
proportion of KI67+/Hoescht+
p= <0.0001
p= 0.0018
KI67+EdU+ / 100 EdU+ cells
p = 0.0002
ns
p = 0.0695
DCX+EdU+ / 100 EdU+ cells
p = 0.0016
ns
p = 0.192
E188K/+
E188K/E188K
W100*/+
W100*/W100*

**Figure S7. Modeling of *THOC6* variant pathogenesis in human cerebral organoids.** *See also Figure 7.* (A) Immunostaining of PH3, N-Cadherin, C.CASP3, and Hoescht in day 28 human cerebral organoids differentiated from $THOC6^{E188K/+}$, $THOC6^{E188K/E188K}$, $THOC6^{W100*/+}$, and $THOC6^{W100*/W100*}$ iPSCs highlighting differences in neural rosette morphology. 40x magnification; Scale bar: 50 µm. (B) Quantifications by genotype of area, thickness, Hoescht+ cells, and C.CASP3+ cells per NR. NR (organoid) number analyzed across one differentiation replicate per genotype: $THOC6^{E188K/+}$ $n$ = 30 (5); $THOC6^{E188K/E188K}$ $n$ = 18 (5); $THOC6^{W100*/+}$ $n$ = 37 (10); $THOC6^{W100*/W100*}$ $n$ = 16 (5). (C) Immunostaining of EDU, KI67, DCX to assess timing of differentiation in day 28 organoids with quantifications by genotype (D). NR (organoid) number analyzed across three differentiation replicates per genotype: $THOC6^{E188K/+}$ $n$ = 100 (34); $THOC6^{E188K/E188K}$ $n$ = 68 (25); $THOC6^{W100*/+}$ $n$ = 87 (53); $THOC6^{W100*/W100*}$ $n$ = 89 (42). Significance, two-tailed unpaired $t$ test. Data shown as mean ±SEM.
