## Supplemental Tables for "Mechanisms of mRNA processing defects in inherited *THOC6* intellectual disability syndrome"

**Table S1. Clinical descriptions of individuals with TIDS in present study.** *Related to Figure 1.*

| Proband | Clinical description | Genetic diagnosis | Variant Interpretation[a] |
|---|---|---|---|
| 1:II:1 | Proband 1 is from the Netherlands and of European ancestry. He was twelve months old at provided evaluation. The pregnancy was complicated by intrauterine growth restriction at thirty weeks of gestation. Proband 1 was born via a spontaneous vaginal delivery after 38 weeks and 3 days of gestation with a birth weight was 2.252 kg, and a length of 47 cm. Feeding problems were noted after birth. Gastroesophageal reflux disease was diagnosed requiring nasogastric tube feeding intervention. Growth parameters at four months of age included a length of 58 cm (<3rd centile), with a weight of 4.275 kg (<3rd centile) and an occipitofrontal circumference (OFC) of 38 cm (<3rd centile). Dysmorphic features include plagiocephaly, upslanting/narrow palpebral fissures, a straight nose with a broad nasal base, a smooth philtrum with a thin upper lip, and protruding ears with large, upturned earlobes (Figure 1B). A renal ultrasound revealed unilateral renal agenesis, though cardiac abnormalities were not detected by ultrasound evaluation. Marked signs of developmental delay at 12 months. Motor delays characterized by deficits in coordination required for prolonged prone positioning on his abdomen and inability to sit unassisted. | Karyotyping results were normal. Trio WES (mother/father/proband) was performed and identified two *THOC6* variants as high priority candidates for the clinical presentation observed for Proband 1. A previously described missense variant was maternally inherited and a novel truncating *THOC6* variant was paternally inherited: c.569G>A, (p.G190E) and c.139C>T, (p.Q47*) (Figure 1A). The c.139C>T, (p.Q47*) variant was not found in the gnomAD Browser (v2.1.1) and was submitted to ClinVar (https://www.ncbi.nlm.nih.gov/clinvar/; Submission ID:SUB5265728). The parents of Proband 1 were heterozygous for the identified variants, consistent with a recessive mode of inheritance. | p.G190E: pathogenic (ii);<br><br>p.Q47*: pathogenic (ia) |
| 2:III:2 | Proband 2 is the second-born of consanguineous parents of Turkish ancestry, three years old at provided evaluation (Figure 1A). Proband 2 was born full-term (41 weeks of gestation) with a birth weight of 2.7 kg (10th centile) and an OFC of 32.5 cm (<3rd centile). She exhibited global developmental delay, feeding problems, and persistent vomiting in the first few postnatal months, consistent with features of gastroesophageal reflux disease. Use of a nasoduodenal feeding tube supported slow, yet persistent weight gain. Physical examination revealed microcephaly and atypical facial features of epicanthus, long nose with low hanging columella, and upslanting palpebral fissures (Figure 1B). Cranial MRI revealed corpus callosum hypoplasia (Figure 1C). Frequent upper respiratory infections occurred one to three times per month. Echocardiogram demonstrated peripheral pulmonary stenosis and patent foramen ovale. Renal anatomy was unremarkable. At 3 years of age, weight was 13 kg (3rd centile), height was 95 cm (3rd centile) and OFC was 45 cm (<3rd centile). Severe intellectual disability was noted with delays in speech and communication requiring special rehabilitation intervention. Motor delays were observed. At time of evaluation, Proband 2 walked with an unsteady gait. While hearing and vision are normal, strabismus was observed. She has a high number of dental caries, in line with previously described BBIS dental anomalies. | Karyotyping results were normal. WES genetic testing identified a novel biallelic truncating variant c.299G>A, (p.W100*) in exon 4 of *THOC6* in Proband 2. At the time of identification, this variant was not reported and not observed in the gnomAD Browser (v2.1.1) and was submitted to ClinVar (https://www.ncbi.nlm.nih.gov/clinvar/; Submission ID: SUB5265724). Heterozygosity of the *THOC6* variant was confirmed by Sanger sequencing in the mother, as well as in an unaffected sibling. | p.W100*: pathogenic (ia) |
| 3:III:1 | Proband 3 is from the United States of European ancestry, eleven years old at provided evaluation. He was evaluated for multiple congenital anomalies during the newborn period. Microcephaly, deep set eyes, mild epicanthal folds, upslanting palpebral fissures were described during examination in infancy; prominent antihelices of the ears, broad nasal bridge, mild depression of the right nostril with evidence of right cleft lip repair, nasal columella extending below nares, and short philtrum were noted (Figure 1B). Clinical history at age eleven was significant for cleft lip, bifid uvula, ankyloglossia, horseshoe kidney, imperforate anus, developmental delay, autism, and partial complex seizures (onset at 11 years). Developmentally, receptive language skills were noted to be better than expressive language that was restricted to sign language, some words, and gesturing. | Karyotype and FISH analysis for 22q11.2 were normal (Oxford Gene Technology Syndrome). Chromosome microarray (Affymetrix CytoScan Dx) identified a maternally inherited deletion on 5q21.1, the inheritance of which is inconsistent with genetic basis of BBIS clinical features. WES analysis of Proband 3 identified three previously described homozygous missense variants in *cis* in *THOC6* c.[298T>A;700G>C;824G>A], (p.[W100R;V234L;G275D]). This haplotype was previously reported in three other individuals with BBIS from | p.W100R, p.V234L, & p.G275D: pathogenic (ii) |

| | | | |
|---|---|---|---|
| | Assistive technology was used to supplement this deficit. Reading and math skills at age eleven were measured to be equivalent to first grade level. On physical examination, both OFC and weight were less than the 2nd centile, and height was at the 5th centile. | unrelated families (6, 20). In all cases, no consanguinity was reported. The parents of Proband 3 are heterozygous for the identified variants, indicating biparental inheritance of the *THOC6* variants. | |
| 4:IV:1 | Proband 4 was evaluated at thirteen years of age and is the first-born of third-degree consanguineous parents from Southern India (Figure 1A). She weighed 2.75 kg at birth and had complaints of repeated lower respiratory tract infections since the newborn period. Developmental delay, predominantly cognitive, was noted. She crawled at nine months, sat at one year and walked at two years. At present, she says bisyllables only. She exhibited friendly behavior. Examination revealed an OFC of 44.5 cm (<3rd centile, -7 SD) and a height of 122 cm (normal). Facial dysmorphism included upslanting palpebral fissures, epicanthal folds, microcornea, long nose with overhanging columella, thick vermillion borders, and crowded teeth (Figure 1B). Upon neurological examination, she had contractures at the ankle, normal to increased tone, and normal deep tendon reflexes. Vision and hearing were normal. Brain imaging results were also normal. | [see below] | p.G275D is likely pathogenic (ii) |
| 4:IV:2 | Individual 5 (4:IV:2) is the affected sibling of Proband 4 (Figure 1A). She was eight years old upon examination. She presented with global developmental delay like her elder sibling. She achieved head control at five months of age, sat with support at one year, and stood with and without support at one and two years, respectively. Presently, she follows simple commands. Parents also reported nocturnal enuresis. Her OFC was 44.5 cm (<3rd centile, -6 SD). She was 104 cm tall and weighed 12 kg. Epicanthal folds, microretrognathia, cleft palate, and U-shaped uvula were noted upon examination (Figure 1B). Vision and hearing were normal. | Karyotyping results were normal for both siblings. The biallelic missense variant c.824G>A, (p.G275D) in exon 12 of *THOC6* was identified in both Proband 4 and Individual 5 using WES testing. Variant p.G275D has previously been observed in individuals with BBIS and its submission accession is SCV000741884.1. The parents were heterozygous for the identified variants, as confirmed by Sanger sequencing and consistent with the mode of inheritance (Figure 1A). | p.G275D is likely pathogenic (ii) |
| 5:II:3 | Proband 5 was born to consanguineous parents of Moroccan ancestry. She has a healthy dizygotic twin sister as well as a healthy older sister and younger brother. She was born at 37 weeks of gestation with birth parameters of -2 SD (birth weight of 2.08 kg, birth length of 44 cm, and OFC of 31 cm). She presented short-segment Hirschsprung disease, submucous cleft palate, and unilateral choanal stenosis that was surgically repaired at 18 months. She has delayed psychomotor development and overall growth (parameters of -2 SD). She is shy with nasal speech limited to short sentences. At 10 years of age, she has a long narrow face, arched eyebrows, convergent strabismus, a tubular nose with a high nasal bridge, short columella, cupid bow-shaped mouth, and normal ears. Cutaneous 2-3 syndactyly on her feet and clinodactyly of the 5th fingers were also noted. No ophthalmologic abnormalities were present except for convergent strabismus. Auditory evoked potential (AEP), brain and temporal bones CT-scan, and cardiac ultrasound were normal. | 800-bands resolution karyotype showed normal chromosomes on lymphocytes, 46XX, with no 22q11.2 deletion by FISH anailysis at the TUPPLE1 locus. WES analysis identified biallelic variants of c.740G>A, (p.R247Q). Variant was absent in unaffected siblings whereas unaffected parents were heterozygous for the variant. Results confirmed by Sanger sequencing and consistent with recessive mode of inheritance. | p.R247Q is likely pathogenic (ii) |
| 6:IV:1 | Proband 6 was 13 months old at last examination. He had an OFC of 42.5 cm (-3/4 SD) that is likely progressive. Proband 6 has dysmorphic facial features. Hypotonicity, no spasticity, and absent tendon reflexes were noted upon neurological examination. He has hypoplastic genitalia. MRI revealed atrophy of cortex and cerebellum as well as ventriculomegaly and a thin corpous callosum. Proband 6 had intermediate delayed psychomotor development. | WES analysis identified biallelic variants of c.562G>A, (p.E188K). Variant was absent in unaffected siblings whereas unaffected parents were heterozygous for the variant. Results confirmed by Sanger sequencing and consistent with recessive mode of inheritance. | p.E188K is likely pathogenic (ii) |

| | | | |
|---|---|---|---|
| 7:V:1 | Proband 7 had an OFC of -5 SD. He has epilepsy and micropenis. | WES analysis identified biallelic variants of c.299G>A, (p.W100*). Variant was absent in unaffected siblings whereas unaffected parents were heterozygous for the variant. Results confirmed by Sanger sequencing and consistent with recessive mode of inheritance. | p.W100*: pathogenic (ia) |

[a]According to American College of Medical Genetics and Genomics (ACMG) guidelines

**Table S2: Clinical summary of all reported individuals with TIDS.** Absence of features may be due to lack of reporting. *Related to Figure 1.*

| Clinical feature | Prevalence | |
|---|---|---|
| | current study | all published (excluding prenatal report) |
| Intellectual disability | 8 / 8 (100%) | 33 / 33 (100%) |
| Facial dysmorphisms | 8 / 8 (100%) | 31 / 33 (93.9%) |
| Microcephaly | 8 / 8 (100%) | 27 / 33 (81.8%) |
| Teeth anomalies | 2 / 8 (25%) | 15 / 33 (45.5 %) |
| Short stature | 3 / 8 (37.5%) | 13 / 33 (39.4%) |
| Cardiac defects | 1 / 8 (12.5%) | 12 / 33 (36.4%) |
| Renal malformations | 3 / 8 (37.5%) | 11 / 33 (33.3%) |
| Genitourinary issues | 4 / 8 (50%) | 21 / 33 (63.6%) |
| Feeding difficulties | 2 / 8 (25%) | 6 / 33 (18.2%) |
| Ventriculomegaly | 1 / 8 (12.5%) | 7 / 33 (21.2%) |
| ASD or autistic features | 1 / 8 (12.5%) | 8 / 33 (24.2%) |

**Table S3: PCR, RT-PCR, and qRT-PCR primer sequences for human and mouse samples.** *Related to Figures 1-6.*

| Primer type | **ID** | **forward sequence** | **reverse sequence** | **Species** |
|---|---|---|---|---|
| *PCR* | THOC6 genotyping | 5' CTTCAGGACTTTGGGTGGGA 3' | 5' AAAGAACTGTGAGTGGTGCC 3' | human |
| | Thoc6 genotyping | 5' ACGAGAAGAGCCACCATCAG 3' | 5' ATCACTTTTCGTGGGCTCAG 3' | mouse |
| *RT-PCR* | ABCA1 SE | 5' CATTCAGATGCACGTCTGCT 3' | 5' GCTGGCTGTCAAAGAGGAAC 3' | human |
| | POU2F2 SE | 5' TTCTGCATGTCCTTCACTGC 3' | 5' CTTGCTCCAATTCCTGCTGT 3' | human |
| | MAPK15 RI | 5' CCAGACAGCAGAAACCCTGT 3' | 5' TTGGGGGTCTCTGACATAGG 3' | human |
| | Thoc6 splice site | 5' GTTCCCATCGTTCGTACTGG 3' | 5' GCTTTGAAGGAAGCCAACTG 3' | mouse |
| | Admts6 RI | 5' GTTCCCATCGTTCGTACTGG 3' | 5' GCTTTGAAGGAAGCCAACTG 3' | mouse |
| | Fam214b RI | 5' CAGGCTTGAGGCTTCTTCAT 3' | 5' GGTTCTTCAGGGGGTAGTCC 3' | mouse |
| | Cenpt RI | 5' TCCTCCAGCACACATGACTC 3' | 5' CTGGATGGGCACTTAGCTGT 3' | mouse |
| *qRT-PCR* | GAPDH qPCR | 5' TCTTTTGCGTCGCCAGCCGA 3' | 5' ACCAGGCGCCCAATACGACC 3' | human |
| | FOS qPCR | 5' GTGGGAATGAAGTTGGCACT 3' | 5' CTACCACTCACCCGCAGACT 3' | human |
| | THOC6 qPCR | 5' CGACTGGATGGTCTGTGGAG 3' | 5' CCTGGTAGAAGGTGACGTGC 3' | human |
| | MEG3 qPCR | 5' GCATTAAGCCCTGACCTTTG 3' | 5' TCCAGTTTGCTAGCAGGTGA 3' | human |
| | MEG8 qPCR | 5' CCTCAGTATCCTGCGAGCTG 3' | 5' AAGTCAGACCCAGGCAACAC 3' | human |
| | ESRG qPCR | 5' CAGCCTTGTACCCTGGTCTT 3' | 5' ATGCATTGGCTTGTTGCTGA 3' | human |
| | NEAT1 qPCR | 5' GGCAGGTCTAGTTTGGGCAT 3' | 5' CCTCATCCCTCCCAGTACCA 3' | human |
| | TGFB2 qPCR | 5' AAGAAGCGTGCTTTGGATGCGG 3' | 5' ATGCTCCAGCACAGAAGTTGGC 3' | human |
| | ID4 qPCR | 5' GGACCTGTCCAGCCGCGCC 3' | 5' TCAGCGGCACAGAATGCTGTCG 3' | human |
| | TP53 qPCR | 5' CCTCAGCATCTTATCCGAGTGG 3' | 5' TGGATGGTGGTACAGTCAGAGC 3' | human |
| | PAX6 qPCR | 5' CTGAGGAATCAGAGAAGACAGGC 3' | 5' ATGGAGCCAGATGTGAAGGAGG 3' | human |
| | CXXC4 qPCR | 5' TGCCCGCAGAATCATTCCTCCT 3' | 5' ACGCCACAGTTGATGAGCCTCT 3' | human |
| | WNT7A qPCR | 5' AGGAGAAGGCTCACAAATGGGC 3' | 5' CGGCAATGATGGCGTAGGTGAA 3' | human |
| | Ier3 qPCR | 5' CCATCTCCACACCATGACTG 3' | 5' CTCCGAGGTCAGGTTCAAAG 3' | mouse |
| | Islr2 qPCR | 5' CTGCAAGTCAGAGAGCAGCA 3' | 5' AACTGGTGGGCGTACTTGTC 3' | mouse |
| | Thoc6 qPCR | 5' GCAACAATTACGGGCAGATT 3' | 5' CAACCCTTGACCTCTCCATC 3' | mouse |
| | Anxa2 qPCR | 5' CATTCTACACCCCCAAGTGC 3' | 5' CTGATAGGCGAAGGCAATGT 3' | mouse |
| | Vegfa qPCR | 5' GGTTCCAGAAGGGAGAGGAG 3' | 5' GGCAGTAGCTTCGCTGGTAG 3' | mouse |
| | Kcnt2 qPCR | 5' AAGGCTGGCAAAATGATGAC 3' | 5' CTGTGACGGTTTCTCAAGCA 3' | mouse |
| | Wnt7a qPCR | 5' GGTGCGAGCATCATCTGTAA 3' | 5' TGGTACTGGCCTTGCTTCTC 3' | mouse |
| *sgRNA* | *Thoc6$^{fs/fs}$* sgRNA | 5' CACCGCACCGCTCGCGGTGCCTCT 3' | 5' CAGAGGCACCGCGAGCGGTGCCAAA 3' | mouse |
